# Improved chromosome level genome assembly of the Glanville fritillary butterfly (*Melitaea cinxia*) based on SMRT Sequencing and linkage map

**DOI:** 10.1101/2020.11.03.364950

**Authors:** Daniel Blande, Olli-Pekka Smolander, Virpi Ahola, Pasi Rastas, Jaakko Tanskanen, Juhana I. Kammonen, Vicencio Oostra, Lorenzo Pellegrini, Suvi Ikonen, Tad Dallas, Michelle F. DiLeo, Anne Duplouy, Ilhan Cem Duru, Pauliina Halimaa, Aapo Kahilainen, Suyog S. Kuwar, Sirpa O. Kärenlampi, Elvira Lafuente, Shiqi Luo, Jenny Makkonen, Abhilash Nair, Maria de la Paz Celorio-Mancera, Ville Pennanen, Annukka Ruokolainen, Tarja Sundell, Arja I. Tervahauta, Victoria Twort, Erik van Bergen, Janina Österman-Udd, Lars Paulin, Mikko J. Frilander, Petri Auvinen, Marjo Saastamoinen

## Abstract

The Glanville fritillary (*Melitaea cinxia*) butterfly is a long-term model system for metapopulation dynamics research in fragmented landscapes. Here, we provide a chromosome level assembly of the butterfly’s genome produced from Pacific Biosciences sequencing of a pool of males, combined with a linkage map from population crosses. The final assembly size of 484 Mb is an increase of 94 Mb on the previously published genome. Estimation of the completeness of the genome with BUSCO, indicates that the genome contains 93 - 95% of the BUSCO genes in complete and single copies. We predicted 14,830 gene models using the MAKER pipeline and manually curated 1,232 of these gene models. The genome and its annotated gene models are a valuable resource for future comparative genomics, molecular biology, transcriptome and genetics studies on this species.

## Data Description

### Context

Identifying and characterizing genes underlying ecologically and evolutionarily relevant phenotypes in natural populations has become possible with novel genomic tools that can also be utilized in ‘non-model’ organisms. The Glanville fritillary (*Melitaea cinxia*) butterfly, and in particular its metapopulation in the Åland Islands (SW Finland), is an ecological model system in spatial ecology[1,2]. In short, within this archipelago, the species inhabits a network of dry outcrop meadows and pastures, and persists as a classic metapopulation with high turnover in patch occupancy[1]. The network of 4,500 potential habitat patches has been systematically surveyed bi-annually for butterfly occupancy and abundance since 1993[3], providing a vast amount of ecological data on population dynamics[2]. Experimental manipulations under more controlled conditions are also possible due to the small size, high fecundity and relatively short generation time of the species. Consequently, the ecological understanding of the species expands to the life history ecology across development stages[4] (e.g. Kahilainen et al. unpubl.), dispersal dynamics[5,6], species interactions (host plants and parasitoids)[7–11], and stress tolerance[12,13]. During the last decade, the system has also been used to study genetic and evolutionary processes, such as identifying candidate genes underlying variation and evolution of dispersal in fragmented habitats[14] and host plant preference[15], and assessing allelic variation and their dynamics in space and time with SNP data across populations[16–18]. Several approaches have been used to explore the genetic underpinnings of phenotypic variation in the Glanville fritillary metapopulation, ranging from candidate gene approaches[12,19] to quantitative genetics[20,21], and whole-genome scans[22,23] (including transcriptomics; Kahilainen et al. unpubl.), and included both laboratory and natural environmental conditions.

The first *M. cinxia* genome was released in 2014[24]. This genome was produced from a combination of 454 sequencing for contig assembly, followed by scaffolding with Illumina paired-end (PE), SOLiD mate-pair reads and PacBio data. The size of the final assembly was 390 Mb made up from 8,261 scaffolds, with a scaffold N50 of 119,328. Scaffolds were assigned to chromosomes based on a linkage map produced from RAD sequencing[25]. It was considered that a new genome, sequenced using longer PacBio reads, would result in a more complete assembly and better represent the repetitive areas of the genome.

Here, a new sequencing and assembly of the *M. cinxia* genome has been carried out using a pool of seven male butterflies from a single larval family collected from Sottunga, an island in an eastern part of the archipelago. Sequencing was conducted using the PacBio RSII sequencer. An initial assembly was created using FALCON[26,27] followed by polishing performed with Quiver[26]. A new linkage map was created and used to assign the assembled scaffolds to their correct positions and orientations within the 31 chromosomes. The scaffolds were then gap-filled producing a final assembly of 484 Mb with a scaffold N50 of 17,331,753 bp. Gene prediction on the genome assembly was carried out using MAKER v 2.31.10[28] that was run iteratively using several independent training sets. Manual annotation was performed for 1,232 of the gene models. The genome assembly increases greatly in contiguity and completeness compared to the first genome (Table 1) with superscaffold N50 values of 17,331,753 bp in the new genome compared to 9,578,794 bp in the version 1 genome.

**Table 1.** Assembly statistics were calculated for the *M. cinxia* v2 genome, *M. cinxia* v1 scaffolds, *M. cinxia* v1 chromosomes and *B. mori* using the assembly-stats program (https://zenodo.org/badge/latestdoi/20772/rjchallis/assembly-stats). Statistics for *H. melpomene* v2.5 and *P. napi* v1.1 were obtained from LepBase[64].

|  | *M. cinxia*<br>Version 2 | *M. cinxia*<br>Version 1<br>Scaffolds | *M. cinxia*<br>Version 1<br>Chromo-<br>somes | *Heliconius*<br>*melpomene*<br>Hmel2.5 | *Bombyx*<br>*mori* | *Pieris napi*<br>v1.1 |
| --- | --- | --- | --- | --- | --- | --- |
| Length (bp) | 484,462,241 | 389,907,520 | 250,926,848 | 275,245,813 | 460,334,017 | 349,759,982 |
| N(%) | 0.00 | 7.42 | 7.84 | 0.38 | 0.10 | 22.47 |
| GC(%) | 33.83 | 32.56 | 32.38 | 32.79 | 38.54 | 33.32 |
| AT(%) | 66.17 | 67.44 | 67.62 | 67.21 | 61.46 | 66.68 |
| Scaffold count | 31 | 8,261 | 31 | 332 | 696 | 2,969 |
| Longest scaffold<br>(bp) | 22,190,643 | 668,473 | 13,295,773 | 18,127,314 | 21,465,692 | 15,427,984 |
| Scaffold N50<br>length (bp) | 17,331,753 | 119,328 | 9,578,794 | 14,308,859 | 16,796.068 | 12,597,868 |
| Scaffold N50<br>count (L50) | 13 | 970 | 12 | 9 | 13 | 13 |
| Contig Count | 529 | 48,180 | 26,662 | 3,126 | 726 | 53,510 |
| Contig N50<br>length (bp) | 1,831,849 | 14,057 | 15,707 | 328,886 | 12,201,325 | 10,538 |
| Contig N50<br>count (L50) | 79 | 7,366 | 4,286 | 214 | 16 | 6,914 |

### Methods

An overview of the processing pipeline for the work is shown in Figure 1.

**Figure 1.**
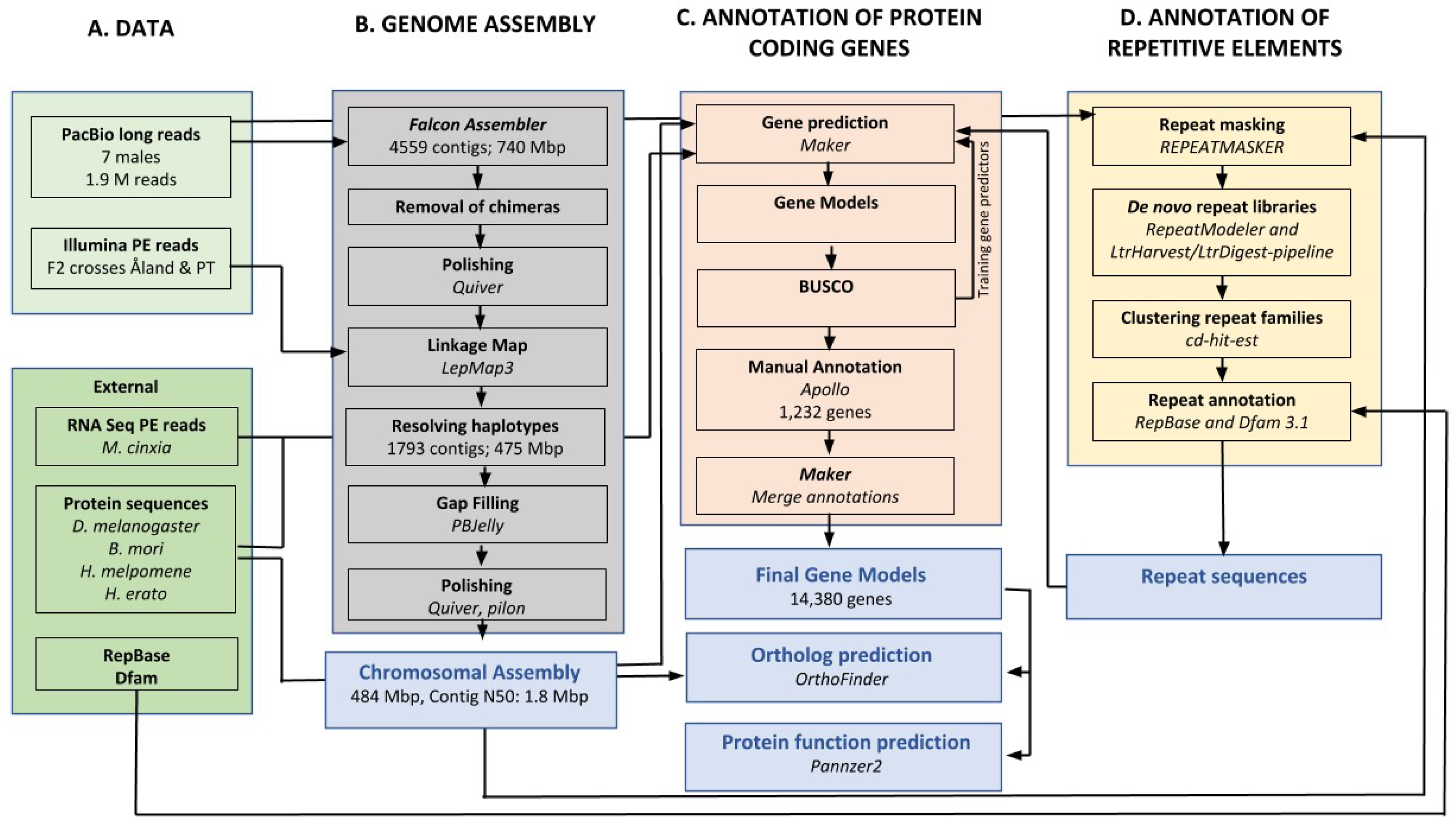
An overview of the assembly and annotation process of the improved Glanville fritillary genome.

#### Genomic samples and DNA extraction

Owing to the facultatively univoltine life cycle of the butterfly in Finland, experimental inbreeding of the species would have taken several years. Therefore, we chose to sample individuals from an island population, Sottunga, expected to harbour low genetic diversity. Sottunga is part of the Åland Islands archipelago in the northern Baltic Sea, and the population was introduced here in 1991 using individuals collected on the mainland of Åland Island[29]. This introduction was carried out with 71 larval families. The distance to the nearest *M. cinxia* population across the water is 5 km, and we therefore assume that the introduced population has been (almost) completely isolated ever since. Furthermore, the effective population size of *M. cinxia* in Sottunga has been very low during the last 24 years (on average 57 larval nests/year in 1993-2019), and it has experienced several strong bottlenecks[30]. Using genomic markers, Fountain et al.[16] demonstrated that samples from the Sottunga population separate clearly from samples collected on the mainland.

During the fall survey of 2014 (see Ojanen et al. for details of the survey[3]) we collected individuals from one larval group on the island of Sottunga (patch number 1439, see Schulz et al. for specific location[31]). The larvae were collected once they were in diapause and most likely comprise full-sibs[17]. The larval group was kept in diapause (+5 °C) until the following spring and then reared to adulthood under common garden conditions (28:8°C; 12L:12D) at the Lammi Biological Station, University of Helsinki. After eclosion, butterflies were sexed and stored at −80°C. High-molecular-weight DNA was isolated from seven adult males using the caesium chloride (CsCl) method[24].

#### SMRT sequencing libraries and sequencing

Library construction for Pacific Biosciences sequencing was carried out using the protocols recommended by the manufacturer (Pacific Biosciences, Menlo Park, CA, USA). Genomic DNA was sheared using a Megaruptor (Diagenode, Seraing, Belgium) followed by damage repair, end-repair, hairpin ligation, and size selection using BluePippin (Sage Science, Beverly, MA, USA). After primer annealing and polymerase binding, the DNA templates were sequenced on a PacBio RSII sequencer using P6/C4 chemistry and 360 min video time at the DNA Sequencing and Genomics Laboratory, Institute of Biotechnology, University of Helsinki, Finland[32].

#### Genome Assembly

The genome was assembled using the FALCON assembler (FALCON-Integrate-1.8.6)[26,27] with a read cut-off of 18,000. This cut-off was experimentally found to give the best contiguity for the assembly, while minimizing (within a small margin of error) the percentage of possibly erroneous contigs. The assembly errors were evaluated based on a genetic map from the previously published genome[24]. The assembly was based on 1,939,889 PacBio reads, 24,409,505,551 bp in total, with an N50 of 18,479 bp which is approximately 50x coverage based on the final genome size. With the selected read cut-off the data produced 10.8 Gb of corrected reads which were further assembled using the Falcon software. The assembly yielded 4,559 primary contigs containing 739.9 Mb with an N50 of 340 kb and 1,661 alternative contigs containing 118.1 Mb with an N50 of 85,246 bp. The data were also assembled using miniasm software (0.2-r137-dirty)[33] which yielded similar results.

To evaluate the putative chimeric contigs and assembly errors suggested by the genetic map, the raw SMRT sequencing data were mapped to the assembly using the Burrows-Wheeler Aligner (BWA-0.7.17) with the MEM algorithm[34]. The alignments of the 425 regions discovered as possibly chimeric were visually inspected. Of these regions, 92 showed even read coverage and no evident signs of assembly errors, while 333 regions contained areas with low coverage and/or repeat regions that led to erroneous overlaps and misassemblies. The assembly was split in the positions where the error was observed. The resulting assembly was polished using the SMRT sequencing data and Quiver[26] software from the SMRT Tools-package (PacBio).

#### Linkage Map

Linkage mapping was constructed from whole genome resequencing data of F2 crosses of *M. cinxia* (Ahola et al., unpublished). The grandparents of these F2 crosses are offspring of wild collected *M. cinxia* originating from two distantly related *M. cinxia* populations around the Baltic Sea; the Åland Islands (ÅL)[1] and Pieni Tytärsaari (PT) populations[35]. Between population crosses of type ÅL♂xPT♀ and ÅL♀xPT♂ were established to create the F1 population. Part of these F1 individuals were used to establish the F2 families, actively avoiding mating among siblings. A subset of the resulting full-sib families were reared to adulthood, and five of these F2 families, together with their parents and grandparents, were selected for resequencing. In total, resequencing included ten grandparental individuals, ten F1 parents and 165 F2 individuals (N=185).

All the larvae from different generations completed development under common garden conditions (28:15°C; 12L:12D) utilizing fresh leaves of greenhouse grown *Veronica spicata*. Diapausing larvae were kept in a growth chamber at +5°C and 80% RH for approximately seven months to mimic the normal wintertime conditions for these butterflies. Adults were kept in hanging cages (of 50 cm height and 40 cm diameter) at ~26:18°C; 9L:15, and fed *ad libitum* with 20% honey-water solution throughout the experiments.

Before DNA extraction the adult butterflies were stored at −80°C, and either thorax or abdomen tissue of these individuals was used for sequencing. Tissues were homogenized prior to extraction using TissueLyser (Qiagen, Venlo, The Netherlands) at 30/s for 1.5 mins with Tungsten Carbide Beads, 3 mm (Qiagen, Venlo, The Netherlands) and ATL buffer (Qiagen, Venlo, The Netherlands). DNA was extracted using the NucleoSpin 96 Tissue Core Kit (Macherey-Nagel) according to the manufacturer’s protocol with the exception that lysing time was extended to overnight. The samples were additionally treated with RNase A (Thermo Scientific) before sequencing. Sequencing was made using Illumina Hi-Seq with 125 bp paired-end reads.

The mapping procedure followed the Lep-MAP3[25] pipeline. First, individual fastq files were mapped to the contig assembly using BWA MEM (BWA-0.7.17) [34] and individual bam files were created using SAMtools (1.6)[36,37]. SAMtools mpileup and the scripts pileupParser2.awk and pileup2posterior.awk were used to obtain input data for Lep-MAP3. Then ParentCall2 (parameter: ZLimit=2) and Filtering2 (parameters: dataTolerance=0.0001; removeNonInformative=1; familyInformativeLimit=4) were run to obtain data with at least four informative families for each marker, resulting in a final input with almost 2.5M markers.

SeparateChromosomes2 was run on the final data (parameters lodLimit=20; samplePair=0.2;numThreads=48) to obtain 31 linkage groups with a total of 2.4M markers. OrderMarkers2 was run (parameter recombination2=0) on each linkage group (chromosome). This map was used to anchor the contig assembly into chromosomes. To validate anchoring, the map construction was repeated in the same way except that OrderMarkers2 was run on the physical order of markers to reduce noise in the linkage map. Finally, the raw data were re-mapped to the gap-filled chromosome level assembly and the linkage map was re-done in the new physical order to infer final recombination rates.

#### Anchoring the genome and resolving haplotypes using the linkage map

The contigs were aligned against each other and lift-over chains were created by running the first two steps (batch A1 and A2) of HaploMerger2[38]. By inspecting this chain, contigs fully contained in some longer contig were removed. Initial contig order and orientation within each chromosome was calculated by the median map position of each contig and the longest increasing subsequence of markers, respectively. For each chromosome, Marey map and contig-contig alignments from the chain were recorded. The contigs orders and orientations were manually fixed when needed and any assembly errors that were found were corrected by splitting the contigs accordingly. Also, partially haplotypic contigs were found and collapsed based on the Marey maps and contig-contig alignments. This manual work resulted in the final reference genome sequence including start and end positions of contigs in the correct order and orientation for each chromosome. Finally, the haplotype corrected genome was gap-filled using PBJelly software (PBSuite_15.8.24)[39] with the original SMRT sequencing data, and polished with the Quiver tool[26] from the SMRT Tools-package 2.3.0 (PacBio) and with Pilon (1.21)[40].

The chromosomes were aligned against the *Heliconius melpomene* (2.5)[41,42] and *Pieris napi*[43] genomes using the LAST aligner(938)[44] to check structural similarity between the species (Supplementary Figures S1-13). An overview alignment for *H. melpomene* was created using D-GENIES (1.2.0)[45] (Figure 2). The data show a high level of collinearity between *M. cinxia* and *H. melpomene* chromosomes, as described before in Ahola et al.[24]. A notably interesting point is the lack of collinearity with sex chromosomes (*M. cinxia* chromosome 1 & *H. melpomene* chromosome 21). Furthermore, the visible vertical lines show the effect of long read assembly on repeat resolution. In *M. cinxia* the repeats are placed in single chromosomes whereas in *H. melpomene* they are present in all chromosomes.

**Figure 2.**
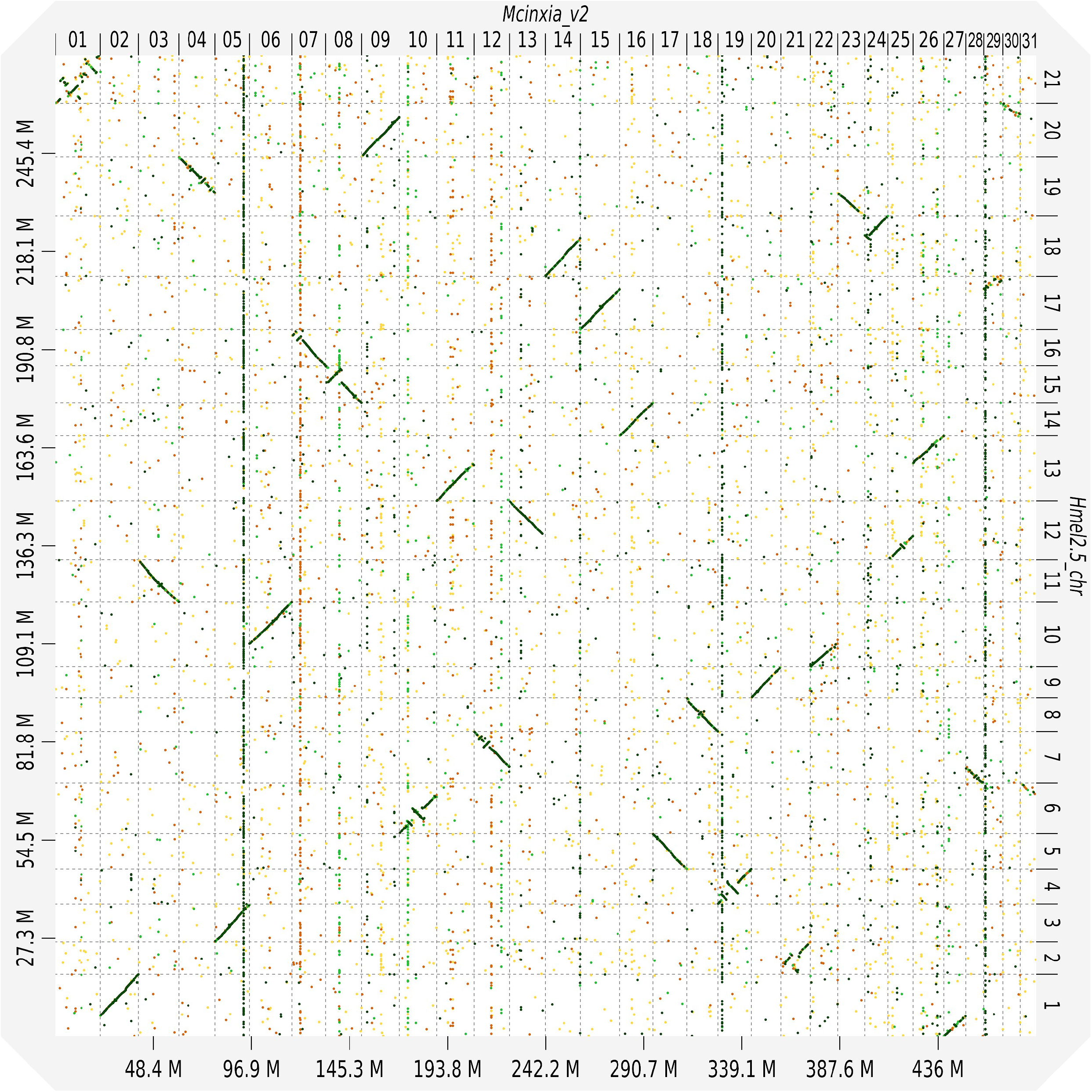
A dot-plot structural comparison of the *H. melpomene* genome against the *M. cinxia* v2 genome. The alignment was created using D-GENIES (1.2.0)[45]. The diagonal lines indicate the collinearity between the two species. The lack of collinearity in sex chromosomes is visible in the upper left corner between Mcnxia_v2 chr 01 and Hmel2.5 chr 21. The visible vertical lines show repeats that are resolved in Mcinxia_v2 but are present in all chromosomes in Hmel2.5_chr.

#### Repeat masking and annotation

Genomic assemblies were masked with *de novo* repeat libraries by RepeatMasker v.4.0.9 (http://www.repeatmasker.org/). *De novo* repeat libraries were constructed from original PacBio reads with lengths over 30,000bp and assembled scaffolds (pseudo chromosomes) using RepeatModeler v 1.0.10 (http://www.repeatmasker.org/RepeatModeler/) and the LtrHarvest/LtrDigest-pipeline[46,47]. Repeat families were clustered using cd-hit-est applying 80/80-rule (80% identity over 80% length)[48]. Repeat annotations were confirmed by RepBase Release 20181026[49] and Dfam version 3.1[50].

#### Transcriptome assembly

In order to aid construction of gene models, we used two transcriptome assemblies, each based on separate RNASeq data sets. Transcriptome 1 was produced using a set of 78 individually sequenced female larvae of an outbred lab population (Kahilainen et al. unpubl. data), sequenced to an average depth of 17.3M reads (read lengths 85 bp and 65 bp for forward and reverse PE reads, respectively). As the two sexes are practically indistinguishable in the larval stages, the females were identified based on homozygosity across a set of 22 Z-chromosome specific SNP loci (Kahilainen et al. unpubl. data). To remove Illumina adapter sequences, we trimmed raw reads using Trimmomatic (Trimmomatic-0.35)[51], and normalised using Trinity v2.6.5[52]. We then used two separate procedures to construct *de novo* transcriptome assemblies, Trinity (v2.6.5) and Velvet / Oases (1.2.10)[53]. Trinity was run with standard settings, whereas Velvet / Oases used a range of nine kmer sizes (21 to 99 bp), producing a separate assembly for each kmer size. We then combined the resulting assemblies, filtered the combined assembly using the EvidentialGene (tr2aacds.pl VERSION 2017.12.21)[54] pipeline, and removed contigs smaller than 200 bp or expressed at a low level (< 1 normalized counts per million), yielding the final assembly. Transcriptome 2 was constructed from a set of 12 adult females (thorax and abdomen, without ovaries) and 48 first instar larvae, as part of a separate gene expression study (Oostra et al. unpubl. data). RNA from these 60 individual samples was sequenced to an average depth of 16.6M reads (86/74 bp PE). The stranded RNA-seq libraries were made using Ovation^®^ Universal RNA-Seq System (Nugen) with custom ribosomal RNA removal. The libraries were paired-end sequenced on a NextSeq 500 using the 150 bp kit (Illumina) at the DNA sequencing and genomics laboratory Institute of Biotechnology University of Helsinki. We trimmed the reads using fastp (v0.20.0)[55], and used the HISAT2 (2.0.4) / StringTie (1.3.5) pipeline[56] to construct a genome-guided transcriptome assembly, mapping the RNAseq reads to the new genome assembly.

#### Gene model Annotation

Initial gene predictions were obtained by running the MAKER v 2.31.10[28] gene prediction programme in an iterative procedure. In the first round of MAKER, transcriptome assembly 1, described above, was provided as evidence, and genes were predicted solely from the aligned transcripts. This resulted in 14,738 gene models. These gene models were then used for training the SNAP (2013-02-16)[57] and AUGUSTUS (3.3.2)[58] gene predictors. A second round of MAKER was run providing the *de novo* transcripts, trained gene prediction models, repeat masking file and protein data from other lepidopteran species. The MAKER settings were adjusted to allow prediction of gene models without requiring a corresponding transcript in the *de novo* transcriptome assembly. Following each round of MAKER gene prediction, the annotation completeness was assessed using BUSCO[59,60].

#### Manual Annotation

Manual annotation was performed for 1,232 genes, using the Apollo collaborative annotation system version 2.1.0[61]. The collaborative annotation environment was set up in Ubuntu Linux 14.04 server with 250 GB RAM and 48 AMD Opteron 6,168 processing cores. This was later upgraded to a cloud server provided by the Finnish IT Center for Science (CSC) and running on Ubuntu Linux 18.04 with 200 GB RAM and 40 Intel Xeon model 85 processing cores. Evidence tracks were produced containing gene predictions from three rounds of MAKER, RNASeq alignments of sequence reads and protein alignments from other species (Table 2). RNASeq alignments comprised a mixed tissue pooled sample, an abdomen pooled sample and six larval samples (from transcriptome 1) selected to represent a diverse range and included, for example, both sexes and different family backgrounds.

**Table 2.**
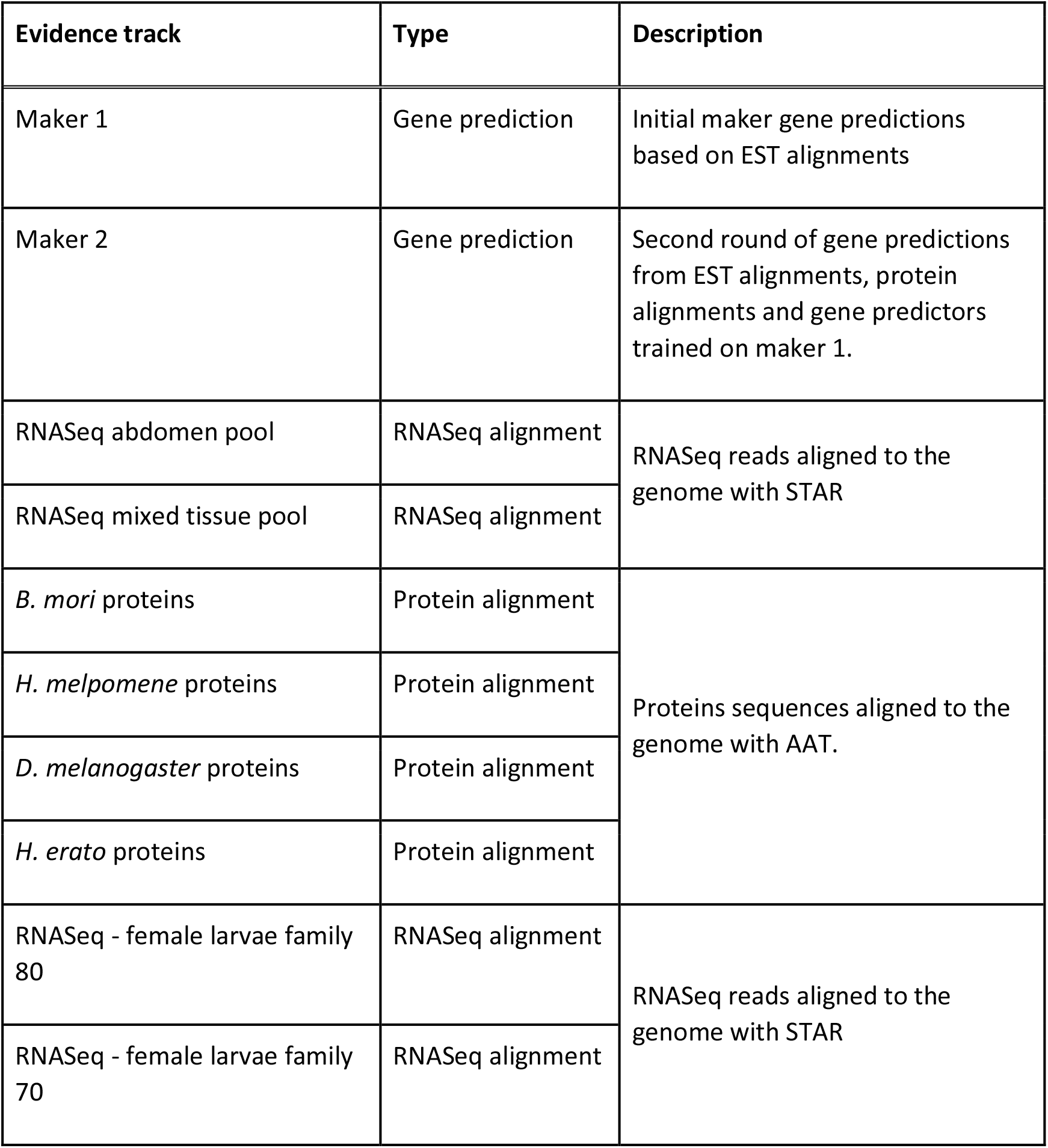
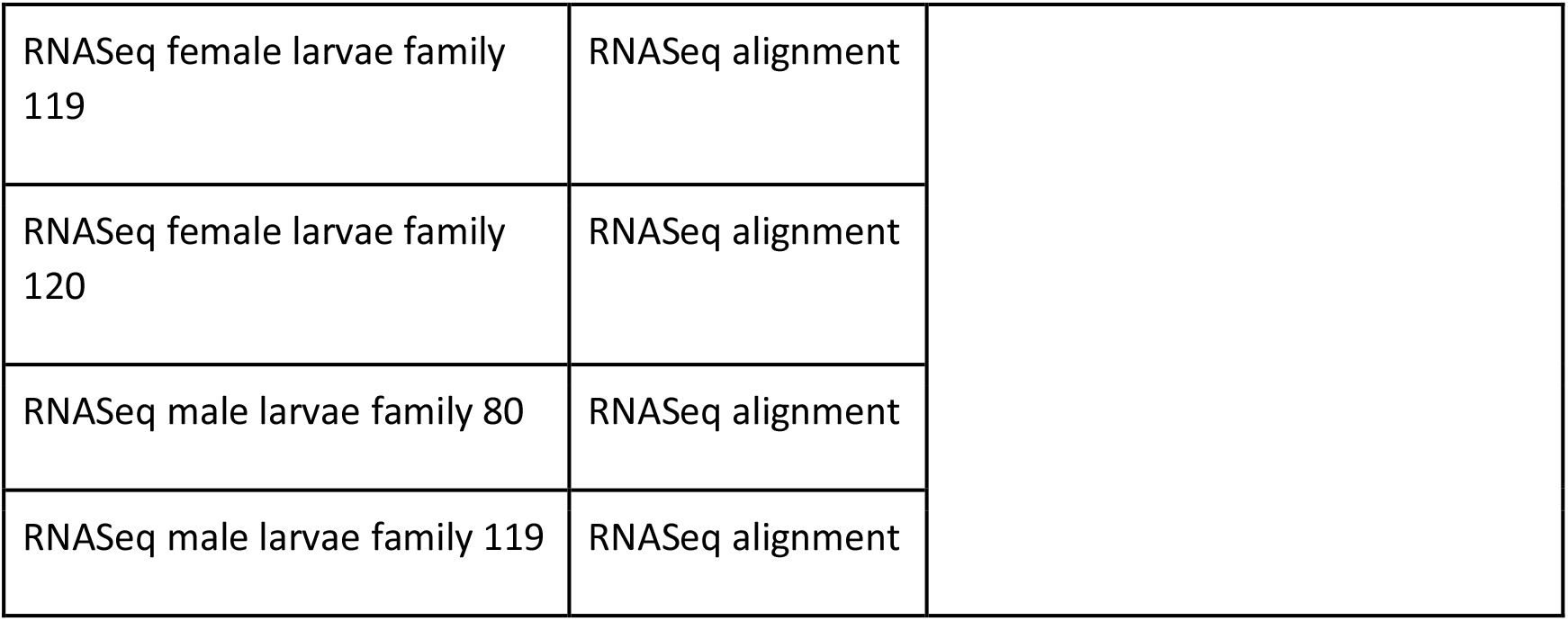
Evidence tracks that were used during the manual annotation of 1,232 M. cinxia genes

#### Final Gene Models

Following the manual annotation, the SNAP[57] and AUGUSTUS[58] gene predictors were retrained using the manually annotated gene models. MAKER was run using the updated gene predictors, transcriptome 1 and 2, and also using a masking file for repeats. As a final step to incorporate the manually annotated gene models, MAKER was run providing the previous MAKER file to pred_gff and the manually annotated models to model_gff. Gene functional prediction was performed using Pannzer v2[62].

#### Ortholog identification

Predicted protein sequences from *Bombyx* mori[63] (January 2017 gene models), *P. napi*[43] and *H. melpomene* (Hmel2.5)[41,42] were downloaded from silkbase http://silkbase.ab.a.u-tokyo.ac.jp/cgi-bin/download.cgi, LepBase[64] and the Butterfly Genome Database http://butterflygenome.org/?q=node/4 respectively. OrthoFinder v2.3.3[65] was run to identify orthologs between *M. cinxia*, *B. mori, P napi* and *H. melpomene* using blast as the search tool (Figure 3 & Supplementary Figure S14).

**Figure 3.**
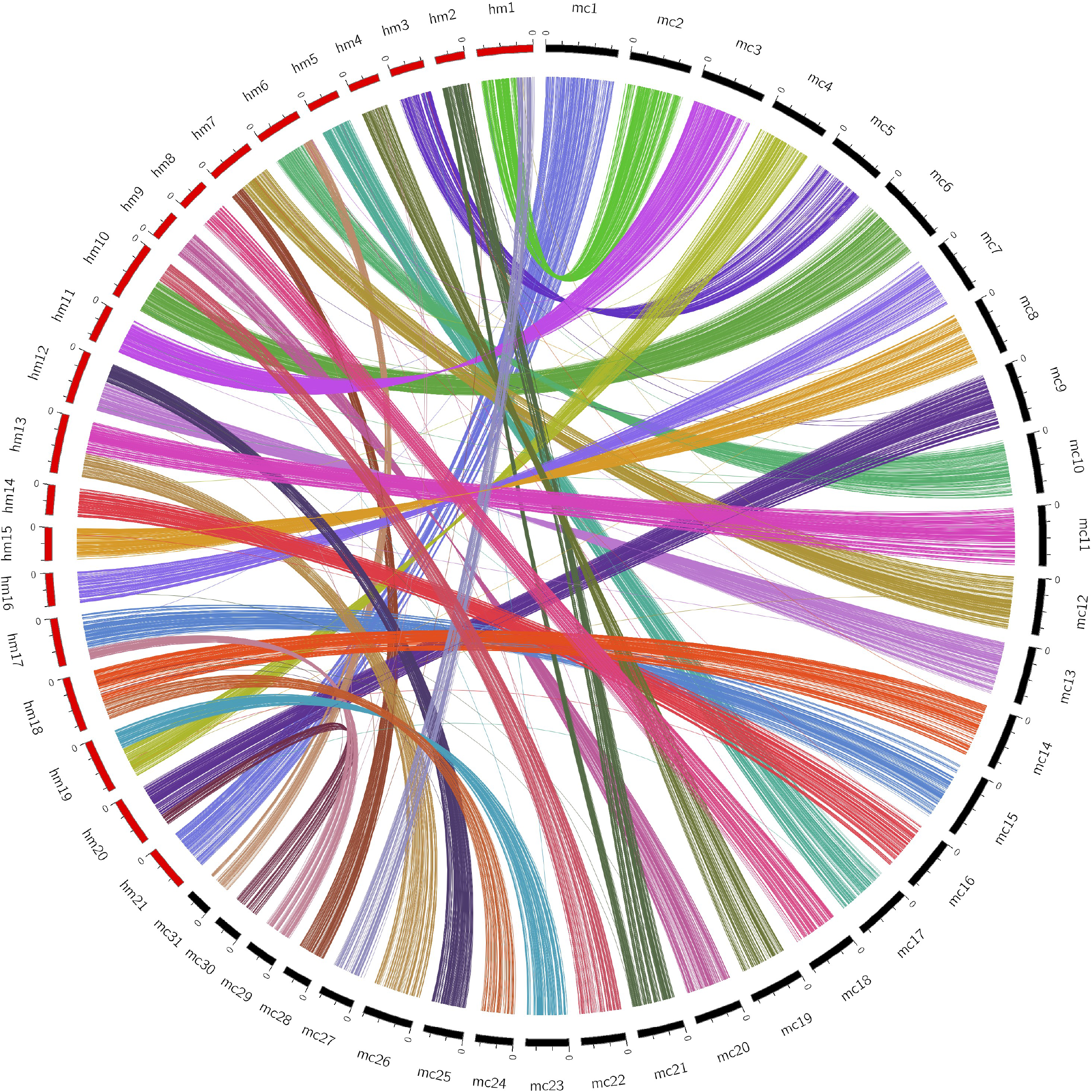
A circos plot showing the orthologs between *M. cinxia* and *H. melpomene* Orthologs between *M. cinxia* and *H. melpomene* were identifies using OrthoFinder and filtered for one-to-one orthologs. The internal links in the circos plot indicate the orthologs between *M. cinxia* and *H. melpomene*. The links are coloured according to the *M. cinxia* chromosome.

### Data Validation and quality control

To assess the quality of the assembly, assembly statistics were generated using assembly-stats[64] and compared to the v1 genome as well as the *H. melpomene*, *B. mori* and *P. napi* genome assemblies (Table 1). The new genome contains 94 Mb more sequence than the previous scaffold assembly. The N50 length and L50 value at scaffold or chromosome level have improved greatly compared to the previous genome. To check for possible duplication or missing areas in the assembly, an assessment was made of the completeness of single copy orthologs from BUSCO[59,60] eukaryota, arthropoda and metazoa gene sets (Table 3). In each of the gene sets, 93.4-94.9% of the expected single copy orthologs were found in complete and single copies. The duplication rate was estimated to be between 1.4 and 2.3%. A total of 1,232 gene models were manually curated using the Apollo annotation system[61] to ensure the quality of the models. To test for contamination, the predicted protein sequences were checked with AAI-profiler[66] to identify sequences originating from different taxa (supplementary files 1-3 (AAI.html, matrix.html, krona.html)). Overall, 42% of the genome was composed of repeat sequences (Figure 4 and Supplementary Figures S15-20 (chromosome specific repeat classes)). There were no clear differences in the repeat contents between chromosomes (Supplementary Table 1). Long interspersed elements (LINE) were the most prevalent.

**Figure 4.**
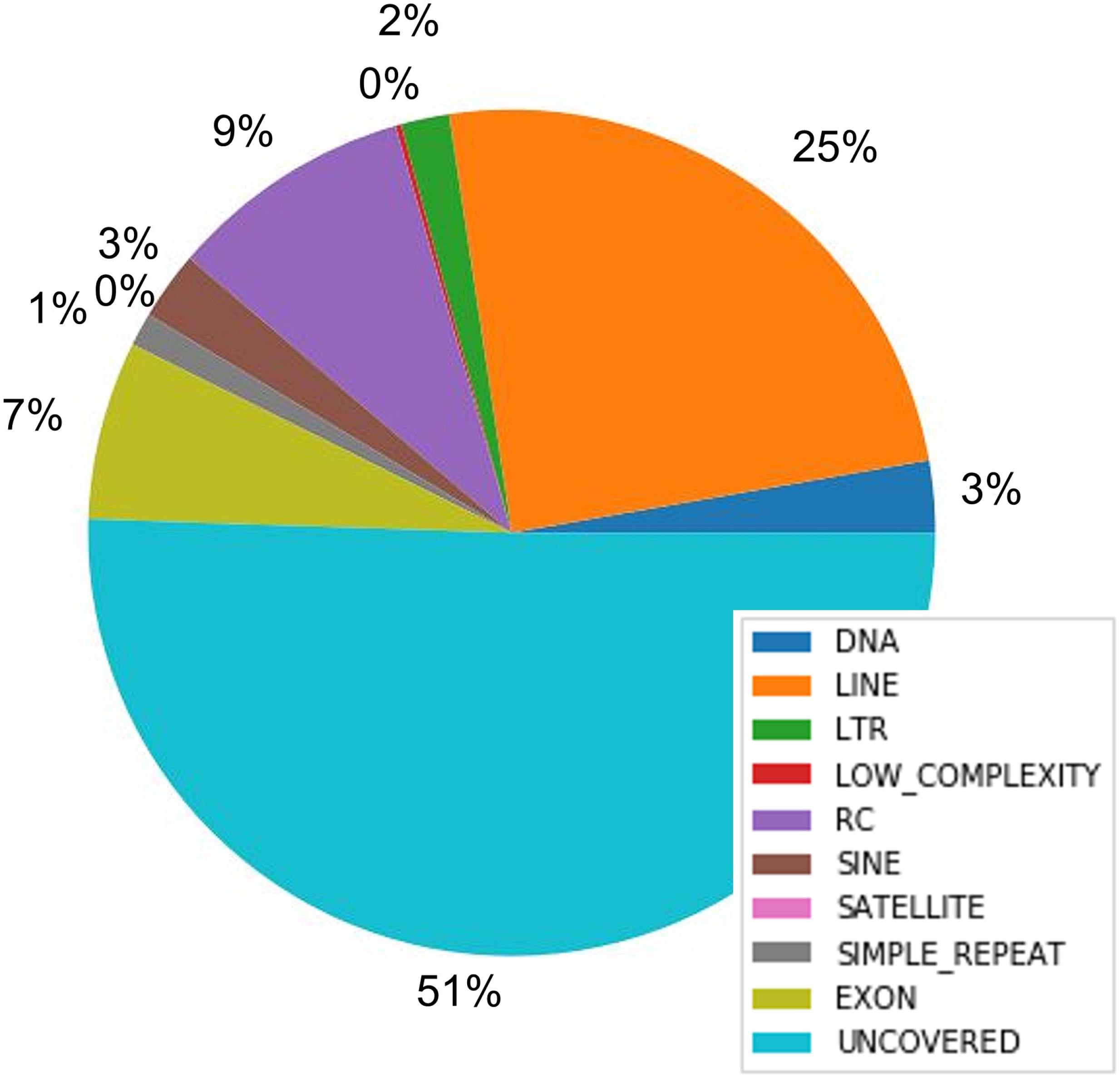
Relative amounts of different repeat classes in *M. cinxia* genome Repeat classes and coverage of the *M. cinxia* genome v2: DNA = classII; LINE = Long interspersed elements; LTR = Long terminal repeats; LOW_COMPLEXITY = Low complexity repeated DNA; RC = Rolling circle elements (e.g. Helitrons); SINE = Short interspersed elements; Satellite = Satellite DNA; SIMPLE_REPEAT = Simple repeated motifs; EXON = Exonic regions; UNCOVERED = rest of the chromosomes.

**Table 3.** BUSCO completeness estimates of the v2 genome based on the eukaryota, arthropoda and metazoa gene sets.

| Lineage | BUSCO Category |  |  |  |  |
| --- | --- | --- | --- | --- | --- |
|  | Complete | Single-copy | Duplicated | Fragmented | Missing |
| Eukaryota | 290 | 283 | 7 | 5 | 8 |
|  | 95.7% | 93.4% | 2.3% | 1.7% | 2.6% |
| Arthropoda | 1027 | 1012 | 15 | 7 | 32 |
|  | 96.3% | 94.9% | 1.4% | 0.7% | 3.0% |
| Metazoa | 935 | 921 | 14 | 12 | 31 |
|  | 95.6% | 94.2% | 1.4% | 1.2% | 3.2% |

### Re-use potential

The substantial improvements in contiguity and gene annotation quality of the new genome will enable a range of important new studies and open up possibilities for future work. Current research aims at identifying mechanisms underlying key life history adaptations, exploring the extent of natural variation and selection on these adaptations in wild populations, and integrating these insights with the exceptional ecological, demographic, and climatic data available for this system. Future studies in this direction will help understand the mechanisms maintaining variation in life-histories across spatial and temporal scales, and on the extent to which phenotypic variation in these and other traits may contribute to a population’s adaptive capacity under climate change. Several studies in different species illustrate how stress responses can be crucial for survival under variable environments, both within and between generations. The Glanville fritillary is being used to explore how environmental information is translated into adaptive phenotypic changes, and how these responses are transmitted to future generations, using transcriptomic and epigenetic approaches. Such studies will greatly benefit from an improved annotation permitting exon-specific expression quantification, and identification of epigenetic marks and other functional variants outside coding regions. Exploiting current and past large-scale sampling efforts, these new studies apply population genomic approaches that are greatly facilitated by the increased assembly contiguity, for instance by permitting linkage disequilibrium (LD) and haplotype-based selection analyses. Other avenues of research enabled by the improved genome assembly include structural variation, regulatory evolution, recombination rate variation, and coalescent-based demographic analyses. The increasing availability of chromosome-level lepidopteran genomes such as ours permits exciting new comparative phylogenetic analyses, for example of chromosome and genome evolution.

### Availability of supporting data

The SMRT sequencing reads used for the genome assembly have been deposited to the sequence read archive.

The genome has been deposited to GenBank.

The Illumina reads used for the linkage map have been deposited to the sequence read archive.

Transcriptome 1 RNASeq reads have been deposited to NCBI GEO.

Transcriptome 2 RNASeq reads have been deposited to NCBI SRA.

## Declarations

### Ethics approval and consent to participate

There are no ethical policies related to working with insect data. The Glanville fritillary is not considered endangered in the Åland islands and no permits are required for sampling. However, we note that within this project the larval sampling for genetic analyses is done non-invasively in the field. ensuring insignificant demographic impact. In addition, as the sampling will take place prior diapause (Åland) when mortality is generally the highest – the collection has negligible effect on the family survival or the demography of populations.

## Supporting information

AAI.html

krona.html

matrix.html

## Competing interests

‘The authors declare that they have no competing interests’.

## Funding

Funding for M.S, D.B, V.O, E.vB, J.T & A.K was provided by a grant from the European Research Council (Independent Starting Grant No. 637412 ‘META-STRESS’ to MS) and J Ö-U, V.A and D.B from the Academy of Finland grant (Decision No. 304041 to MS & Decision No. 283108 to Ilkka Hanski). A.D was funded by a Marie Sklodowska Curie Individual Fellowship (#790531, Host Sweet Home). O-P.S. was supported by the “TTÜ development program 2016– 2022”, project code 2014-2020.4.01.16-0032.

## Authors’ contributions

O-P.S assembled the genome, processed the chimeric contigs, performed the gap filling and the polishing of the assembly, and participated in the genome analysis.

V.A was responsible for the initial idea of the approach for the genome related activities, coordinated first part of the project, designed and produced data for the linkage map, worked on solving the haplotypes from the initial assembly.

D.B performed gene prediction, functional annotation, ortholog prediction, and managed the manual annotation.

J.K installed and managed the Apollo annotation server.

S.I was responsible for larval rearing and preparation of butterfly crosses.

P.R performed the linkage mapping and anchored the genome onto chromosomes.

V.O assembled the transcriptomes used for gene prediction.

Lo.P manually inspected the chimeric contigs.

A.R performed DNA extraction.

D.B, J.K, V.O, T.D, M.F.D, A.D, I.C.D, P.H, A.K, S.S.K, S.O.K, E.L, S.L, J.M, A.N, M.C-M, V.P, T.S, A.I.T, V.T, E.vB, J.Ö-U and M.S participated in manual annotation.

J.T performed the annotation of transposable elements and repeat classes.

L.P was responsible for the management of the DNA sequencing.

M.J.F was responsible for the management of the genome analysis.

P.A was responsible for the initial idea of the approach for the genome related activities, and the management of the genome analysis.

M.S was responsible for the management of the *M. cinxia* database and genome analysis.

D.B, O-P.S, V.A, P.R, J.T, J.K, V.O, L.P, M.J.F, P.A and M.S wrote the manuscript.

## Acknowledgements

The authors wish to acknowledge CSC – IT Center for Science, Finland, for computational resources. We thank Torsti Schulz and Emily Hornett for annotating > 10 genes. We thank the personnel of the DNA sequencing and genomics laboratory (Institute of Biotechnology, Helsinki, Finland) for performing the NGS sequencing.

## Notes

### Competing Interest Statement

The authors have declared no competing interest.

