## Supplementary material for "Improved chromosome level genome assembly of the Glanville fritillary butterfly (*Melitaea cinxia*) based on SMRT Sequencing and linkage map": AAI.html: AAI.html

AAI-profiler


AAI-profiler home

### AAI-profiler


Toggle parameters

AAI method
 Onesided
 Bidirectional

Scatterplot
Exclude data points with AAI less than   

AAI histograms
Show top  species with AAI values between
 and 
and matched fraction between  and 

#### AAI of Uniprot species

Species are grouped and coloured by genus. Literature suggests > 95 % identity at the species boundary. Matched fraction is maximal in completely sequenced genomes.
Use Plotly option (at top right of plot area) to show closest data on hover and to autoscale or zoom.

#### AAI histograms of Uniprot species

##### 1. Spodoptera frugiperda

Average amino acid identity = 0.741, Median amino acid identity = 0.76, Matched fraction = 0.544, Multiplicity = 1.0
  
Lineage: Eukaryota; Metazoa; Ecdysozoa; Arthropoda; Hexapoda; Insecta; Pterygota; Neoptera; Holometabola; Lepidoptera; Glossata; Ditrysia; Noctuoidea; Noctuidae; Amphipyrinae; Spodoptera

##### 2. Danaus plexippus plexippus

Average amino acid identity = 0.773, Median amino acid identity = 0.80, Matched fraction = 0.519, Multiplicity = 1.0
  
Lineage: Eukaryota; Metazoa; Ecdysozoa; Arthropoda; Hexapoda; Insecta; Pterygota; Neoptera; Holometabola; Lepidoptera; Glossata; Ditrysia; Papilionoidea; Nymphalidae; Danainae; Danaini; Danaina; Danaus; Danaus

##### 3. Papilio xuthus

Average amino acid identity = 0.747, Median amino acid identity = 0.77, Matched fraction = 0.523, Multiplicity = 1.0
  
Lineage: Eukaryota; Metazoa; Ecdysozoa; Arthropoda; Hexapoda; Insecta; Pterygota; Neoptera; Holometabola; Lepidoptera; Glossata; Ditrysia; Papilionoidea; Papilionidae; Papilioninae; Papilio

##### 4. Papilio machaon

Average amino acid identity = 0.745, Median amino acid identity = 0.77, Matched fraction = 0.507, Multiplicity = 1.0
  
Lineage: Eukaryota; Metazoa; Ecdysozoa; Arthropoda; Hexapoda; Insecta; Pterygota; Neoptera; Holometabola; Lepidoptera; Glossata; Ditrysia; Papilionoidea; Papilionidae; Papilioninae; Papilio

##### 5. Chilo suppressalis

Average amino acid identity = 0.739, Median amino acid identity = 0.76, Matched fraction = 0.495, Multiplicity = 1.0
  
Lineage: Eukaryota; Metazoa; Ecdysozoa; Arthropoda; Hexapoda; Insecta; Pterygota; Neoptera; Holometabola; Lepidoptera; Glossata; Ditrysia; Pyraloidea; Crambidae; Crambinae; Chilo

##### 6. Bombyx mori

Average amino acid identity = 0.737, Median amino acid identity = 0.76, Matched fraction = 0.495, Multiplicity = 1.0
  
Lineage: Eukaryota; Metazoa; Ecdysozoa; Arthropoda; Hexapoda; Insecta; Pterygota; Neoptera; Holometabola; Lepidoptera; Glossata; Ditrysia; Bombycoidea; Bombycidae; Bombycinae; Bombyx

##### 7. Heliothis virescens

Average amino acid identity = 0.742, Median amino acid identity = 0.76, Matched fraction = 0.491, Multiplicity = 1.0
  
Lineage: Eukaryota; Metazoa; Ecdysozoa; Arthropoda; Hexapoda; Insecta; Pterygota; Neoptera; Holometabola; Lepidoptera; Glossata; Ditrysia; Noctuoidea; Noctuidae; Heliothinae; Heliothis

##### 8. Leptidea sinapis

Average amino acid identity = 0.734, Median amino acid identity = 0.75, Matched fraction = 0.495, Multiplicity = 1.0
  
Lineage: Eukaryota; Metazoa; Ecdysozoa; Arthropoda; Hexapoda; Insecta; Pterygota; Neoptera; Holometabola; Lepidoptera; Glossata; Ditrysia; Papilionoidea; Pieridae; Dismorphiinae; Leptidea

##### 9. Eumeta japonica

Average amino acid identity = 0.700, Median amino acid identity = 0.71, Matched fraction = 0.472, Multiplicity = 1.0
  
Lineage: Eukaryota; Metazoa; Ecdysozoa; Arthropoda; Hexapoda; Insecta; Pterygota; Neoptera; Holometabola; Lepidoptera; Glossata; Ditrysia; Tineoidea; Psychidae; Oiketicinae; Eumeta

##### 10. Operophtera brumata

Average amino acid identity = 0.715, Median amino acid identity = 0.73, Matched fraction = 0.445, Multiplicity = 1.0
  
Lineage: Eukaryota; Metazoa; Ecdysozoa; Arthropoda; Hexapoda; Insecta; Pterygota; Neoptera; Holometabola; Lepidoptera; Glossata; Ditrysia; Geometroidea; Geometridae; Larentiinae; Operophtera

##### 11. Pararge aegeria

Average amino acid identity = 0.824, Median amino acid identity = 0.85, Matched fraction = 0.317, Multiplicity = 1.0
  
Lineage: Eukaryota; Metazoa; Ecdysozoa; Arthropoda; Hexapoda; Insecta; Pterygota; Neoptera; Holometabola; Lepidoptera; Glossata; Ditrysia; Papilionoidea; Nymphalidae; Satyrinae; Satyrini; Parargina; Pararge

##### 12. Helicoverpa armigera

Average amino acid identity = 0.728, Median amino acid identity = 0.74, Matched fraction = 0.275, Multiplicity = 1.0
  
Lineage: Eukaryota; Metazoa; Ecdysozoa; Arthropoda; Hexapoda; Insecta; Pterygota; Neoptera; Holometabola; Lepidoptera; Glossata; Ditrysia; Noctuoidea; Noctuidae; Heliothinae; Helicoverpa

##### 13. Pectinophora gossypiella

Average amino acid identity = 0.754, Median amino acid identity = 0.78, Matched fraction = 0.261, Multiplicity = 1.0
  
Lineage: Eukaryota; Metazoa; Ecdysozoa; Arthropoda; Hexapoda; Insecta; Pterygota; Neoptera; Holometabola; Lepidoptera; Glossata; Ditrysia; Gelechioidea; Gelechiidae; Pexicopiinae; Pectinophora

##### 14. Tribolium castaneum

Average amino acid identity = 0.599, Median amino acid identity = 0.59, Matched fraction = 0.320, Multiplicity = 1.0
  
Lineage: Eukaryota; Metazoa; Ecdysozoa; Arthropoda; Hexapoda; Insecta; Pterygota; Neoptera; Holometabola; Coleoptera; Polyphaga; Cucujiformia; Tenebrionidae; Tenebrionidae incertae sedis; Tribolium

##### 15. Cryptotermes secundus

Average amino acid identity = 0.604, Median amino acid identity = 0.60, Matched fraction = 0.317, Multiplicity = 1.0
  
Lineage: Eukaryota; Metazoa; Ecdysozoa; Arthropoda; Hexapoda; Insecta; Pterygota; Neoptera; Polyneoptera; Dictyoptera; Blattodea; Blattoidea; Termitoidae; Kalotermitidae; Cryptotermitinae; Cryptotermes

##### 16. Photinus pyralis

Average amino acid identity = 0.591, Median amino acid identity = 0.59, Matched fraction = 0.313, Multiplicity = 1.0
  
Lineage: Eukaryota; Metazoa; Ecdysozoa; Arthropoda; Hexapoda; Insecta; Pterygota; Neoptera; Holometabola; Coleoptera; Polyphaga; Elateriformia; Elateroidea; Lampyridae; Lampyrinae; Photinus

##### 17. Ooceraea biroi

Average amino acid identity = 0.596, Median amino acid identity = 0.59, Matched fraction = 0.302, Multiplicity = 1.0
  
Lineage: Eukaryota; Metazoa; Ecdysozoa; Arthropoda; Hexapoda; Insecta; Pterygota; Neoptera; Holometabola; Hymenoptera; Apocrita; Aculeata; Formicoidea; Formicidae; Dorylinae; Ooceraea

##### 18. Aedes aegypti

Average amino acid identity = 0.595, Median amino acid identity = 0.59, Matched fraction = 0.295, Multiplicity = 1.0
  
Lineage: Eukaryota; Metazoa; Ecdysozoa; Arthropoda; Hexapoda; Insecta; Pterygota; Neoptera; Holometabola; Diptera; Nematocera; Culicoidea; Culicidae; Culicinae; Aedini; Aedes; Stegomyia

##### 19. Graphocephala atropunctata

Average amino acid identity = 0.606, Median amino acid identity = 0.60, Matched fraction = 0.289, Multiplicity = 1.0
  
Lineage: Eukaryota; Metazoa; Ecdysozoa; Arthropoda; Hexapoda; Insecta; Pterygota; Neoptera; Paraneoptera; Hemiptera; Auchenorrhyncha; Membracoidea; Cicadellidae; Cicadellinae; Cicadellini; Graphocephala

##### 20. Dendroctonus ponderosae

Average amino acid identity = 0.598, Median amino acid identity = 0.59, Matched fraction = 0.292, Multiplicity = 1.0
  
Lineage: Eukaryota; Metazoa; Ecdysozoa; Arthropoda; Hexapoda; Insecta; Pterygota; Neoptera; Holometabola; Coleoptera; Polyphaga; Cucujiformia; Curculionidae; Scolytinae; Dendroctonus
