## Supplementary material for "Improved chromosome level genome assembly of the Glanville fritillary butterfly (*Melitaea cinxia*) based on SMRT Sequencing and linkage map": krona.html: krona.html

Javascript must be enabled to view this page.

magnitude
magnitudeUnassigned

kt

10988.785

10966.835

2.81

2.81

2.555

0.425

0.425

0.425

0.425

2.13

2.13

2.13

0.225

0.225

0.375

0.375

1.53

0.215

0.205

0.715

0.395

0.255

0.255

0.255

0.255

0.255

0.255

0.255

0.815

0.815

0.815

0.265

0.265

0.265

0.55

0.55

0.55

10941.265

0.63

0.63

0.63

0.63

0.63

0.63

0.63

0.63

10881.76

6.745

4.87

2.84

0.355

0.355

0.355

0.355

0.92

0.92

0.415

0.415

0.505

0.505

0.555

0.555

0.555

0.555

0.555

1.01

1.01

1.01

1.01

1.01

2.03

0.83

0.83

0.83

0.83

0.335

0.335

0.495

0.495

1.2

1.2

1.2

1.2

1.2

1.2

1.875

1.875

1.875

1.875

1.875

0.415

0.715

0.745

0.745

10872.55

10846.825

10839.185

10839.185

10839.185

29.955

27.78

27.78

11.18

11.18

11.18

11.18

11.18

16.6

16.6

7.135

7.135

7.135

7.135

9.465

9.465

9.465

2.175

2.175

2.175

2.175

2.175

2.175

2.175

2.175

2.175

76.38

2.415

2.415

2.415

2.415

2.415

2.415

73.965

23.26

23.26

3.785

3.785

3.785

3.785

3.785

3.785

19.475

10.37

10.37

1.21

0.785

0.425

6.18

4.715

1.465

2.98

2.98

9.105

9.105

9.105

9.105

21.095

11.91

11.91

11.91

7.94

4.47

4.47

3.47

3.47

3.97

3.97

3.97

2.13

2.13

2.13

2.13

2.13

7.055

7.055

7.055

7.055

29.61

7.495

7.495

7.495

7.495

21.67

21.67

19.87

13.86

13.86

13.86

3.715

2.665

2.665

1.05

1.05

2.295

2.295

1.8

1.8

1.8

0.445

0.445

0.445

0.445

10732.85

10484.08

10484.08

10483.455

0.495

0.495

0.495

0.495

255.355

255.355

255.355

255.355

255.355

276.39

276.39

276.39

276.39

276.39

284.225

284.225

280.735

280.735

280.735

3.49

0.705

0.705

2.785

2.785

1490.03

1487.735

7.565

6.495

0.575

0.575

5.92

5.195

0.725

1.07

0.535

0.535

41.79

1.68

0.945

0.735

40.11

12.66

12.695

14.755

5.345

5.345

5.345

1424.975

996.725

996.725

428.25

3.385

1.39

423.475

8.06

2.935

2.935

5.125

5.125

1.3

1.3

0.695

0.695

0.605

0.605

0.995

0.995

0.995

0.995

13.25

13.25

13.25

13.25

1188.99

1173.645

1156.19

1156.19

0.885

1155.305

3.515

3.515

12.71

1.845

1.845

10.865

0.935

4.985

4.945

1.23

0.425

0.425

0.805

0.805

15.345

11.065

11.065

11.065

4.28

0.625

0.625

3.655

3.655

0.805

0.805

0.805

0.805

0.805

6386.28

634.015

21.79

21.79

2.785

19.005

612.225

612.225

612.225

1560.405

0.995

0.995

0.995

0.995

0.995

1559.41

0.895

0.895

1558.515

1.55

733.565

793.51

0.995

28.895

4191.86

62.805

40.7

0.925

39.775

3.93

3.93

3.37

3.37

13.93

13.93

0.875

0.875

3082.185

3082.185

3082.185

1.99

1.99

3080.195

3080.195

3070.6

9.595

154.56

0.985

0.985

0.985

0.985

0.985

0.985

152.59

0.535

0.535

2.83

2.83

0.715

0.715

148.51

72.05

3

0.565

0.735

3.4

7.83

0.595

24.245

1.06

31.36

0.855

1.725

1.09

0.995

0.995

0.995

0.995

891.315

891.315

1.96

0.995

0.995

0.965

0.965

38.87

38.87

38.87

850.485

850.485

850.485

560.29

2.77

2.77

0.695

2.075

27.19

27.19

27.19

27.19

27.19

12.74

11.43

2.44

2.44

2.44

8.99

8.99

3.485

5.505

1.31

1.31

1.31

517.59

517.59

517.59

517.59

17.52

17.52

5.295

1.815

1.815

3.48

3.48

12.225

9.26

3.365

3.365

0.925

0.925

4.97

3.86

1.11

2.965

1.05

1.05

1.915

1.915

0.625

0.625

0.625

0.625

0.625

9.2

9.2

9.2

9.2

9.2

0.625

0.625

0.625

0.625

0.625

0.475

0.475

0.475

0.475

0.475

0.475

0.475

60.57

60.57

24.665

0.795

0.795

0.795

0.795

0.795

10.425

10.425

10.425

10.425

10.425

13.445

13.445

13.445

13.445

13.445

33.195

6.725

6.23

6.23

6.23

6.23

6.23

0.495

0.495

0.495

0.495

0.495

8.96

8.96

8.96

8.96

17.51

10.52

10.52

10.52

6.99

6.99

6.99

2.71

2.71

2.71

2.71

2.71

90.795

0.315

0.315

0.315

0.315

90.48

80.87

56.425

56.425

3.33

3.33

3.33

3.33

26.395

2.585

2.585

23.81

23.81

23.81

19.705

2.63

2.63

2.79

2.79

2.79

2.79

3.92

2.66

1.26

2.955

2.955

4.62

3.3

1.32

6.995

6.995

6.995

24.445

22.57

7.02

7.02

10.245

2.965

0.945

3.115

3.22

1.54

1.54

3.765

3.765

1.875

1.875

1.875

1.875

9.61

3.935

3.935

0.475

0.475

0.475

3.46

3.46

3.46

5.675

5.675

5.675

0.775

0.775

4.9

4.9

96.93

52.935

37.995

2.29

2.29

2.29

35.705

24.73

24.73

0.865

3.035

0.425

0.625

1.24

1.24

1.345

0.565

0.595

1.27

4.285

3.88

2.055

0.865

2.54

0.745

0.395

10.975

9.18

9.18

9.18

2.7

6.48

1.795

1.795

1.795

0.655

1.14

6.83

6.83

2.785

2.785

2.785

0.615

0.615

3.43

3.43

3.43

8.11

6.77

6.77

6.77

1.34

1.34

1.34

1.34

1.34

43.995

2.085

2.085

2.085

2.085

2.085

41.91

3.74

3.74

1.18

1.18

0.585

0.585

1.975

1.975

13.595

13.595

13.595

1.24

3.605

0.815

2.16

3.535

2.24

1.905

1.905

1.905

1.905

1.905

5.41

5.41

0.99

0.99

0.99

0.685

0.685

0.685

3.735

3.735

2.16

0.98

0.595

15.755

15.755

15.755

13.99

0.745

1.13

1.915

0.645

1.845

2.865

4.845

0.395

1.37

1.505

1.505

1.505

1.505

1.505

7.64

7.64

1.11

1.11

1.11

1.11

1.11

6.53

6.53

6.53

6.53

6.53

9.14

5.445

0.645

0.645

0.645

0.645

0.645

0.645

4.8

4.8

3.18

3.18

0.91

0.91

0.91

0.91

0.91

0.91

0.91

2.27

2.27

2.27

2.27

2.27

1.62

0.93

0.93

0.93

0.93

0.93

0.93

0.69

0.69

0.69

0.69

0.69

0.69

3.695

3.695

3.695

3.695

3.695

3.695

3.26

0.435

16.585

16.585

3.41

3.41

3.41

3.41

1.535

0.375

0.375

0.375

1.16

1.16

1.16

0.435

0.725

1.875

1.875

0.555

1.32

12.62

12.62

12.62

0.465

0.465

0.465

0.465

12.155

1.07

1.07

1.07

11.085

11.085

11.085

0.555

0.555

0.555

0.555

0.555

0.555

2.465

2.465

2.465

2.465

2.465

2.465

2.465

48.59

1.995

1.995

1.995

1.995

46.595

46.595

0.595

0.595

0.595

0.595

0.595

0.595

45.575

3.69

0.545

0.545

0.545

0.545

0.545

0.545

3.145

2.25

2.25

2.25

2.25

2.25

2.25

0.695

0.695

0.695

0.695

0.93

0.93

0.305

0.305

0.305

0.305

0.625

0.625

0.625

0.625

0.625

0.625

0.625

0.895

0.895

0.895

0.895

0.895

1.725

1.725

1.725

1.725

1.725

1.725

0.795

0.795

0.795

0.795

0.795

0.795

0.93

0.93

0.93

0.93

25.195

25.195

25.195

1.225

0.59

0.59

0.59

0.59

0.635

0.635

0.635

0.635

0.635

17.7

17.7

1.635

0.335

0.335

0.335

0.335

0.705

0.705

0.705

0.705

0.595

0.595

0.595

0.595

0.595

0.595

1.44

1.44

0.435

0.435

0.435

1.005

1.005

1.005

5.715

2.56

0.715

0.715

0.715

0.715

0.715

1.845

1.845

1.845

1.845

1.845

1.965

1.965

1.965

1.19

0.475

0.475

0.475

0.475

0.715

0.715

0.715

8.91

2.52

2.52

2.52

2.125

2.125

0.535

0.535

0.535

0.925

0.925

0.925

0.665

0.665

0.665

0.395

0.395

0.395

0.395

0.395

5.995

3.33

3.33

3.33

3.33

0.505

2.825

2.665

0.465

0.465

0.465

0.655

0.655

0.655

1.545

1.545

0.585

0.585

0.575

0.575

0.385

0.385

0.395

0.395

0.395

1.855

1.855

1.855

1.35

1.35

1.35

0.505

0.505

0.505

4.415

0.315

0.315

0.315

0.315

0.315

1.02

1.02

0.405

0.405

0.615

0.615

0.615

3.08

3.08

1.575

1.575

0.85

0.85

0.655

0.655

1.18

1.18

1.18

1.18

1.39

1.39

1.39

0.855

0.855

0.855

0.855

0.855

0.855

0.535

0.535

0.535

0.535

0.535

0.535

12.395

0.445

0.445

0.445

0.445

0.445

11.95

7.585

4.53

4.53

2.855

1.23

0.515

0.515

0.715

0.715

0.535

0.535

0.535

1.09

1.09

1.09

0.615

0.475

1.675

1.675

0.715

0.715

0.715

0.96

0.96

0.96

0.96

0.835

0.835

0.835

0.835

2.22

2.22

2.22

2.22

2.22

0.585

0.585

0.585

1.635

1.635

1.635

4.365

1.915

1.915

0.615

0.615

0.615

1.3

0.735

0.735

0.565

0.565

2.45

0.615

0.615

0.615

0.615

0.765

0.765

0.765

0.765

0.765

0.475

0.475

0.475

0.475

0.475

0.475

0.595

0.595

0.595

0.595

0.425

0.425

0.425

0.425

0.425

0.425

0.425

0.425

7.535

0.795

0.795

0.795

0.795

0.795

0.795

0.795

0.795

0.725

0.725

0.725

0.725

0.725

0.725

0.725

6.015

3.25

1.645

1.645

1.645

1.645

1.645

1.645

1.645

1.645

1.605

0.495

0.495

0.495

0.495

0.495

1.11

1.11

1.11

1.11

1.11

2.02

2.02

2.02

2.02

2.02

2.02

2.02

2.02

2.02

0.745

0.745

0.745

0.745

0.745

0.745

0.745

2.75

0.78

0.78

0.465

0.465

0.465

0.465

0.465

0.315

0.315

0.315

0.315

0.315

0.315

1.435

1.435

1.435

1.435

1.435

1.435

0.535

0.535

0.535

0.535

0.535

0.535

0.245

0.245

0.245

0.245

0.245

0.245

0.245

0.245

8.755

2.855

0.55

0.55

0.55

0.55

0.55

0.55

0.55

1.265

0.305

0.305

0.305

0.305

0.305

0.305

0.96

0.555

0.555

0.555

0.555

0.555

0.405

0.405

0.405

0.405

0.405

0.405

0.405

1.04

1.04

1.04

1.04

1.04

0.485

0.485

0.555

0.555

5.9

3.865

3.35

0.73

0.73

0.73

0.73

0.73

0.335

0.395

0.495

0.495

0.495

0.495

0.495

0.9

0.9

0.405

0.405

0.495

0.495

0.495

0.495

1.225

0.81

0.265

0.265

0.265

0.265

0.265

0.545

0.545

0.545

0.545

0.415

0.415

0.415

0.415

0.415

0.415

0.515

0.515

0.515

0.515

0.515

0.515

0.515

2.035

1.42

1.42

1.42

1.42

1.42

0.615

0.805

0.615

0.615

0.615

0.615

0.615

0.615

11.36

10.675

10.02

0.625

0.625

0.625

0.625

0.625

0.625

0.625

9.395

9.395

9.395

1.385

0.395

0.395

0.395

0.395

0.395

0.585

0.585

0.585

0.585

0.585

0.405

0.405

0.405

0.405

0.405

0.405

0.405

0.405

8.01

8.01

8.01

5.5

0.94

0.94

0.94

0.94

0.94

0.94

0.94

0.94

0.94

0.94

0.94

4.56

2.425

2.425

0.935

0.935

0.935

1.49

1.49

0.765

0.725

2.135

2.135

2.135

2.135

2.135

2.51

2.51

2.51

2.51

1.1

0.525

0.525

0.525

0.575

0.575

0.575

1.41

1.41

1.41

1.41

0.655

0.655

0.655

0.655

0.655

0.685

0.685

0.685

0.685

0.685

0.685

0.525

0.525

0.525

0.525

0.525

0.525

0.525

1.06

0.465

0.465

0.465

0.595

0.595

0.595

0.595

0.595

17.425

4.29

0.545

0.545

0.545

0.545

0.485

0.915

0.915

0.915

0.915

0.545

0.545

0.545

0.545

1.8

1.385

0.285

0.285

1.1

0.625

0.475

0.415

0.415

0.415

0.415

2.235

1.72

1.72

0.69

0.325

0.325

0.325

0.365

0.365

0.365

1.03

0.515

5.505

0.145

0.145

0.145

0.145

0.145

2.32

0.405

0.405

0.405

0.405

0.975

0.975

0.975

0.975

0.595

0.595

0.595

0.595

0.345

0.345

0.345

0.345

0.365

0.365

0.365

0.365

0.365

2.675

0.375

0.375

0.375

0.375

0.375

1.795

1.795

1.795

1.795

0.505

1.46

1.46

1.46

1.46

1.46

1.46

2.61

0.72

0.72

0.295

0.425

0.425

0.425

1.89

0.275

0.275

0.275

0.275

1.615

1.615

1.615

0.275

0.545

0.545

0.795

0.485

0.485

0.485

0.485

0.485

0.485

0.365

0.365

0.475

0.475

0.475

0.475

3.885

1.7

1.7

1.7

1.7

1.415

0.635

0.635

0.78

0.78

0.78

0.78

0.78

0.78

0.415

0.365

0.465

0.465

0.465

0.465

0.465

0.305

0.305

0.305

0.305

0.345

0.345

0.345

0.295

0.295

0.295

0.295
