## Supplementary Figures for "Improved chromosome level genome assembly of the Glanville fritillary butterfly (*Melitaea cinxia*) based on SMRT Sequencing and linkage map"

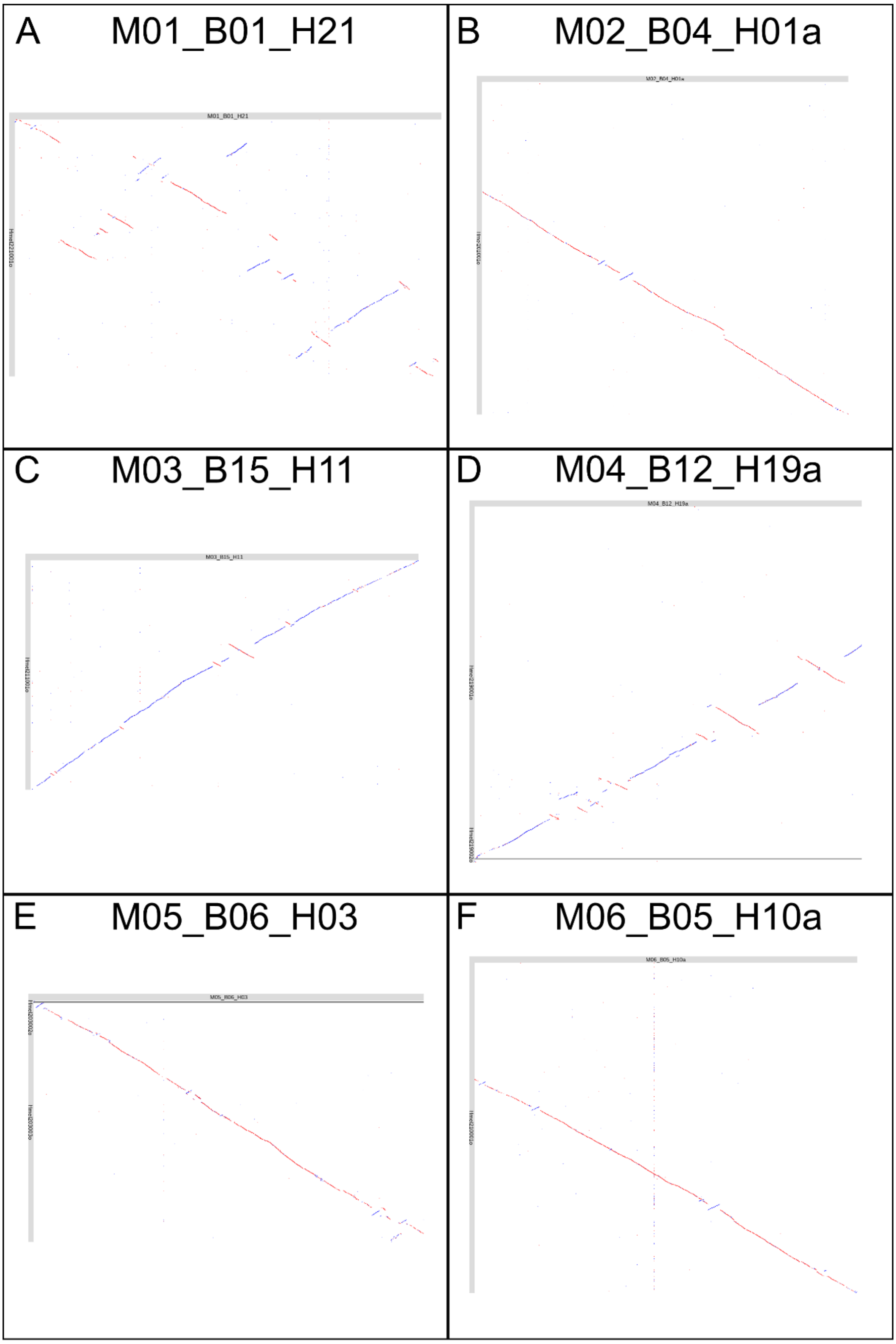


Figure S1: *M. cinxia* aligned against *H. melpomene* using the last aligner (Kielbasa et al. 2011). A: *M. cinxia* chromosome 1 (M01_B01_H21), B: chromosome 2 (M02_B04_H01a), C: chromosome 3 (M03_B15_H11), D: chromosome 4 (M04_B12_H19a), E: chromosome 5 (M05_B06_H03), and F: chromosome 6 (M06_B05_H10a).

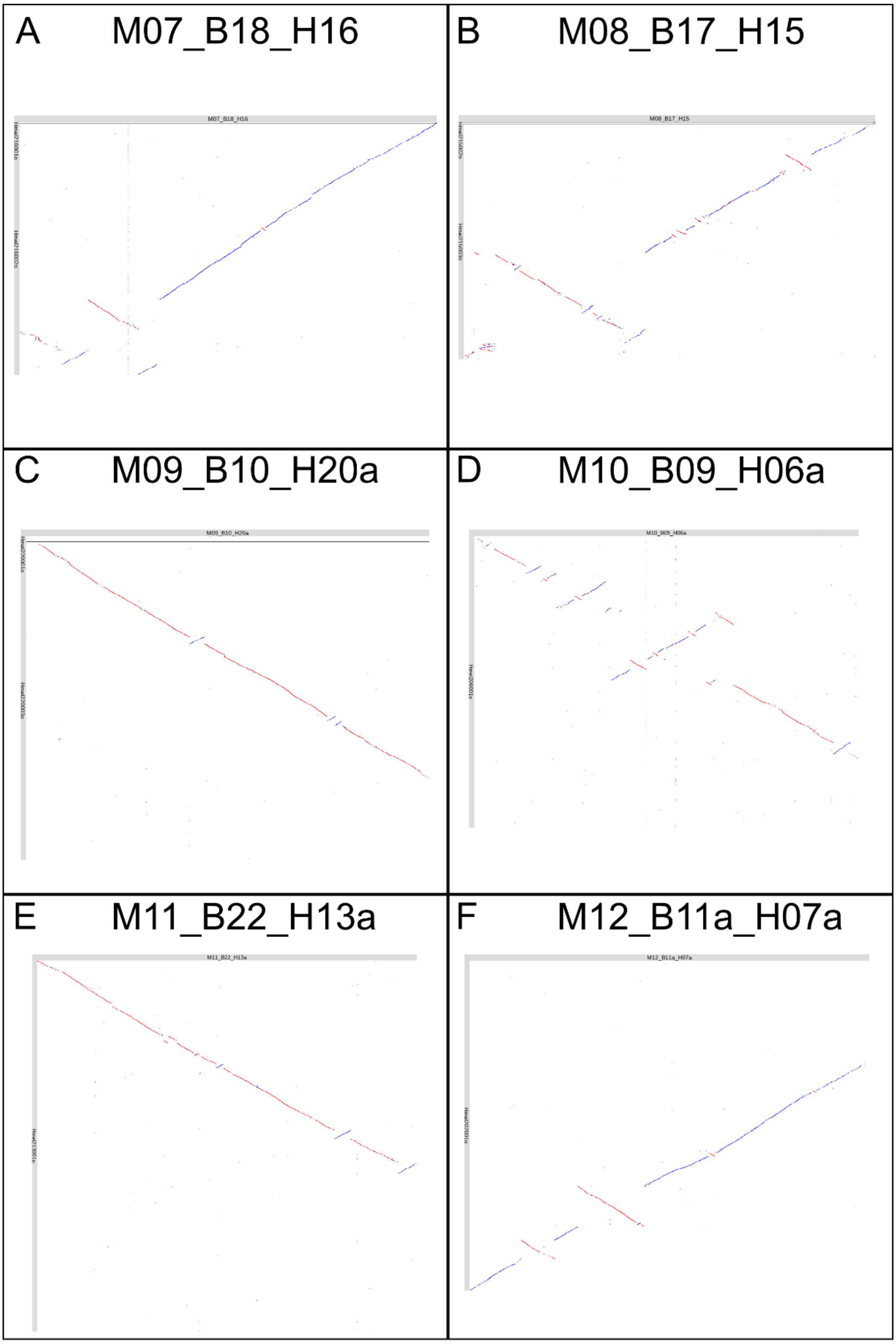


Figure S2: *M. cinxia* aligned against *H. melpomene* using the last aligner (Kielbasa et al. 2011). A: *M. cinxia* chromosome 7 (M07_B18_H16), B: chromosome 8 (M08_B17_H15), C: chromosome 9 (M09_B10_H20a), D: chromosome 10 (M10_B09_H06a), E: chromosome 11 (M11_B22_H13a), and F: chromosome 12 (M12_B11a_H07a).

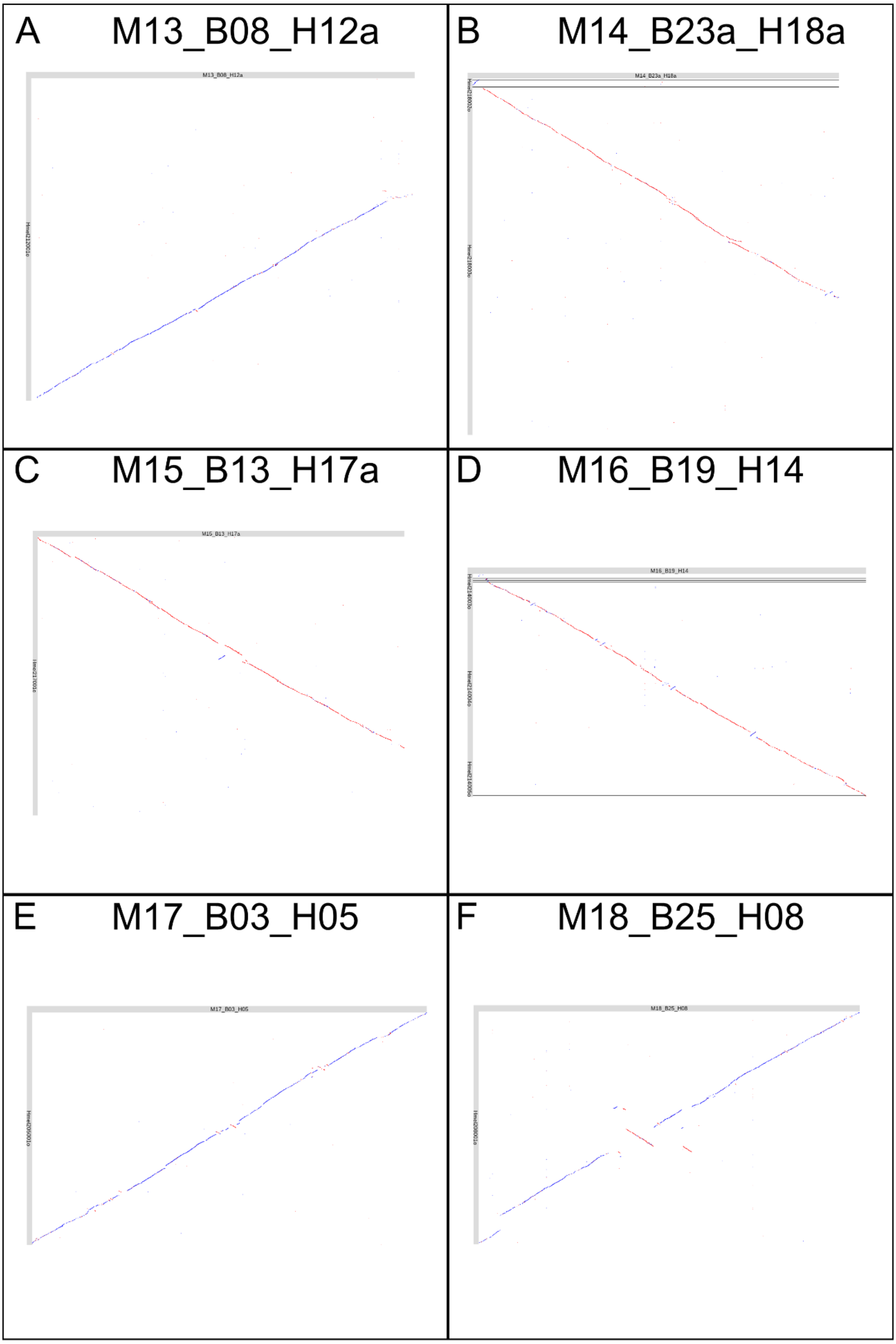


Figure S3: *M. cinxia* aligned against *H. melpomene* using the last aligner (Kielbasa et al. 2011). A: *M. cinxia* chromosome 13 (M13_B08_H12a), B: chromosome 14 (M14_B23a_H18a), C: chromosome 15 (M15_B13_H17a), D: chromosome 16 (M16_B19_H14), E: chromosome 17 (M17_B03_H05), and F: chromosome 18 (M18_B25_H08).

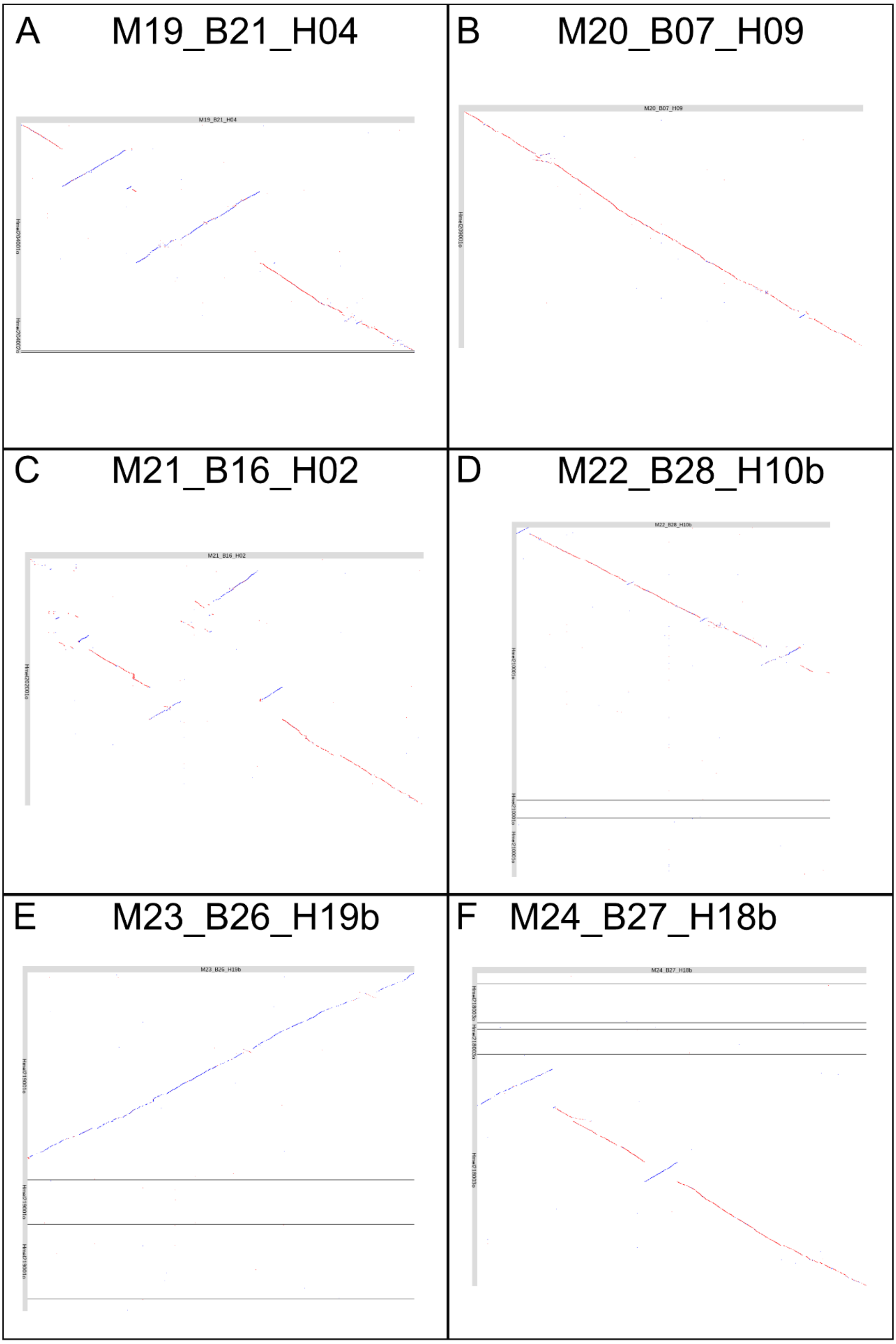


Figure S4: *M. cinxia* aligned against *H. melpomene* using the last aligner (Kielbasa et al. 2011). A: *M. cinxia* chromosome 19 (M19_B21_H04), B: chromosome 20 (M20_B07_H09), C: chromosome 21 (M21_B16_H02), D: chromosome 22 (M22_B28_H10b), E: chromosome 23 (M23_B26_H19b), and F: chromosome 24 (M24_B27_H18b).

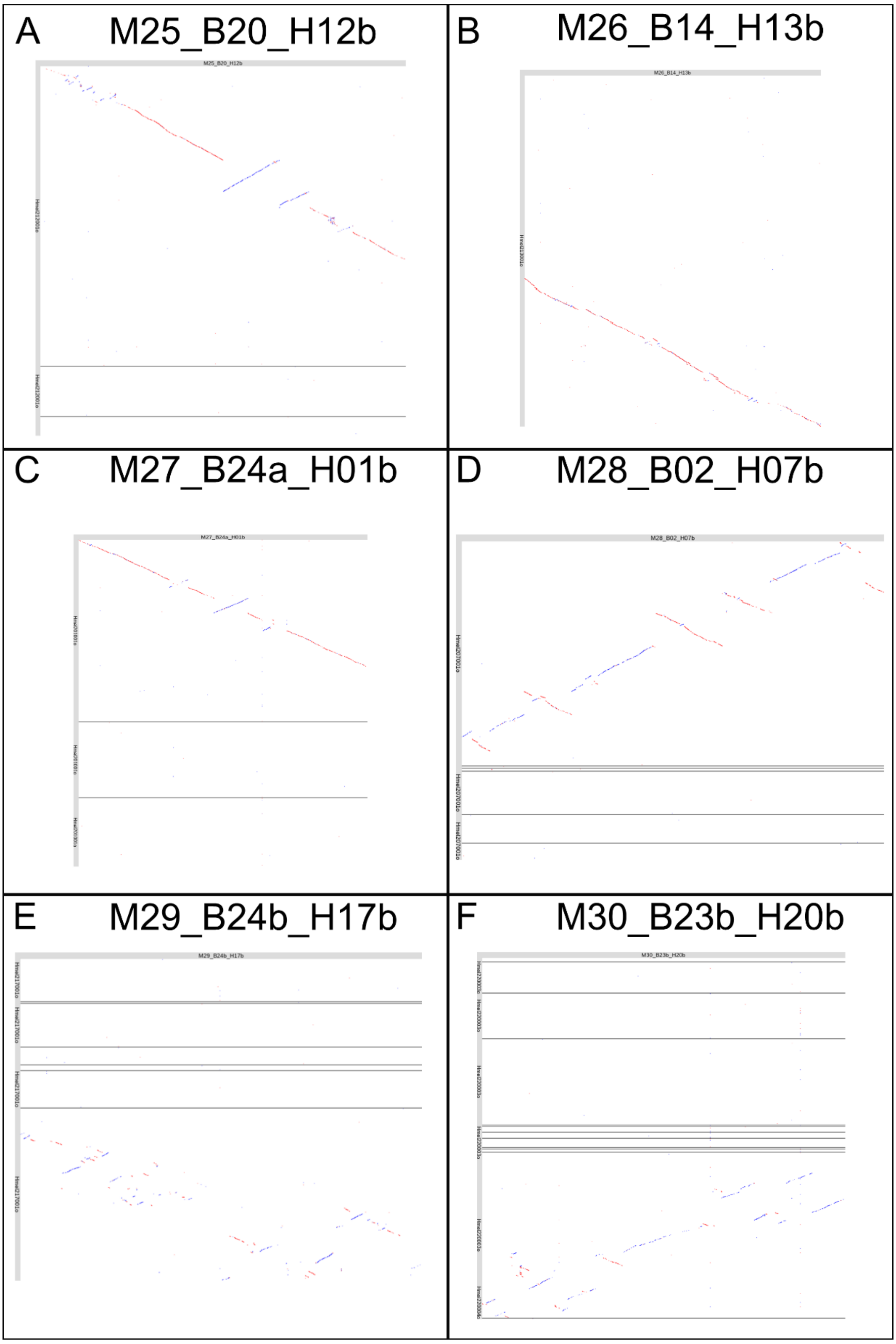


Figure S5: *M. cinxia* aligned against *H. melpomene* using the last aligner (Kielbasa et al. 2011). A: *M. cinxia* chromosome 25 (M25_B20_H12b), B: chromosome 26 (M26_B14_H13b), C: chromosome 27 (M27_B24a_H01b), D: chromosome 28 (M28_B02_H07b), E: chromosome 29 (M29_B24b_H17b), and F: chromosome 30 (M30_B23b_H20b).

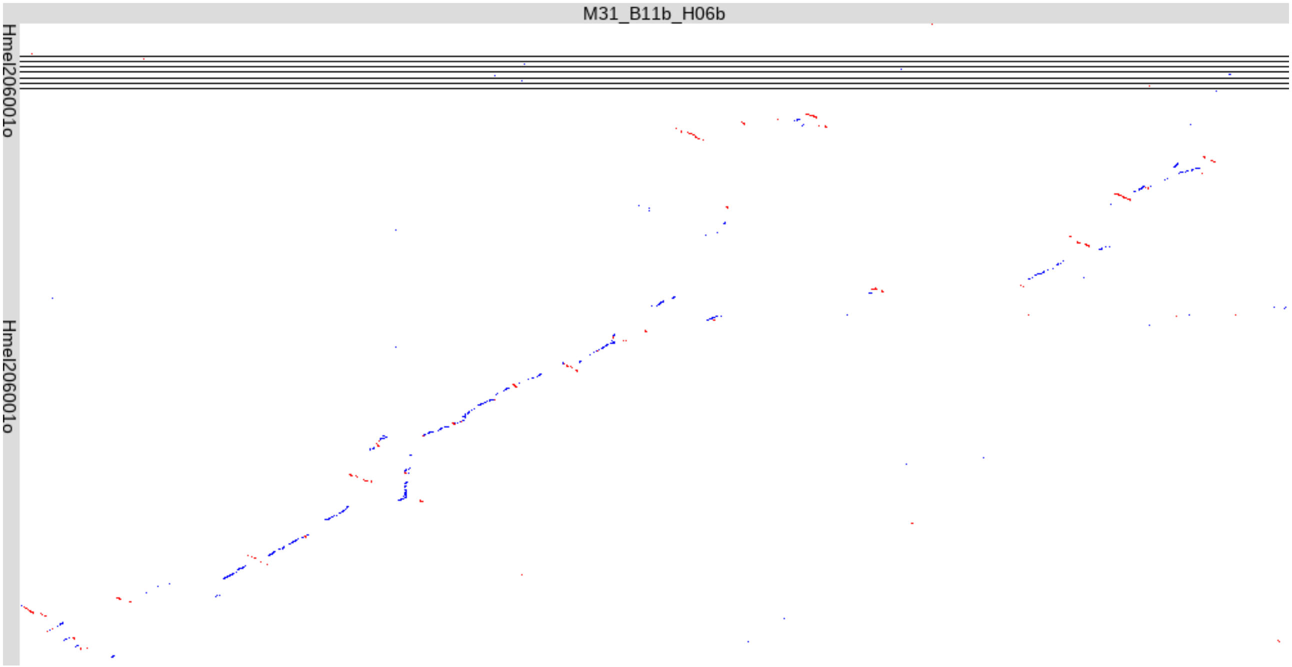


Figure S6: *M. cinxia* aligned against *H. melpomene* using the last aligner (Kielbasa et al. 2011). *M. cinxia* chromosome 31 (M31_B11b_H06b)

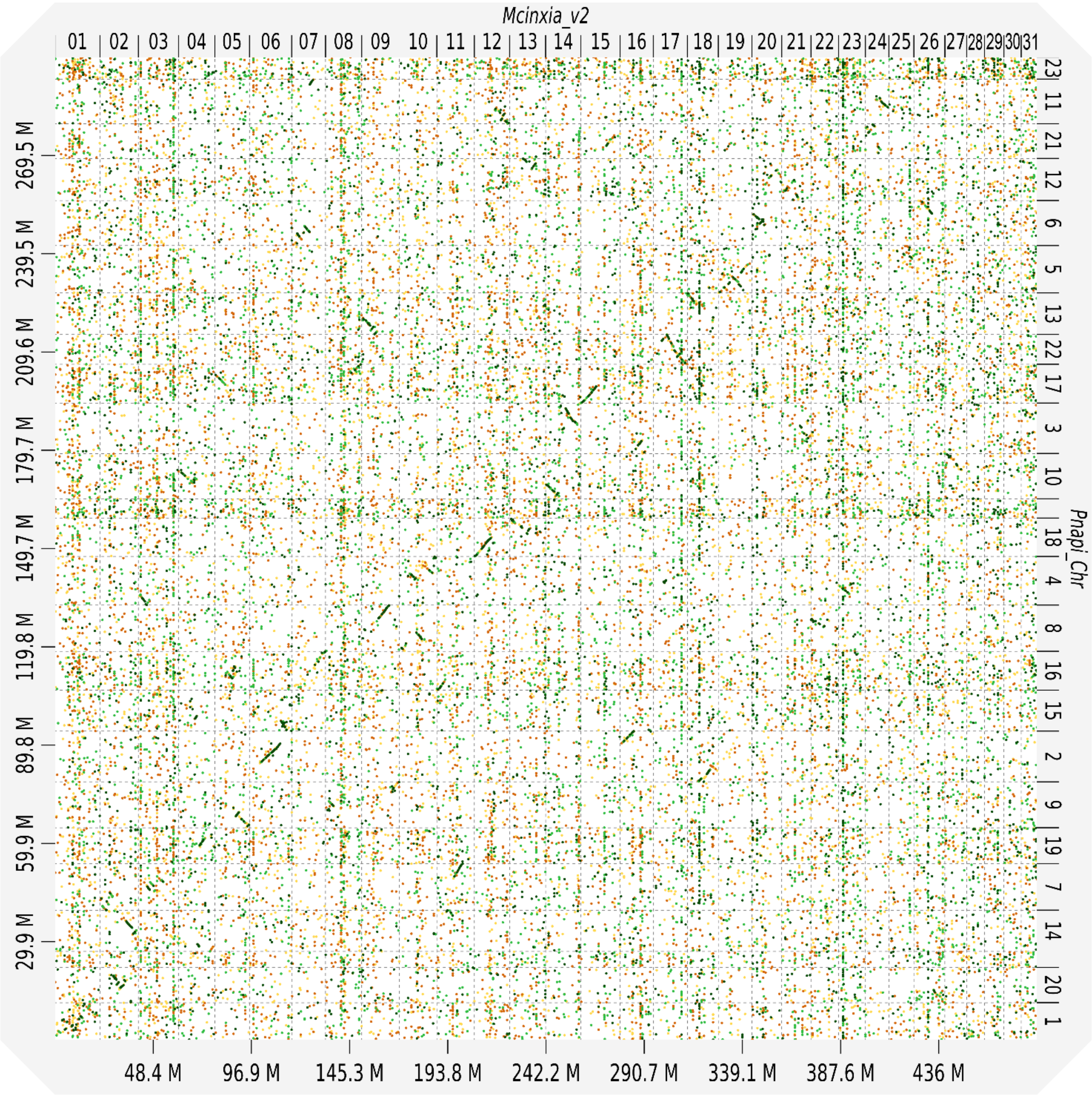


Figure S7: A dot-plot showing the structure of *P. napi* genome against *M. cinxia* genome v.2. The diagonal lines indicate the collinearity between the two species.

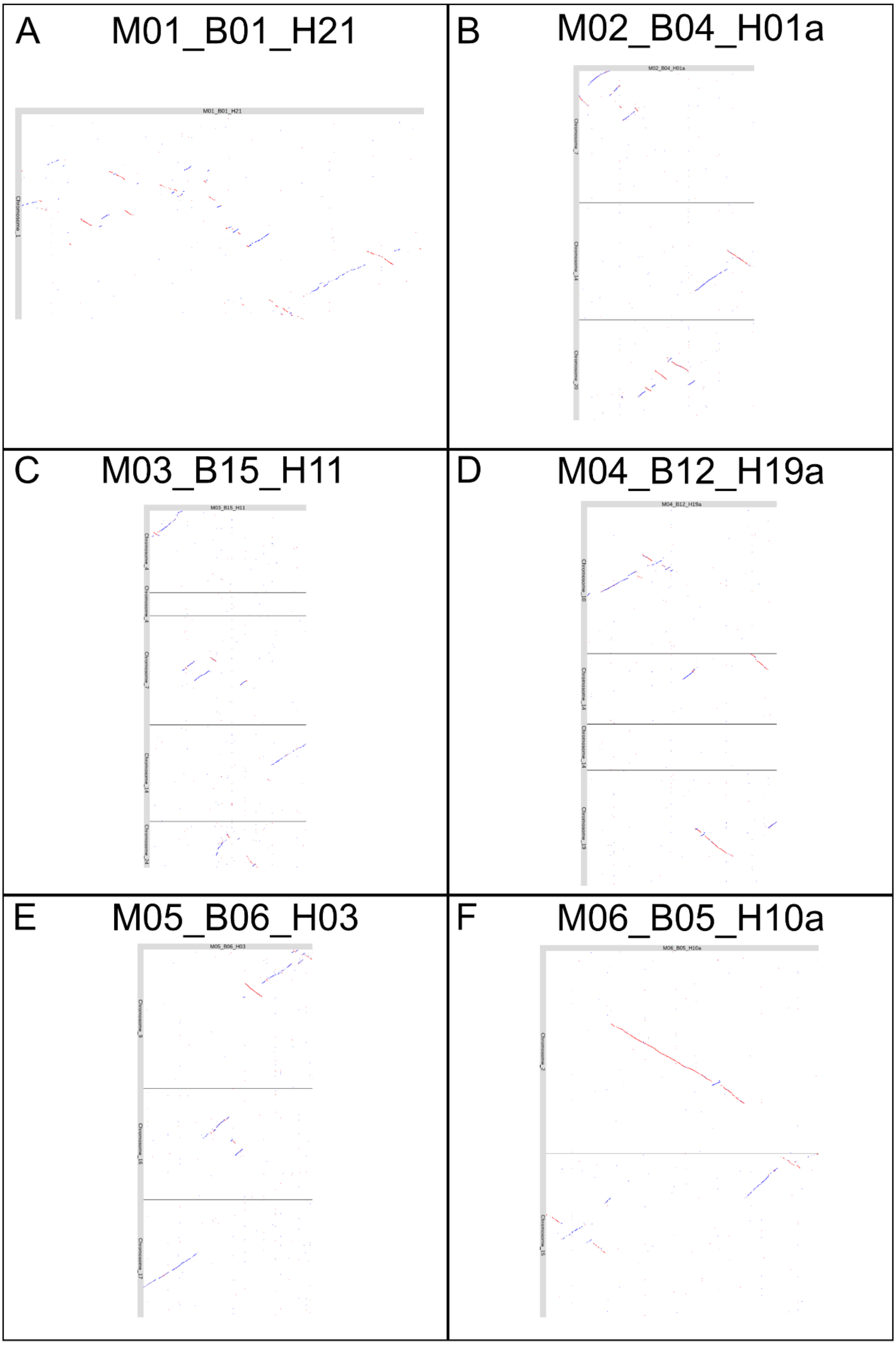


Figure S8: *M. cinxia* aligned against *P. napi* using the last aligner (Kielbasa et al. 2011). A: *M. cinxia* chromosome 1 (M01_B01_H21), B: chromosome 2 (M02_B04_H01a), C: chromosome 3 (M03_B15_H11), D: chromosome 4 (M04_B12_H19a), E: chromosome 5 (M05_B06_H03), and F: chromosome 6 (M06_B05_H10a).

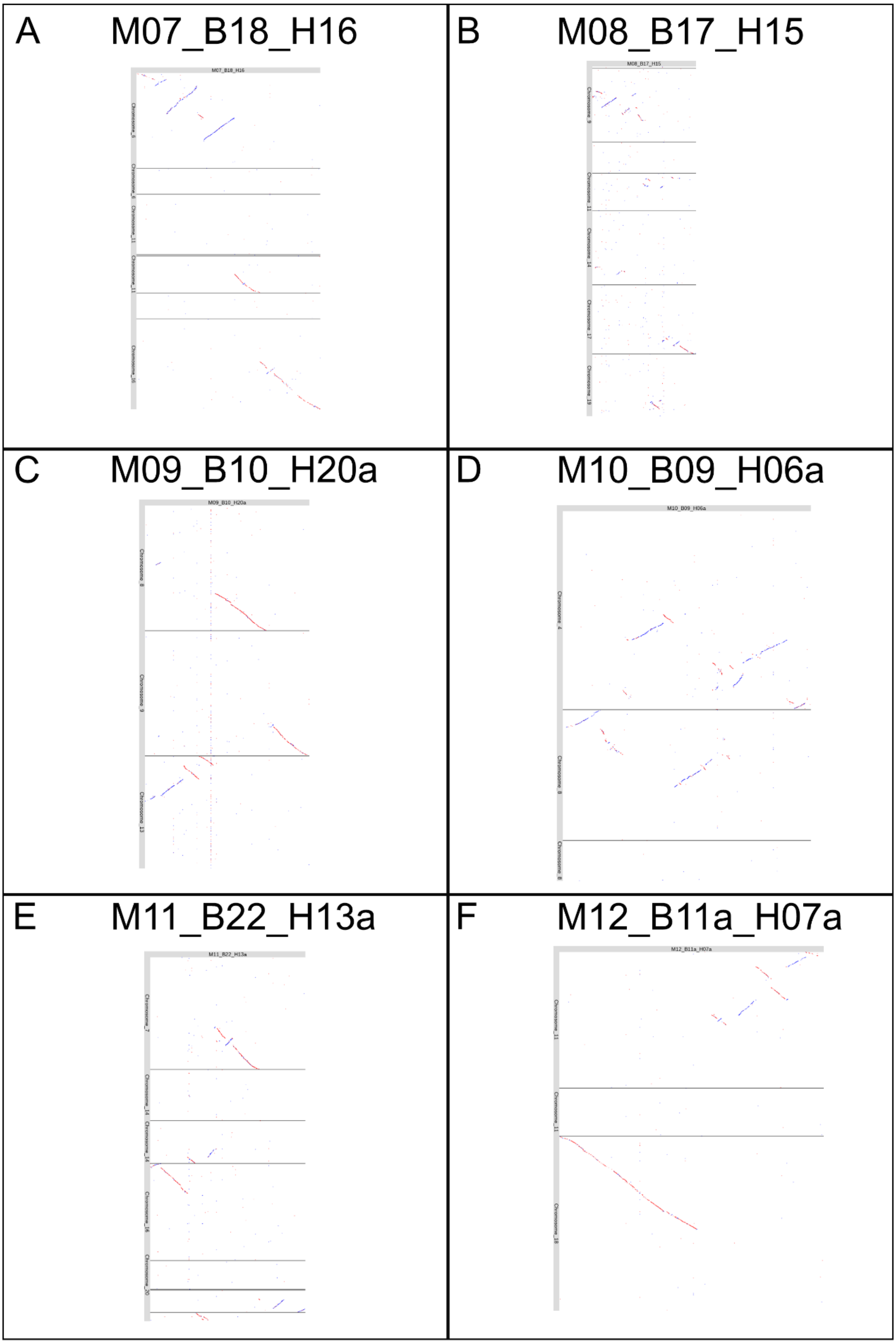


Figure S9: *M. cinxia* aligned against *P. napi* using the last aligner (Kielbasa et al. 2011). A: *M. cinxia* chromosome 7 (M07_B18_H16), B: chromosome 8 (M08_B17_H15), C: chromosome 9 (M09_B10_H20a), D: chromosome 10 (M10_B09_H06a), E: chromosome 11 (M11_B22_H13a), and F: chromosome 12 (M12_B11a_H07a).

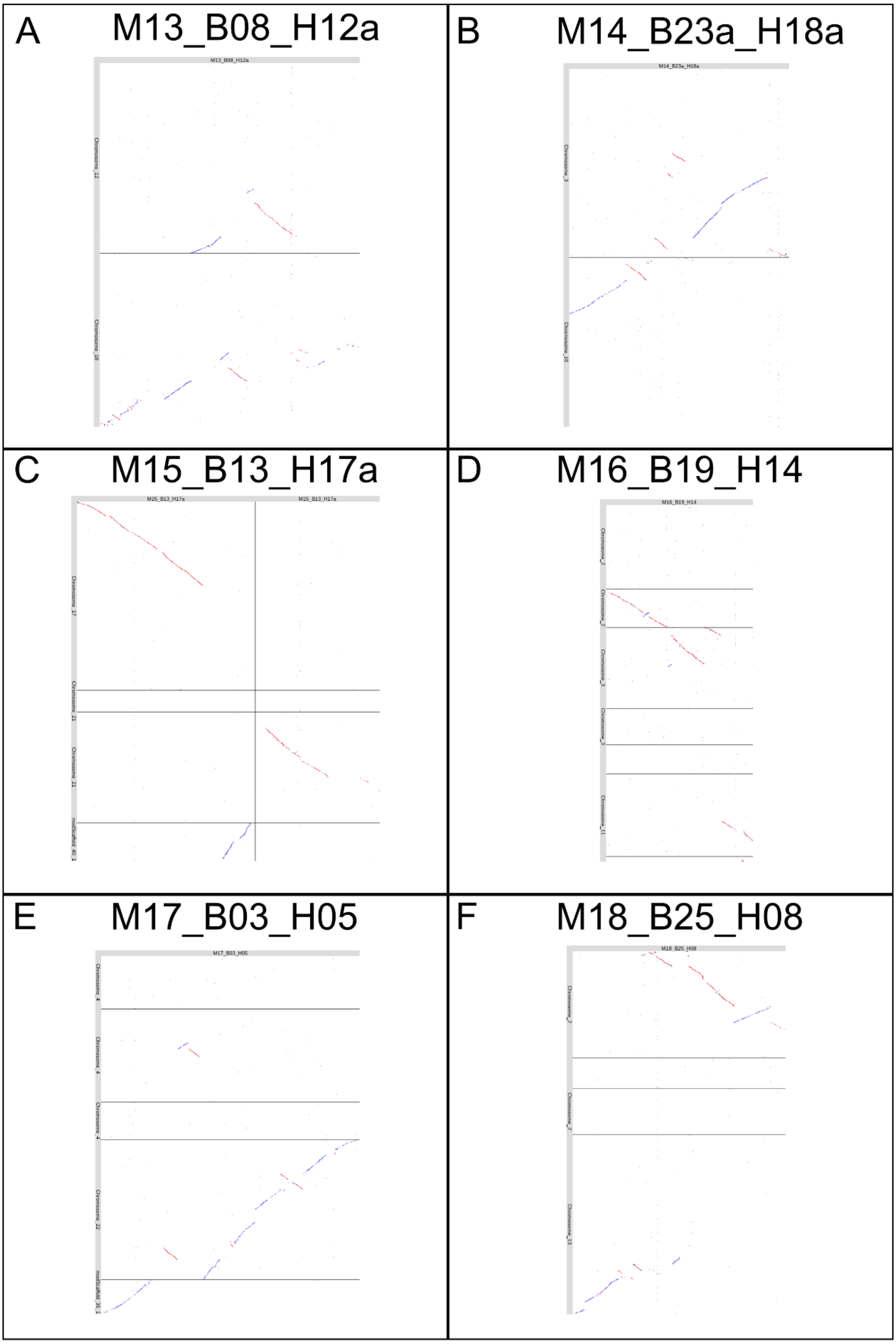


Figure S10: *M. cinxia* aligned against *P. napi* using the last aligner (Kielbasa et al. 2011). A: *M. cinxia* chromosome 13 (M13_B08_H12a), B: chromosome 14 (M14_B23a_H18a), C: chromosome 15 (M15_B13_H17a), D: chromosome 16 (M16_B19_H14), E: chromosome 17 (M17_B03_H05), and F: chromosome 18 (M18_B25_H08).

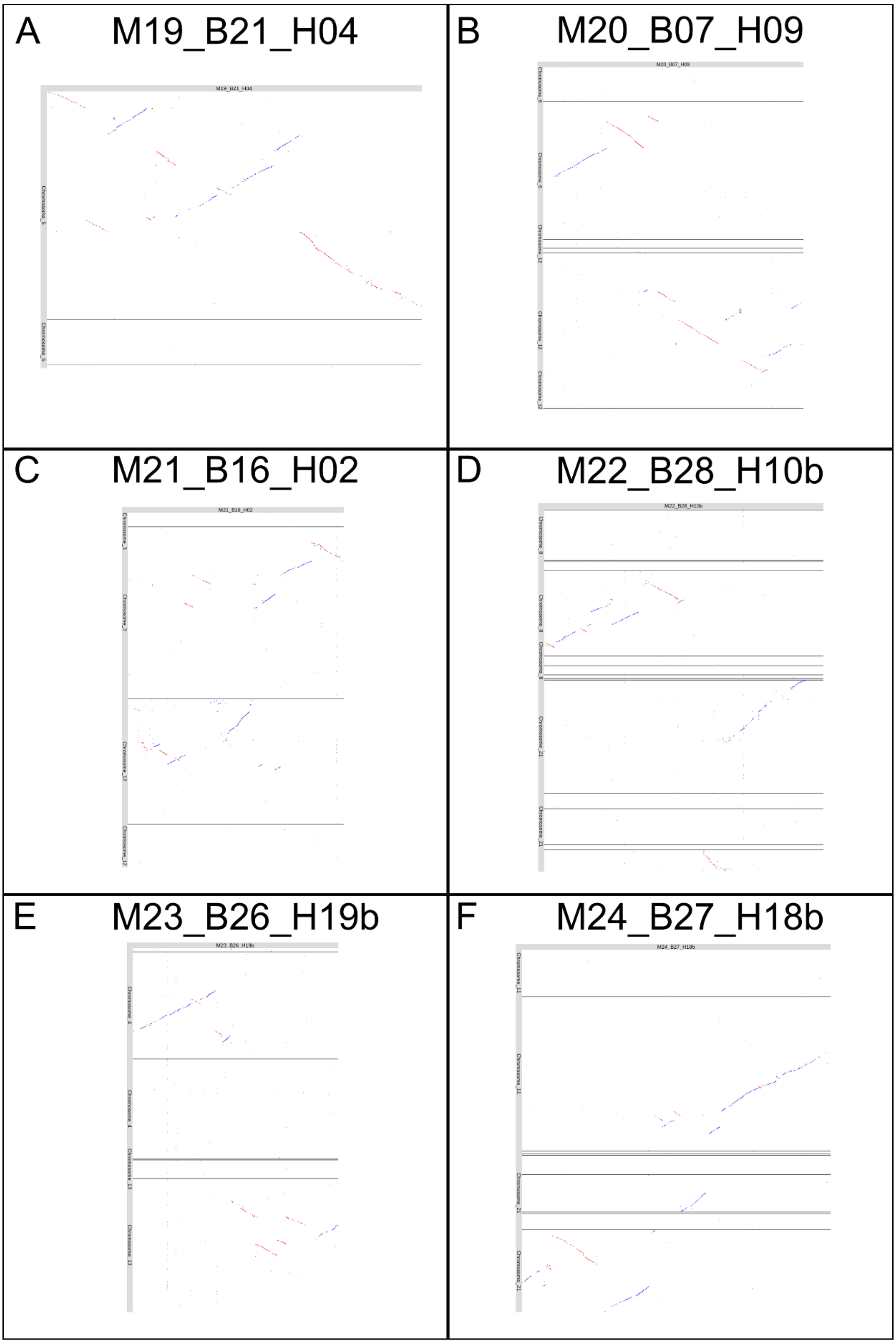


Figure S11: *M. cinxia* aligned against *P. napi* using the last aligner (Kielbasa et al. 2011). A: *M. cinxia* chromosome 19 (M19_B21_H04), B: chromosome 20 (M20_B07_H09), C: chromosome 21 (M21_B16_H02), D: chromosome 22 (M22_B28_H10b), E: chromosome 23 (M23_B26_H19b), and F: chromosome 24 (M24_B27_H18b).

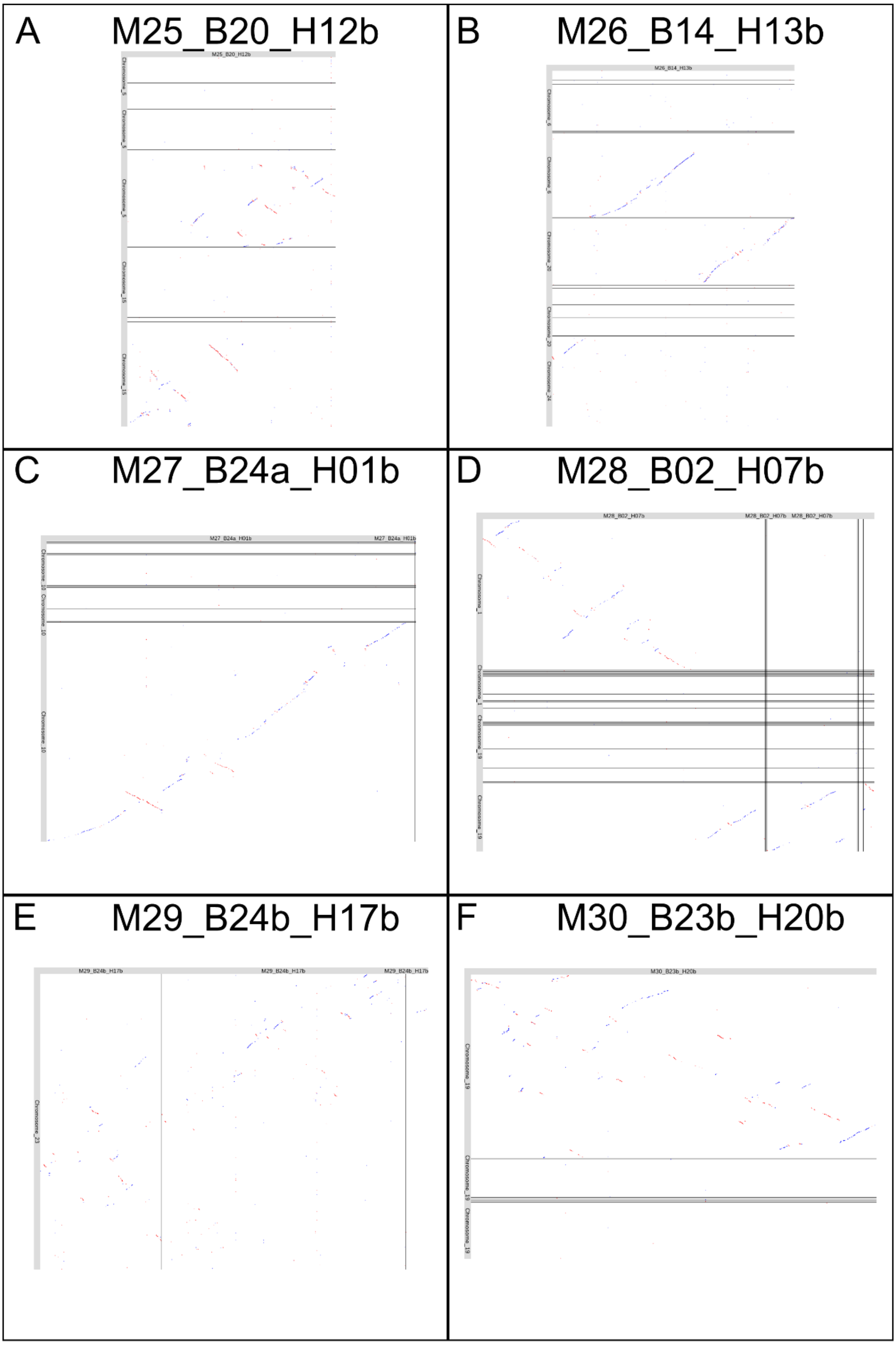


Figure S12: *M. cinxia* aligned against *P. napi* using the last aligner (Kielbasa et al. 2011). A: *M. cinxia* chromosome 25 (M25_B20_H12b), B: chromosome 26 (M26_B14_H13b), C: chromosome 27 (M27_B24a_H01b), D: chromosome 28 (M28_B02_H07b), E: chromosome 29 (M29_B24b_H17b), and F: chromosome 30 (M30_B23b_H20b).

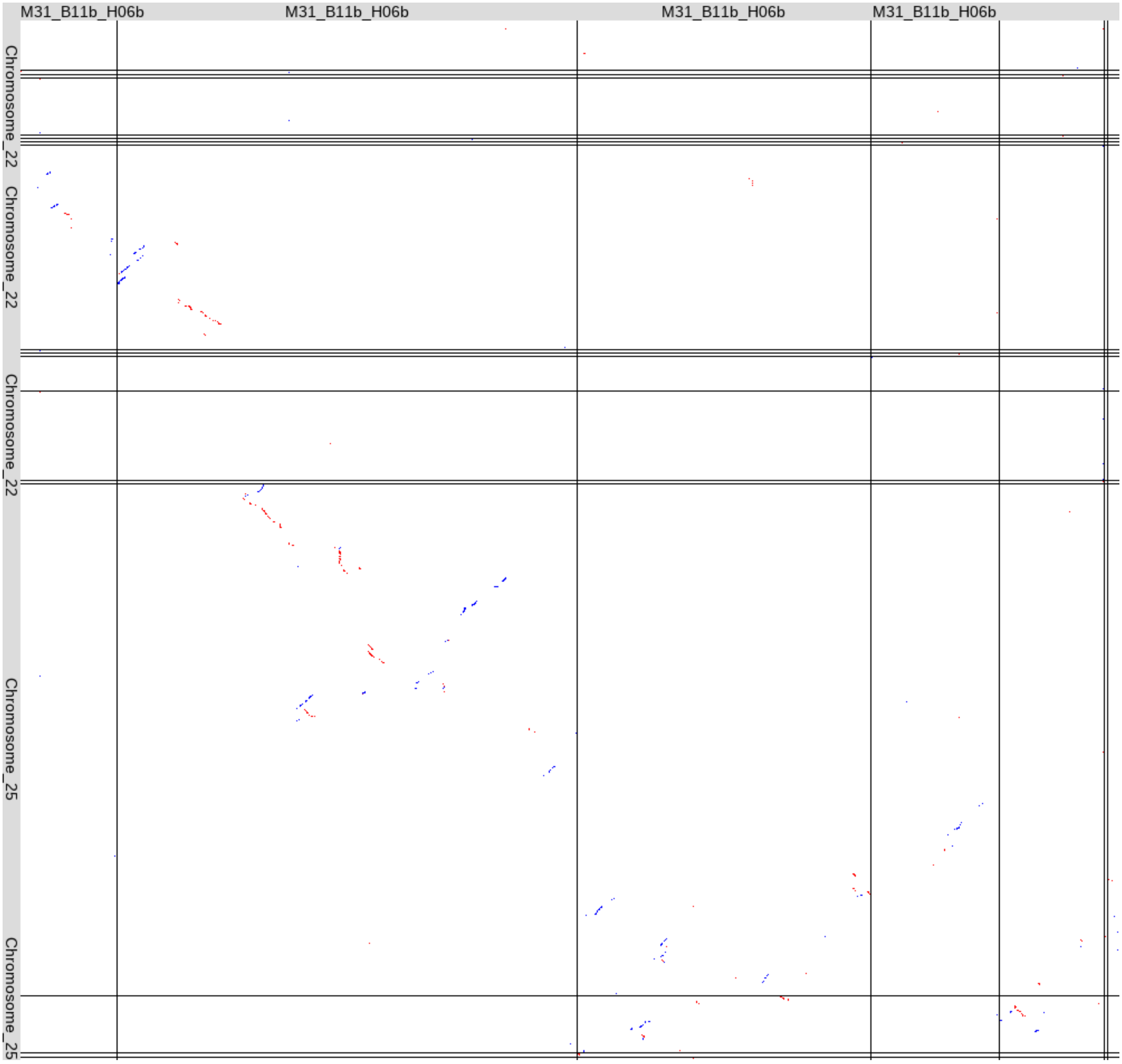


Figure S13: *M. cinxia* aligned against *P. napi* using the last aligner (Kielbasa et al. 2011). *M. cinxia* chromosome 31 (M31_B11b_H06b).

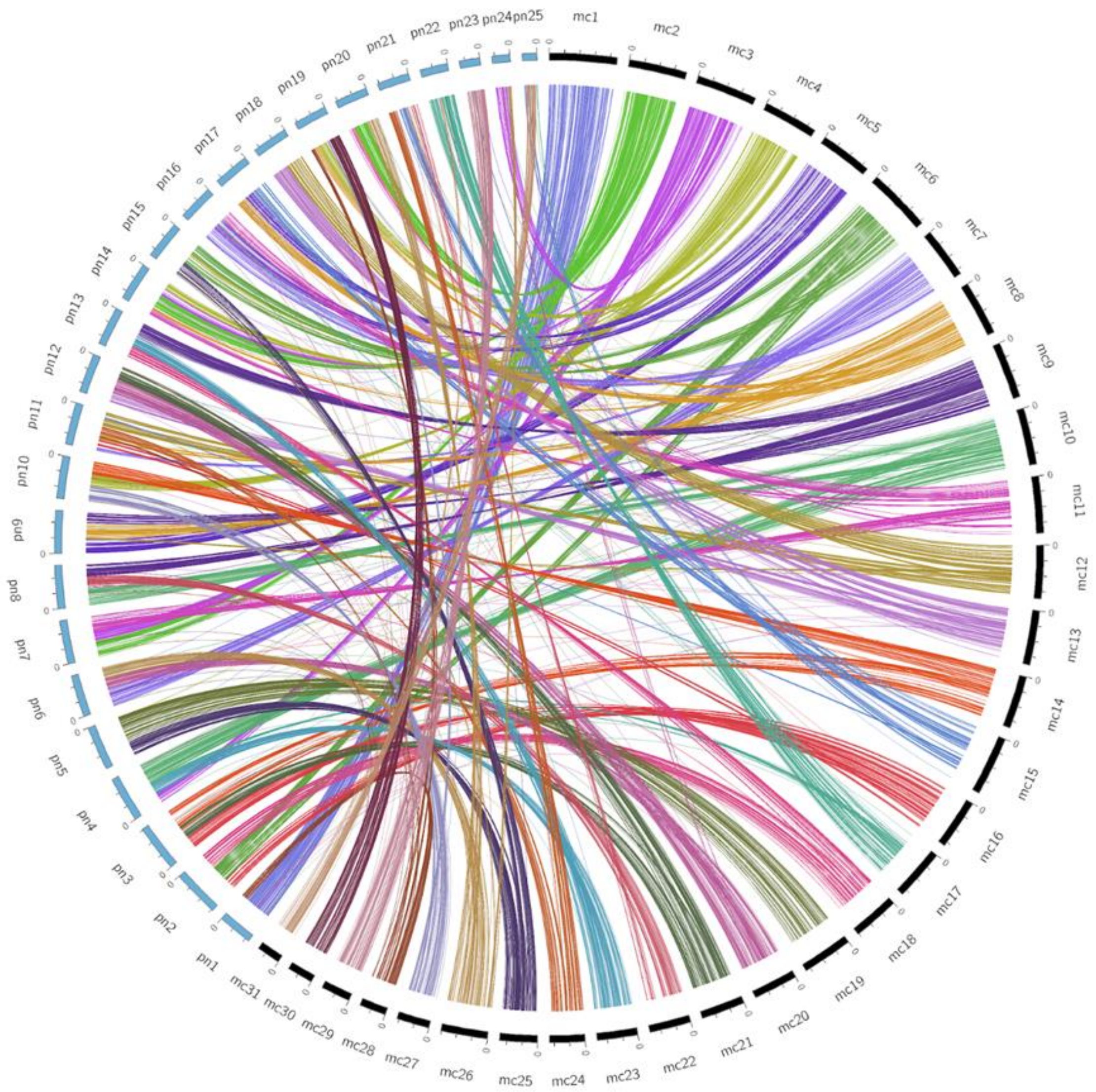


Figure S14: Orthologs between *M. cinxia* and *P. napi* were identified using OrthoFinder and filtered for one-to-one orthologs. The internal links in the circos plot indicate the orthologs between *M. cinxia* and *P. napi*. The links are coloured according to the *M. cinxia* chromosome.

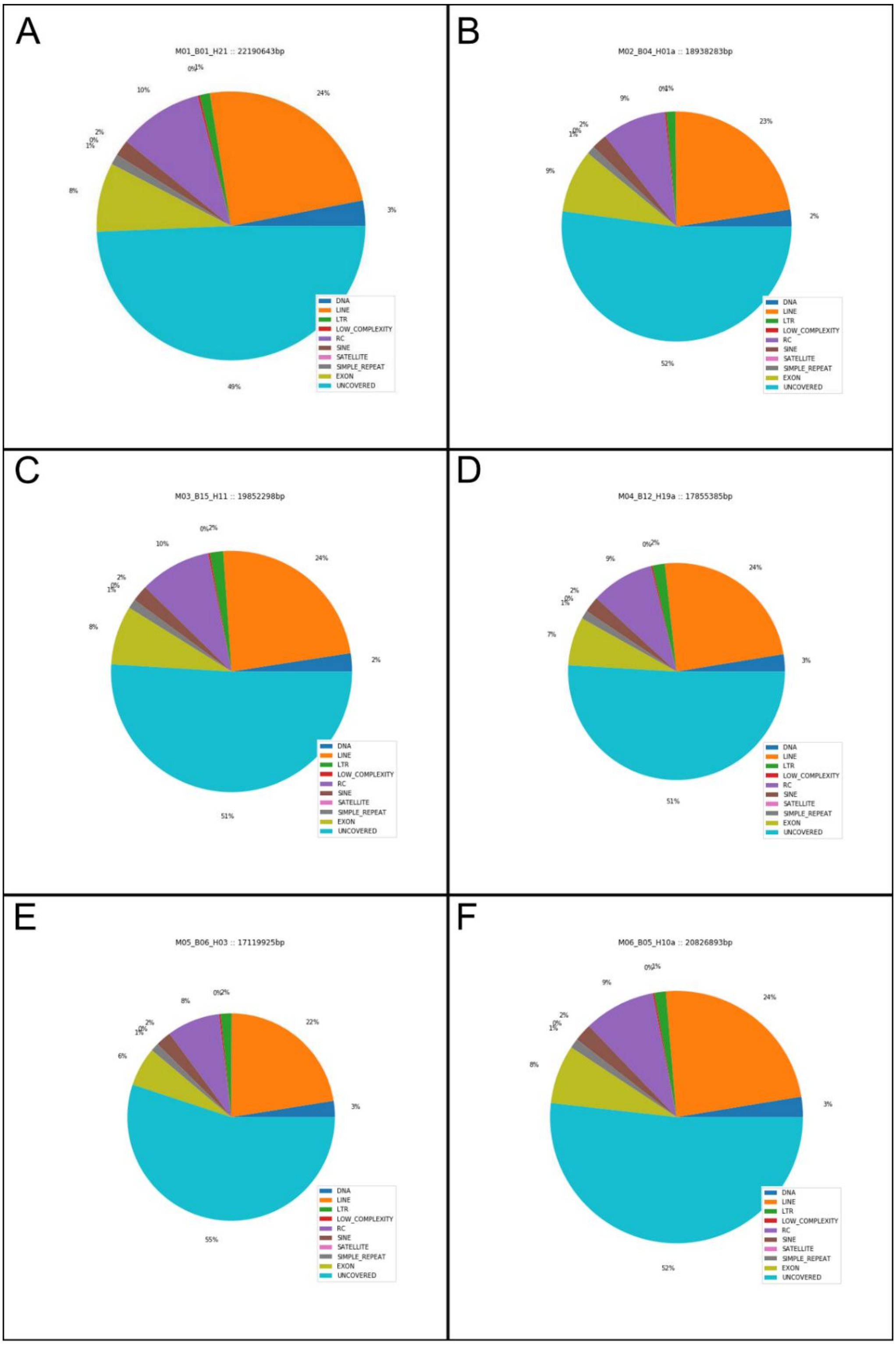


Figure S15: Repeat classes and coverage of the *M. cinxia* genome v.2. A: M. cinxia chromosome 1 (M01_B01_H21), B: chromosome 2 (M02_B04_H01a), C: chromosome 3 (M03_B15_H11), D: chromosome 4 (M04_B12_H19a), E: chromosome 5 (M05_B06_H03), and F: chromosome 6 (M06_B05_H10a). (DNA = class II; LINE = Long interspersed elements; LTR = Long terminal repeats; Low_complexity = Low complexity repeated DNA; RC = Rolling circle elements (e.g. Helitrons); SINE = Short interspersed elements; Satellite = Satellite DNA; Simple_repeat = Simple repeated motifs; Exon = exonic regions; Uncovered = rest of the chromosome).

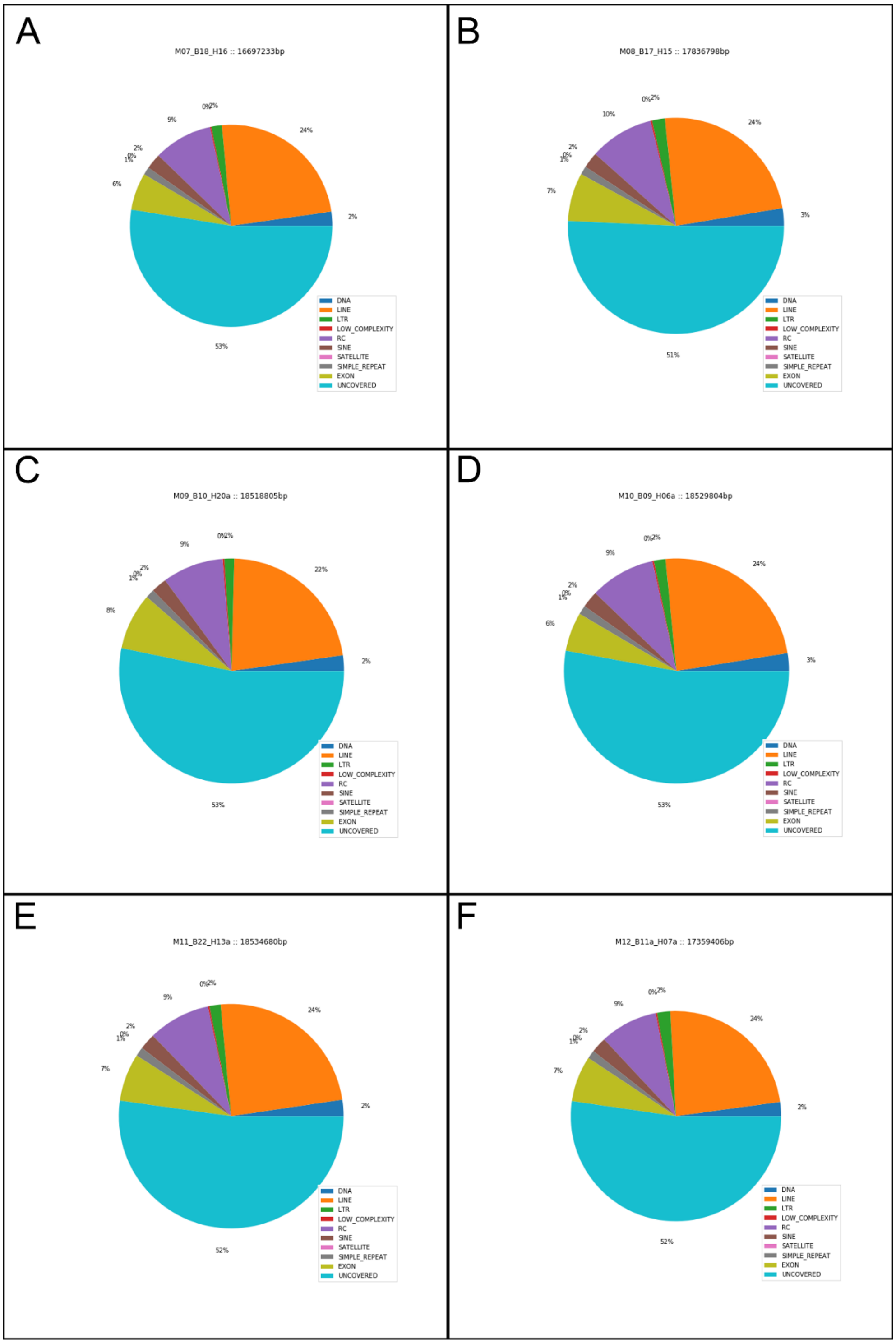


Figure S16: Repeat classes and coverage of the *M. cinxia* genome v.2. A: *M. cinxia* chromosome 7 (M07_B18_H16), B: chromosome 8 (M08_B17_H15), C: chromosome 9 (M09_B10_H20a), D: chromosome 10 (M10_B09_H06a), E: chromosome 11 (M11_B22_H13a), and F: chromosome 12 (M12_B11a_H07a). (DNA = class II; LINE = Long interspersed elements; LTR = Long terminal repeats; Low_complexity = Low complexity repeated DNA; RC = Rolling circle elements (e.g. Helitrons); SINE = Short interspersed elements; Satellite = Satellite DNA; Simple_repeat = Simple repeated motifs; Exon = exonic regions; Uncovered = rest of the chromosome).

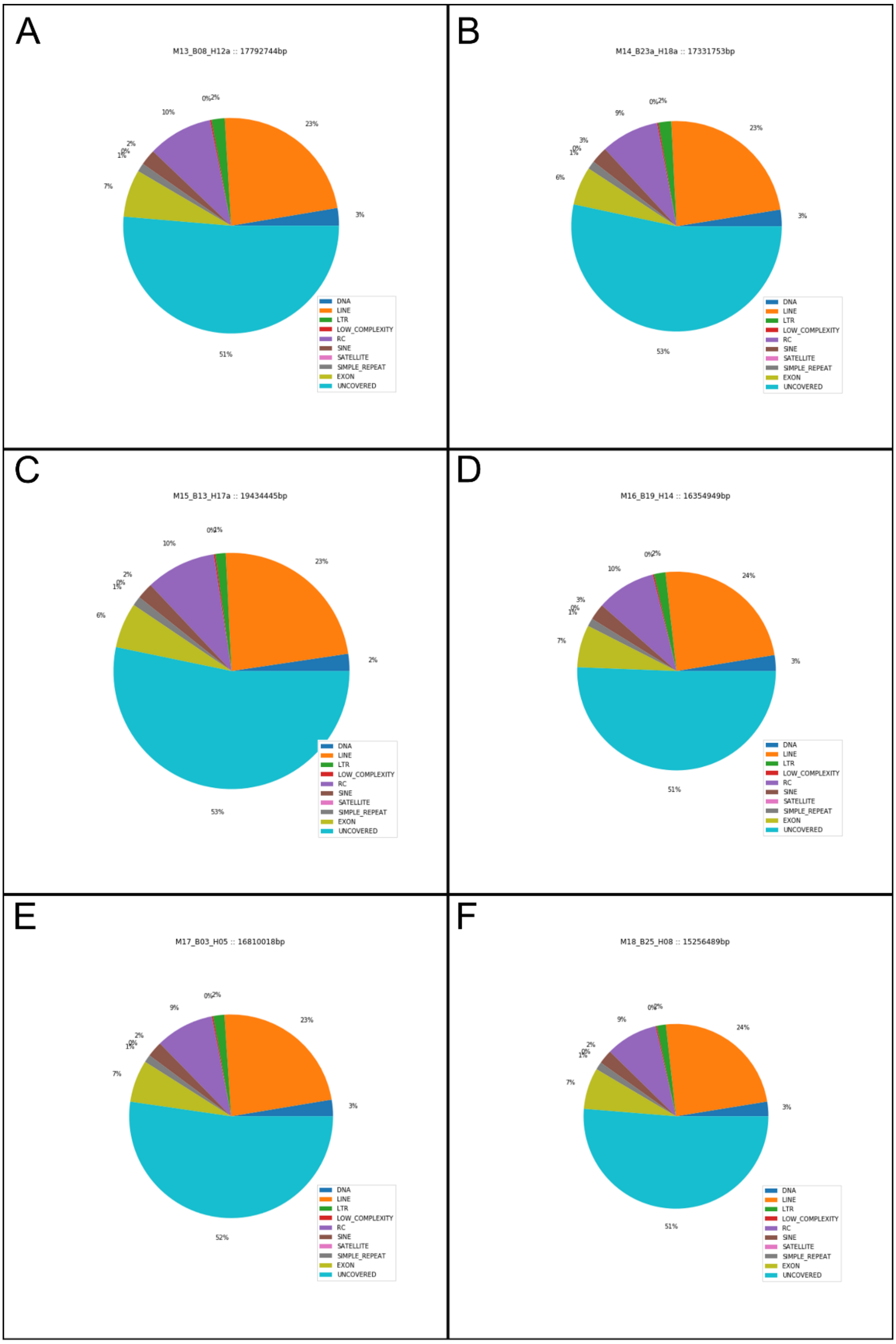


Figure S17: Repeat classes and coverage of the *M. cinxia* genome v.2. A: *M. cinxia* chromosome 13 (M13_B08_H12a), B: chromosome 14 (M14_B23a_H18a), C: chromosome 15 (M15_B13_H17a), D: chromosome 16 (M16_B19_H14), E: chromosome 17 (M17_B03_H05), and F: chromosome 18 (M18_B25_H08). (DNA = class II; LINE = Long interspersed elements; LTR = Long terminal repeats; Low_complexity = Low complexity repeated DNA; RC = Rolling circle elements (e.g. Helitrons); SINE = Short interspersed elements; Satellite = Satellite DNA; Simple_repeat = Simple repeated motifs; Exon = exonic regions; Uncovered = rest of the chromosome).

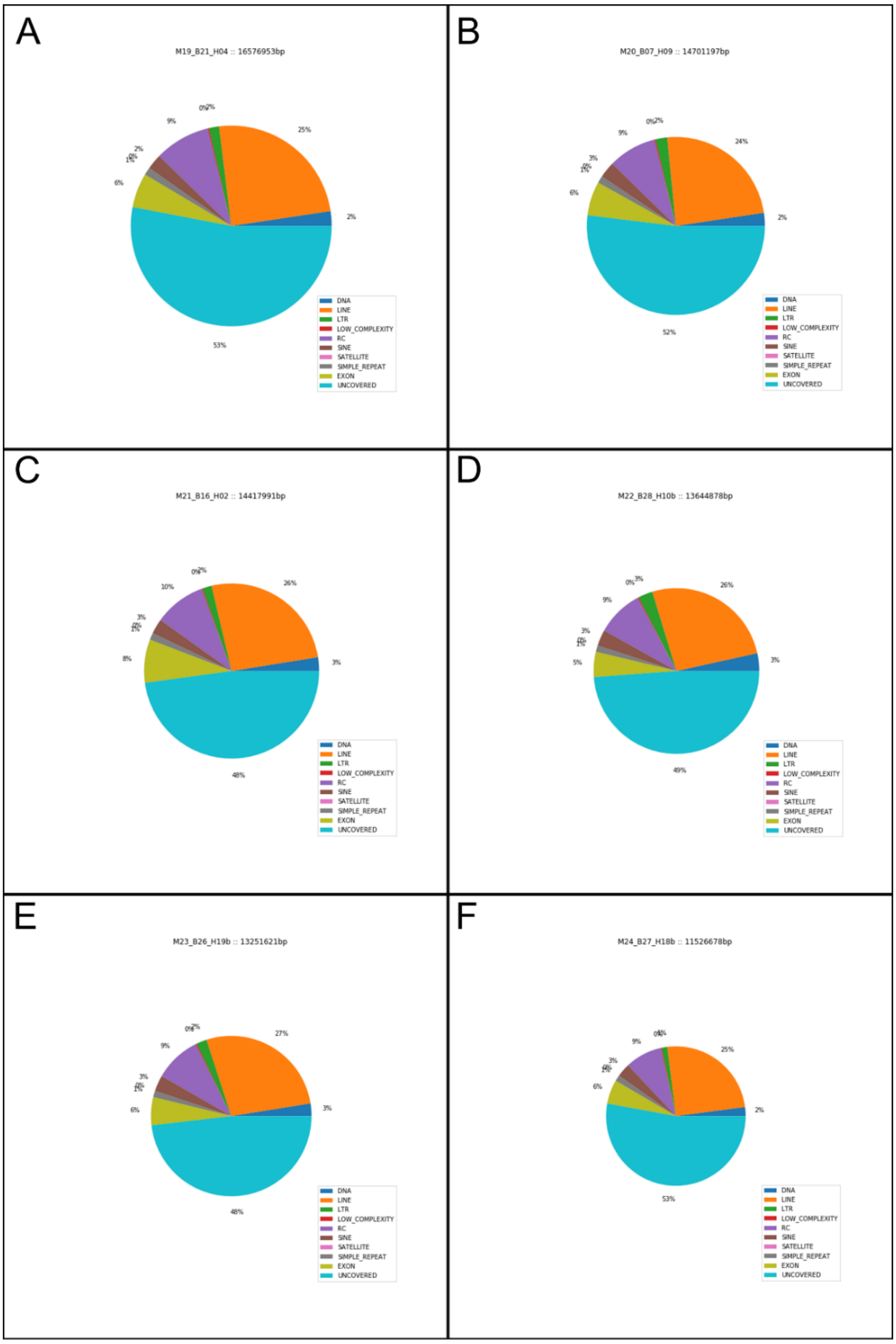


Figure S18: Repeat classes and coverage of the *M. cinxia* genome v.2. A: *M. cinxia* chromosome 19 (M19_B21_H04), B: chromosome 20 (M20_B07_H09), C: chromosome 21 (M21_B16_H02), D: chromosome 22 (M22_B28_H10b), E: chromosome 23 (M23_B26_H19b), and F: chromosome 24 (M24_B27_H18b). (DNA = class II; LINE = Long interspersed elements; LTR = Long terminal repeats; Low_complexity = Low complexity repeated DNA; RC = Rolling circle elements (e.g. Helitrons); SINE = Short interspersed elements; Satellite = Satellite DNA; Simple_repeat = Simple repeated motifs; Exon = exonic regions; Uncovered = rest of the chromosome).

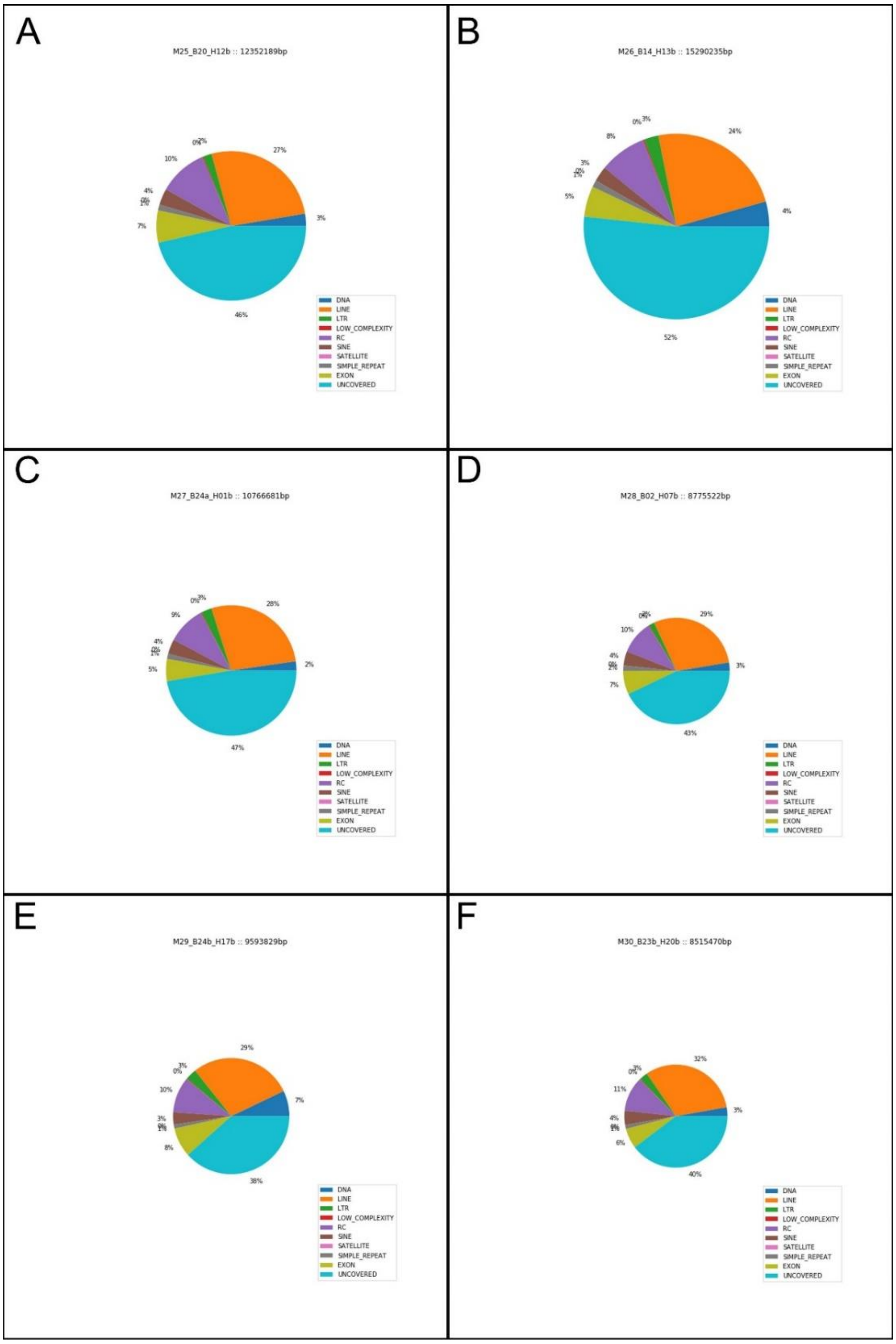


Figure S19: Repeat classes and coverage of the *M. cinxia* genome v.2. A: *M. cinxia* chromosome 25 (M25_B20_H12b), B: chromosome 26 (M26_B14_H13b), C: chromosome 27 (M27_B24a_H01b), D: chromosome 28 (M28_B02_H07b), E: chromosome 29 (M29_B24b_H17b), and F: chromosome 30 (M30_B23b_H20b). (DNA = class II; LINE = Long interspersed elements; LTR = Long terminal repeats; Low_complexity = Low complexity repeated DNA; RC = Rolling circle elements (e.g. Helitrons); SINE = Short interspersed elements; Satellite = Satellite DNA; Simple_repeat = Simple repeated motifs; Exon = exonic regions; Uncovered = rest of the chromosome).

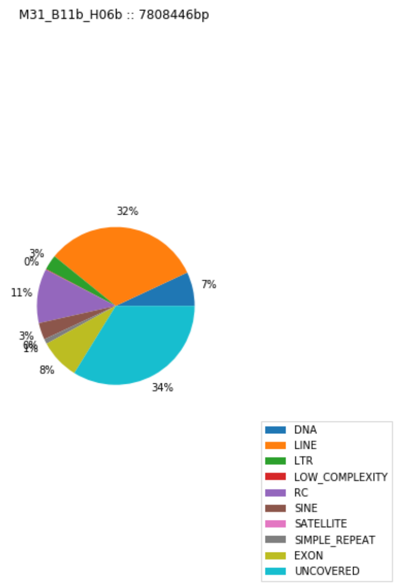


Figure S20: Repeat classes and coverage of the *M. cinxia* genome v.2. *M. cinxia* chromosome 31 (M31_B11b_H06b). (DNA = class II; LINE = Long interspersed elements; LTR = Long terminal repeats; Low_complexity = Low complexity repeated DNA; RC = Rolling circle elements (e.g. Helitrons); SINE = Short interspersed elements; Satellite = Satellite DNA; Simple_repeat = Simple repeated motifs; Exon = exonic regions; Uncovered = rest of the chromosome).
