## Supplementary Table 1 for "Improved chromosome level genome assembly of the Glanville fritillary butterfly (*Melitaea cinxia*) based on SMRT Sequencing and linkage map"

| | chromosome | genus | superfamily | coverage | perc_coverage | count |
|---|---|---|---|---|---|---|
| 0 | M01_B01_H21 | DNA | Academ-1 | 5007 | 0.022563564 | 18 |
| 1 | M01_B01_H21 | DNA | CMC-Chapaev | 944 | 0.004254045 | 2 |
| 2 | M01_B01_H21 | DNA | CMC-Transib | 24963 | 0.11249336 | 57 |
| 3 | M01_B01_H21 | DNA | Ginger-2 | 4685 | 0.021112502 | 4 |
| 4 | M01_B01_H21 | DNA | Maveric_Unknown | 2394 | 0.010788331 | 10 |
| 5 | M01_B01_H21 | DNA | P | 2383 | 0.01073876 | 5 |
| 6 | M01_B01_H21 | DNA | PIF-Harbing | 14445 | 0.065095004 | 66 |
| 7 | M01_B01_H21 | DNA | PIF-Spy | 14907 | 0.067176963 | 74 |
| 8 | M01_B01_H21 | DNA | PiggyBac | 27739 | 0.125003138 | 63 |
| 9 | M01_B01_H21 | DNA | Sola-1 | 58164 | 0.262110476 | 186 |
| 10 | M01_B01_H21 | DNA | Sola-2 | 18584 | 0.08374701 | 46 |
| 11 | M01_B01_H21 | DNA | TcMar-Fot1 | 34345 | 0.154772442 | 149 |
| 12 | M01_B01_H21 | DNA | TcMar-Marin | 221656 | 0.998871461 | 933 |
| 13 | M01_B01_H21 | DNA | TcMar-Tc1 | 202941 | 0.914534112 | 570 |
| 14 | M01_B01_H21 | DNA | TcMar-m44 | 18128 | 0.08169209 | 99 |
| 15 | M01_B01_H21 | DNA | Zator | 1140 | 0.0051373 | 9 |
| 16 | M01_B01_H21 | DNA | hAT-Ac | 22204 | 0.100060192 | 47 |
| 17 | M01_B01_H21 | DNA | hAT-Tip100 | 5640 | 0.025416118 | 11 |
| 18 | M01_B01_H21 | DNA | hAT-hATm | 67 | 0.000301929 | 1 |
| 19 | M01_B01_H21 | DNA | hAT-hATx | 236 | 0.001063511 | 2 |
| 20 | M01_B01_H21 | LINE | CR1 | 252340 | 1.137145958 | 632 |
| 21 | M01_B01_H21 | LINE | CR1-Zenon | 165460 | 0.745629588 | 670 |
| 22 | M01_B01_H21 | LINE | CRE | 102058 | 0.459914568 | 503 |
| 23 | M01_B01_H21 | LINE | Dong-R4 | 108029 | 0.486822306 | 281 |
| 24 | M01_B01_H21 | LINE | I | 163877 | 0.738495951 | 370 |
| 25 | M01_B01_H21 | LINE | I-Jockey | 38066 | 0.171540771 | 91 |
| 26 | M01_B01_H21 | LINE | L1 | 11867 | 0.053477495 | 139 |
| 27 | M01_B01_H21 | LINE | L1-Tx1 | 19285 | 0.086905999 | 64 |
| 28 | M01_B01_H21 | LINE | L2 | 567478 | 2.557285068 | 2087 |
| 29 | M01_B01_H21 | LINE | Penelope | 24395 | 0.109933723 | 184 |
| 30 | M01_B01_H21 | LINE | Proto2 | 66644 | 0.300324781 | 173 |
| 31 | M01_B01_H21 | LINE | R1 | 260069 | 1.171975954 | 542 |
| 32 | M01_B01_H21 | LINE | R1-LOA | 54463 | 0.245432275 | 83 |
| 33 | M01_B01_H21 | LINE | R2 | 202 | 0.000910294 | 2 |
| 34 | M01_B01_H21 | LINE | RTE | 29535 | 0.133096639 | 213 |
| 35 | M01_B01_H21 | LINE | RTE-BovB | 177544 | 0.800084973 | 661 |
| 36 | M01_B01_H21 | LINE | RTE-RTE | 581502 | 2.620482876 | 2539 |
| 37 | M01_B01_H21 | LINE | RTE-X | 25572 | 0.11523776 | 41 |
| 38 | M01_B01_H21 | LINE | Tad1 | 622 | 0.002802983 | 1 |
| 39 | M01_B01_H21 | LINE | Unknown | 2776886 | 12.51376988 | 14956 |
| 40 | M01_B01_H21 | LTR | Copia | 235 | 0.001059005 | 1 |
| 41 | M01_B01_H21 | LTR | DIRS | 555 | 0.002501054 | 2 |
| 42 | M01_B01_H21 | LTR | Gypsy | 199230 | 0.897810848 | 279 |
| 43 | M01_B01_H21 | LTR | Pao | 74368 | 0.335132245 | 88 |
| 44 | M01_B01_H21 | RC | Helitron | 2259561 | 10.18249449 | 10766 |
| 45 | M01_B01_H21 | SINE | SINE | 436110 | 1.965287802 | 2449 |

| | | | | | | |
|---|---|---|---|---|---|---|
| 46 | M02_B04_H01a | DNA | Academ-1 | 1686 | 0.008902602 | 8 |
| 47 | M02_B04_H01a | DNA | CMC-Transib | 27370 | 0.144522077 | 69 |
| 48 | M02_B04_H01a | DNA | Ginger-2 | 3263 | 0.017229651 | 2 |
| 49 | M02_B04_H01a | DNA | Kolobok-Hyd | 61 | 0.000322099 | 1 |
| 50 | M02_B04_H01a | DNA | MULE-MuDR | 1810 | 0.009557361 | 5 |
| 51 | M02_B04_H01a | DNA | P | 1379 | 0.007281547 | 3 |
| 52 | M02_B04_H01a | DNA | PIF-Harbing | 13327 | 0.070370688 | 51 |
| 53 | M02_B04_H01a | DNA | PIF-Spy | 8140 | 0.042981721 | 61 |
| 54 | M02_B04_H01a | DNA | PiggyBac | 32880 | 0.173616584 | 57 |
| 55 | M02_B04_H01a | DNA | Sola-1 | 28756 | 0.151840587 | 125 |
| 56 | M02_B04_H01a | DNA | Sola-2 | 8246 | 0.043541434 | 27 |
| 57 | M02_B04_H01a | DNA | TcMar-Fot1 | 22066 | 0.116515315 | 157 |
| 58 | M02_B04_H01a | DNA | TcMar-Marin | 141742 | 0.748441662 | 714 |
| 59 | M02_B04_H01a | DNA | TcMar-Tc1 | 116810 | 0.61679298 | 385 |
| 60 | M02_B04_H01a | DNA | TcMar-m44 | 13638 | 0.072012864 | 70 |
| 61 | M02_B04_H01a | DNA | Zator | 1290 | 0.0068116 | 9 |
| 62 | M02_B04_H01a | DNA | hAT-Ac | 20561 | 0.108568448 | 30 |
| 63 | M02_B04_H01a | DNA | hAT-Tip100 | 3043 | 0.016067983 | 9 |
| 64 | M02_B04_H01a | DNA | hAT-hATm | 1419 | 0.00749276 | 4 |
| 65 | M02_B04_H01a | DNA | hAT-hATx | 497 | 0.002624314 | 5 |
| 66 | M02_B04_H01a | LINE | CR1 | 172653 | 0.911661316 | 411 |
| 67 | M02_B04_H01a | LINE | CR1-Zenon | 138422 | 0.730911033 | 529 |
| 68 | M02_B04_H01a | LINE | CRE | 63876 | 0.337285064 | 337 |
| 69 | M02_B04_H01a | LINE | Dong-R4 | 73979 | 0.390632034 | 158 |
| 70 | M02_B04_H01a | LINE | I | 164169 | 0.866863168 | 353 |
| 71 | M02_B04_H01a | LINE | I-Jockey | 37354 | 0.19724069 | 74 |
| 72 | M02_B04_H01a | LINE | L1 | 12433 | 0.065650091 | 137 |
| 73 | M02_B04_H01a | LINE | L1-Tx1 | 8713 | 0.046007339 | 33 |
| 74 | M02_B04_H01a | LINE | L2 | 478948 | 2.528993785 | 1778 |
| 75 | M02_B04_H01a | LINE | Penelope | 23930 | 0.126357812 | 180 |
| 76 | M02_B04_H01a | LINE | Proto2 | 43981 | 0.232233302 | 125 |
| 77 | M02_B04_H01a | LINE | R1 | 152335 | 0.804375983 | 370 |
| 78 | M02_B04_H01a | LINE | R1-LOA | 44576 | 0.235375087 | 58 |
| 79 | M02_B04_H01a | LINE | R2 | 148 | 0.000781486 | 2 |
| 80 | M02_B04_H01a | LINE | RTE | 23857 | 0.125972349 | 180 |
| 81 | M02_B04_H01a | LINE | RTE-BovB | 148781 | 0.785609762 | 437 |
| 82 | M02_B04_H01a | LINE | RTE-RTE | 419747 | 2.216394168 | 1926 |
| 83 | M02_B04_H01a | LINE | RTE-X | 20846 | 0.110073337 | 29 |
| 84 | M02_B04_H01a | LINE | Tad1 | 107 | 0.000564993 | 2 |
| 85 | M02_B04_H01a | LINE | Unknown | 2290313 | 12.09356202 | 12601 |
| 86 | M02_B04_H01a | LTR | Copia | 78 | 0.000411864 | 1 |
| 87 | M02_B04_H01a | LTR | Gypsy | 151544 | 0.800199258 | 215 |
| 88 | M02_B04_H01a | LTR | Pao | 84024 | 0.443672745 | 93 |
| 89 | M02_B04_H01a | RC | Helitron | 1707489 | 9.016070781 | 7908 |
| 90 | M02_B04_H01a | SINE | SINE | 417157 | 2.202718166 | 2310 |
| 91 | M03_B15_H11 | DNA | Academ-1 | 1160 | 0.005843152 | 7 |
| 92 | M03_B15_H11 | DNA | CMC-Chapaev | 635 | 0.003198622 | 2 |

| | | | | | | |
|---|---|---|---|---|---|---|
| **93** | M03_B15_H11 | DNA | CMC-Transib | 13911 | 0.070072492 | 50 |
| **94** | M03_B15_H11 | DNA | Ginger-2 | 3435 | 0.017302783 | 6 |
| **95** | M03_B15_H11 | DNA | MULE-MuDR | 168 | 0.00084625 | 1 |
| **96** | M03_B15_H11 | DNA | P | 1030 | 0.005188316 | 4 |
| **97** | M03_B15_H11 | DNA | PIF-Harbing | 11613 | 0.058497006 | 25 |
| **98** | M03_B15_H11 | DNA | PIF-Spy | 10955 | 0.055182528 | 76 |
| **99** | M03_B15_H11 | DNA | PiggyBac | 26381 | 0.132886379 | 54 |
| **100** | M03_B15_H11 | DNA | Sola-1 | 39116 | 0.197035124 | 146 |
| **101** | M03_B15_H11 | DNA | Sola-2 | 16100 | 0.081098924 | 27 |
| **102** | M03_B15_H11 | DNA | TcMar-Fot1 | 31828 | 0.160324009 | 129 |
| **103** | M03_B15_H11 | DNA | TcMar-Marin | 143184 | 0.721246477 | 755 |
| **104** | M03_B15_H11 | DNA | TcMar-Tc1 | 142089 | 0.715730743 | 419 |
| **105** | M03_B15_H11 | DNA | TcMar-m44 | 19437 | 0.097908061 | 87 |
| **106** | M03_B15_H11 | DNA | Zator | 3412 | 0.017186927 | 16 |
| **107** | M03_B15_H11 | DNA | hAT-Ac | 16901 | 0.085133721 | 32 |
| **108** | M03_B15_H11 | DNA | hAT-Tip100 | 1785 | 0.008991402 | 9 |
| **109** | M03_B15_H11 | DNA | hAT-hATx | 2871 | 0.014461802 | 2 |
| **110** | M03_B15_H11 | LINE | CR1 | 181728 | 0.915400323 | 431 |
| **111** | M03_B15_H11 | LINE | CR1-Zenon | 144095 | 0.725835367 | 571 |
| **112** | M03_B15_H11 | LINE | CRE | 82809 | 0.417125514 | 375 |
| **113** | M03_B15_H11 | LINE | Dong-R4 | 63570 | 0.320214818 | 191 |
| **114** | M03_B15_H11 | LINE | I | 181313 | 0.913309885 | 438 |
| **115** | M03_B15_H11 | LINE | I-Jockey | 31142 | 0.156868489 | 66 |
| **116** | M03_B15_H11 | LINE | L1 | 12998 | 0.065473529 | 142 |
| **117** | M03_B15_H11 | LINE | L1-Tx1 | 6593 | 0.033210261 | 33 |
| **118** | M03_B15_H11 | LINE | L2 | 461154 | 2.322925034 | 1777 |
| **119** | M03_B15_H11 | LINE | Penelope | 24464 | 0.123230066 | 177 |
| **120** | M03_B15_H11 | LINE | Proto2 | 43577 | 0.219506074 | 127 |
| **121** | M03_B15_H11 | LINE | R1 | 215029 | 1.083144128 | 399 |
| **122** | M03_B15_H11 | LINE | R1-LOA | 72967 | 0.367549389 | 77 |
| **123** | M03_B15_H11 | LINE | R2 | 10 | 5.0372E-05 | 1 |
| **124** | M03_B15_H11 | LINE | R2-NeSL | 805 | 0.004054946 | 1 |
| **125** | M03_B15_H11 | LINE | RTE | 28238 | 0.14224046 | 193 |
| **126** | M03_B15_H11 | LINE | RTE-BovB | 159961 | 0.805755586 | 524 |
| **127** | M03_B15_H11 | LINE | RTE-RTE | 466355 | 2.349123512 | 2003 |
| **128** | M03_B15_H11 | LINE | RTE-X | 21168 | 0.106627454 | 28 |
| **129** | M03_B15_H11 | LINE | Unknown | 2499616 | 12.59106628 | 13661 |
| **130** | M03_B15_H11 | LTR | Copia | 5102 | 0.025699796 | 4 |
| **131** | M03_B15_H11 | LTR | DIRS | 7777 | 0.039174306 | 5 |
| **132** | M03_B15_H11 | LTR | Gypsy | 228247 | 1.14972584 | 267 |
| **133** | M03_B15_H11 | LTR | Pao | 116244 | 0.585544303 | 102 |
| **134** | M03_B15_H11 | RC | Helitron | 1900152 | 9.571446086 | 8702 |
| **135** | M03_B15_H11 | SINE | SINE | 436673 | 2.199609335 | 2464 |
| **136** | M04_B12_H19a | DNA | Academ-1 | 2532 | 0.014180596 | 6 |
| **137** | M04_B12_H19a | DNA | CMC-Transib | 11037 | 0.061813285 | 55 |
| **138** | M04_B12_H19a | DNA | Ginger-2 | 6346 | 0.035541099 | 9 |
| **139** | M04_B12_H19a | DNA | Maveric_Unknown | 146 | 0.00081768 | 2 |

| | | | | | | |
|---|---|---|---|---|---|---|
| **140** | M04_B12_H19a | DNA | P | 640 | 0.003584353 | 3 |
| **141** | M04_B12_H19a | DNA | PIF-Harbing | 4236 | 0.023723935 | 32 |
| **142** | M04_B12_H19a | DNA | PIF-Spy | 13499 | 0.075601842 | 78 |
| **143** | M04_B12_H19a | DNA | PiggyBac | 37170 | 0.208172493 | 64 |
| **144** | M04_B12_H19a | DNA | Sola-1 | 50536 | 0.283029461 | 129 |
| **145** | M04_B12_H19a | DNA | Sola-2 | 5265 | 0.029486903 | 25 |
| **146** | M04_B12_H19a | DNA | TcMar-Fot1 | 25142 | 0.140809061 | 113 |
| **147** | M04_B12_H19a | DNA | TcMar-Marin | 150872 | 0.844966378 | 754 |
| **148** | M04_B12_H19a | DNA | TcMar-Tc1 | 113536 | 0.635864194 | 337 |
| **149** | M04_B12_H19a | DNA | TcMar-m44 | 13500 | 0.075607443 | 49 |
| **150** | M04_B12_H19a | DNA | Zator | 316 | 0.001769774 | 2 |
| **151** | M04_B12_H19a | DNA | hAT-Ac | 18265 | 0.10229407 | 23 |
| **152** | M04_B12_H19a | DNA | hAT-Charlie | 196 | 0.001097708 | 1 |
| **153** | M04_B12_H19a | DNA | hAT-Tip100 | 5418 | 0.030343787 | 5 |
| **154** | M04_B12_H19a | DNA | hAT-hATm | 115 | 0.000644063 | 1 |
| **155** | M04_B12_H19a | LINE | CR1 | 174169 | 0.975442423 | 452 |
| **156** | M04_B12_H19a | LINE | CR1-Zenon | 141404 | 0.791940359 | 508 |
| **157** | M04_B12_H19a | LINE | CRE | 72159 | 0.404130183 | 339 |
| **158** | M04_B12_H19a | LINE | Dong-R4 | 54118 | 0.303090636 | 149 |
| **159** | M04_B12_H19a | LINE | I | 101987 | 0.571183427 | 352 |
| **160** | M04_B12_H19a | LINE | I-Jockey | 20275 | 0.113551178 | 52 |
| **161** | M04_B12_H19a | LINE | L1 | 13052 | 0.073098396 | 147 |
| **162** | M04_B12_H19a | LINE | L1-Tx1 | 6166 | 0.034532999 | 28 |
| **163** | M04_B12_H19a | LINE | L2 | 504727 | 2.826749465 | 1765 |
| **164** | M04_B12_H19a | LINE | Penelope | 26532 | 0.148593828 | 190 |
| **165** | M04_B12_H19a | LINE | Proto2 | 36004 | 0.20164225 | 118 |
| **166** | M04_B12_H19a | LINE | R1 | 181272 | 1.015223139 | 379 |
| **167** | M04_B12_H19a | LINE | R1-LOA | 52785 | 0.295625101 | 54 |
| **168** | M04_B12_H19a | LINE | RTE | 27243 | 0.15257582 | 195 |
| **169** | M04_B12_H19a | LINE | RTE-BovB | 166453 | 0.932228569 | 478 |
| **170** | M04_B12_H19a | LINE | RTE-RTE | 436358 | 2.443845372 | 1916 |
| **171** | M04_B12_H19a | LINE | RTE-X | 19688 | 0.110263654 | 28 |
| **172** | M04_B12_H19a | LINE | Unknown | 2280132 | 12.76999628 | 12579 |
| **173** | M04_B12_H19a | LTR | Copia | 4810 | 0.026938652 | 3 |
| **174** | M04_B12_H19a | LTR | DIRS | 6251 | 0.035009046 | 7 |
| **175** | M04_B12_H19a | LTR | Gypsy | 184617 | 1.033956983 | 227 |
| **176** | M04_B12_H19a | LTR | Pao | 134455 | 0.753022127 | 102 |
| **177** | M04_B12_H19a | RC | Helitron | 1677793 | 9.396565798 | 7831 |
| **178** | M04_B12_H19a | SINE | SINE | 426767 | 2.390130484 | 2423 |
| **179** | M05_B06_H03 | DNA | Academ-1 | 231 | 0.001349305 | 2 |
| **180** | M05_B06_H03 | DNA | CMC-Transib | 9830 | 0.057418476 | 60 |
| **181** | M05_B06_H03 | DNA | Ginger-2 | 1647 | 0.009620369 | 1 |
| **182** | M05_B06_H03 | DNA | Maveric_Unknown | 7231 | 0.042237335 | 2 |
| **183** | M05_B06_H03 | DNA | PIF-Harbing | 11962 | 0.069871801 | 46 |
| **184** | M05_B06_H03 | DNA | PIF-Spy | 6434 | 0.03758194 | 53 |
| **185** | M05_B06_H03 | DNA | PiggyBac | 48952 | 0.285935832 | 69 |
| **186** | M05_B06_H03 | DNA | Sola-1 | 48762 | 0.284826014 | 120 |

| | | | | | | |
|---|---|---|---|---|---|---|
| 187 | M05_B06_H03 | DNA | Sola-2 | 6593 | 0.038510683 | 15 |
| 188 | M05_B06_H03 | DNA | TcMar-Fot1 | 22323 | 0.130391926 | 146 |
| 189 | M05_B06_H03 | DNA | TcMar-Marin | 122212 | 0.713858267 | 660 |
| 190 | M05_B06_H03 | DNA | TcMar-Tc1 | 117342 | 0.685411881 | 335 |
| 191 | M05_B06_H03 | DNA | TcMar-m44 | 7525 | 0.043954632 | 46 |
| 192 | M05_B06_H03 | DNA | Zator | 387 | 0.002260524 | 6 |
| 193 | M05_B06_H03 | DNA | hAT-Ac | 18224 | 0.106449064 | 35 |
| 194 | M05_B06_H03 | DNA | hAT-Tip100 | 1100 | 0.006425262 | 8 |
| 195 | M05_B06_H03 | DNA | hAT-hATx | 163 | 0.000952107 | 2 |
| 196 | M05_B06_H03 | LINE | CR1 | 145550 | 0.850178958 | 395 |
| 197 | M05_B06_H03 | LINE | CR1-Zenon | 114669 | 0.669798495 | 498 |
| 198 | M05_B06_H03 | LINE | CRE | 68526 | 0.400270445 | 371 |
| 199 | M05_B06_H03 | LINE | Dong-R4 | 49426 | 0.288704536 | 144 |
| 200 | M05_B06_H03 | LINE | I | 90362 | 0.527817733 | 334 |
| 201 | M05_B06_H03 | LINE | I-Jockey | 37890 | 0.221321063 | 57 |
| 202 | M05_B06_H03 | LINE | L1 | 11292 | 0.065958233 | 125 |
| 203 | M05_B06_H03 | LINE | L1-Tx1 | 5215 | 0.030461582 | 33 |
| 204 | M05_B06_H03 | LINE | L2 | 419837 | 2.452329669 | 1579 |
| 205 | M05_B06_H03 | LINE | Penelope | 31032 | 0.181262476 | 189 |
| 206 | M05_B06_H03 | LINE | Proto2 | 35935 | 0.20990162 | 129 |
| 207 | M05_B06_H03 | LINE | R1 | 170178 | 0.994034729 | 375 |
| 208 | M05_B06_H03 | LINE | R1-LOA | 21001 | 0.12266993 | 41 |
| 209 | M05_B06_H03 | LINE | R2 | 266 | 0.001553745 | 2 |
| 210 | M05_B06_H03 | LINE | RTE | 25703 | 0.150135004 | 192 |
| 211 | M05_B06_H03 | LINE | RTE-BovB | 129177 | 0.754541857 | 489 |
| 212 | M05_B06_H03 | LINE | RTE-RTE | 378795 | 2.21259731 | 1778 |
| 213 | M05_B06_H03 | LINE | RTE-X | 20128 | 0.117570609 | 25 |
| 214 | M05_B06_H03 | LINE | Unknown | 2080627 | 12.15324833 | 11690 |
| 215 | M05_B06_H03 | LTR | Copia | 35 | 0.00020444 | 1 |
| 216 | M05_B06_H03 | LTR | DIRS | 3369 | 0.019678825 | 1 |
| 217 | M05_B06_H03 | LTR | Gypsy | 183155 | 1.069835294 | 219 |
| 218 | M05_B06_H03 | LTR | Pao | 105432 | 0.615843819 | 103 |
| 219 | M05_B06_H03 | RC | Helitron | 1413943 | 8.259049032 | 7006 |
| 220 | M05_B06_H03 | SINE | SINE | 417875 | 2.440869338 | 2386 |
| 221 | M06_B05_H10a | DNA | Academ-1 | 1456 | 0.006990961 | 8 |
| 222 | M06_B05_H10a | DNA | CMC-Transib | 16419 | 0.078835571 | 60 |
| 223 | M06_B05_H10a | DNA | Ginger-2 | 6539 | 0.031396906 | 4 |
| 224 | M06_B05_H10a | DNA | Maveric_Unknown | 70 | 0.000336104 | 1 |
| 225 | M06_B05_H10a | DNA | PIF-Harbing | 10105 | 0.048518999 | 62 |
| 226 | M06_B05_H10a | DNA | PIF-Spy | 10905 | 0.052360186 | 72 |
| 227 | M06_B05_H10a | DNA | PiggyBac | 28188 | 0.13534424 | 69 |
| 228 | M06_B05_H10a | DNA | Sola-1 | 43661 | 0.209637607 | 120 |
| 229 | M06_B05_H10a | DNA | Sola-2 | 10132 | 0.048648639 | 33 |
| 230 | M06_B05_H10a | DNA | TcMar-Fot1 | 32740 | 0.157200596 | 150 |
| 231 | M06_B05_H10a | DNA | TcMar-Marin | 182243 | 0.875036905 | 816 |
| 232 | M06_B05_H10a | DNA | TcMar-Tc1 | 162149 | 0.77855588 | 438 |
| 233 | M06_B05_H10a | DNA | TcMar-m44 | 10432 | 0.050089084 | 54 |

| | | | | | | |
|---|---|---|---|---|---|---|
| **234** | M06_B05_H10a | DNA | Zator | 872 | 0.004186894 | 8 |
| **235** | M06_B05_H10a | DNA | hAT-Ac | 22955 | 0.110218072 | 34 |
| **236** | M06_B05_H10a | DNA | hAT-Tip100 | 246 | 0.001181165 | 3 |
| **237** | M06_B05_H10a | DNA | hAT-hATx | 2793 | 0.013410546 | 1 |
| **238** | M06_B05_H10a | LINE | CR1 | 217821 | 1.045864114 | 471 |
| **239** | M06_B05_H10a | LINE | CR1-Zenon | 162093 | 0.778286997 | 592 |
| **240** | M06_B05_H10a | LINE | CRE | 94683 | 0.454618939 | 389 |
| **241** | M06_B05_H10a | LINE | Dong-R4 | 89178 | 0.428186768 | 181 |
| **242** | M06_B05_H10a | LINE | I | 147238 | 0.706960947 | 417 |
| **243** | M06_B05_H10a | LINE | I-Jockey | 31174 | 0.149681472 | 78 |
| **244** | M06_B05_H10a | LINE | L1 | 11369 | 0.054588075 | 151 |
| **245** | M06_B05_H10a | LINE | L1-Tx1 | 10014 | 0.048082064 | 37 |
| **246** | M06_B05_H10a | LINE | L2 | 530000 | 2.544786685 | 1959 |
| **247** | M06_B05_H10a | LINE | Penelope | 43310 | 0.207952286 | 224 |
| **248** | M06_B05_H10a | LINE | Proto2 | 44757 | 0.214900033 | 137 |
| **249** | M06_B05_H10a | LINE | R1 | 218757 | 1.050358304 | 463 |
| **250** | M06_B05_H10a | LINE | R1-LOA | 51864 | 0.249024182 | 73 |
| **251** | M06_B05_H10a | LINE | R2 | 345 | 0.001656512 | 3 |
| **252** | M06_B05_H10a | LINE | RTE | 28376 | 0.136246919 | 215 |
| **253** | M06_B05_H10a | LINE | RTE-BovB | 197285 | 0.947260832 | 569 |
| **254** | M06_B05_H10a | LINE | RTE-RTE | 432309 | 2.075724881 | 2095 |
| **255** | M06_B05_H10a | LINE | RTE-X | 31504 | 0.151265962 | 39 |
| **256** | M06_B05_H10a | LINE | Unknown | 2604376 | 12.50487051 | 14077 |
| **257** | M06_B05_H10a | LTR | Copia | 235 | 0.001128349 | 1 |
| **258** | M06_B05_H10a | LTR | Gypsy | 179263 | 0.860728482 | 264 |
| **259** | M06_B05_H10a | LTR | Pao | 125780 | 0.603930697 | 123 |
| **260** | M06_B05_H10a | RC | Helitron | 1903662 | 9.140403228 | 8996 |
| **261** | M06_B05_H10a | SINE | SINE | 474489 | 2.27825149 | 2740 |
| **262** | M07_B18_H16 | DNA | Academ-1 | 1172 | 0.007019127 | 5 |
| **263** | M07_B18_H16 | DNA | CMC-Transib | 14291 | 0.085589031 | 39 |
| **264** | M07_B18_H16 | DNA | Ginger-2 | 3305 | 0.019793699 | 2 |
| **265** | M07_B18_H16 | DNA | Maveric_Unknown | 262 | 0.001569122 | 2 |
| **266** | M07_B18_H16 | DNA | P | 651 | 0.00389885 | 3 |
| **267** | M07_B18_H16 | DNA | PIF-Harbing | 7646 | 0.045792018 | 51 |
| **268** | M07_B18_H16 | DNA | PIF-Spy | 12819 | 0.076773199 | 71 |
| **269** | M07_B18_H16 | DNA | PiggyBac | 34531 | 0.206806721 | 67 |
| **270** | M07_B18_H16 | DNA | Sola-1 | 35354 | 0.211735681 | 118 |
| **271** | M07_B18_H16 | DNA | Sola-2 | 5113 | 0.03062184 | 34 |
| **272** | M07_B18_H16 | DNA | TcMar-Fot1 | 23589 | 0.141274905 | 149 |
| **273** | M07_B18_H16 | DNA | TcMar-Marin | 113176 | 0.677812905 | 640 |
| **274** | M07_B18_H16 | DNA | TcMar-Tc1 | 84660 | 0.507030117 | 316 |
| **275** | M07_B18_H16 | DNA | TcMar-m44 | 11338 | 0.067903466 | 55 |
| **276** | M07_B18_H16 | DNA | Zator | 414 | 0.002479453 | 5 |
| **277** | M07_B18_H16 | DNA | hAT-Ac | 25611 | 0.153384696 | 28 |
| **278** | M07_B18_H16 | DNA | hAT-Tip100 | 869 | 0.005204455 | 7 |
| **279** | M07_B18_H16 | DNA | hAT-hATm | 93 | 0.000556979 | 3 |
| **280** | M07_B18_H16 | DNA | hAT-hATx | 2955 | 0.017697543 | 3 |

| | | | | | | |
|---|---|---|---|---|---|---|
| 281 | M07_B18_H16 | LINE | CR1 | 205912 | 1.233210317 | 418 |
| 282 | M07_B18_H16 | LINE | CR1-Zenon | 117325 | 0.702661333 | 496 |
| 283 | M07_B18_H16 | LINE | CRE | 62655 | 0.375241814 | 311 |
| 284 | M07_B18_H16 | LINE | Dong-R4 | 71585 | 0.428723729 | 181 |
| 285 | M07_B18_H16 | LINE | I | 83970 | 0.502897696 | 338 |
| 286 | M07_B18_H16 | LINE | I-Jockey | 37002 | 0.22160558 | 60 |
| 287 | M07_B18_H16 | LINE | L1 | 12703 | 0.076078474 | 125 |
| 288 | M07_B18_H16 | LINE | L1-Tx1 | 7796 | 0.046690371 | 28 |
| 289 | M07_B18_H16 | LINE | L2 | 430686 | 2.579385459 | 1702 |
| 290 | M07_B18_H16 | LINE | Penelope | 31157 | 0.18659978 | 202 |
| 291 | M07_B18_H16 | LINE | Proto2 | 42256 | 0.253071871 | 134 |
| 292 | M07_B18_H16 | LINE | R1 | 184353 | 1.104093115 | 372 |
| 293 | M07_B18_H16 | LINE | R1-LOA | 42523 | 0.254670939 | 55 |
| 294 | M07_B18_H16 | LINE | R2 | 450 | 0.002695057 | 3 |
| 295 | M07_B18_H16 | LINE | RTE | 21228 | 0.127134837 | 179 |
| 296 | M07_B18_H16 | LINE | RTE-BovB | 140329 | 0.840432663 | 487 |
| 297 | M07_B18_H16 | LINE | RTE-RTE | 339888 | 2.03559476 | 1775 |
| 298 | M07_B18_H16 | LINE | RTE-X | 17800 | 0.106604489 | 31 |
| 299 | M07_B18_H16 | LINE | Unknown | 2193562 | 13.13727849 | 11999 |
| 300 | M07_B18_H16 | LTR | Copia | 558 | 0.003341871 | 2 |
| 301 | M07_B18_H16 | LTR | DIRS | 9446 | 0.056572248 | 3 |
| 302 | M07_B18_H16 | LTR | Gypsy | 156722 | 0.938610607 | 208 |
| 303 | M07_B18_H16 | LTR | Pao | 113050 | 0.677058289 | 97 |
| 304 | M07_B18_H16 | RC | Helitron | 1567719 | 9.389094588 | 7494 |
| 305 | M07_B18_H16 | SINE | SINE | 414336 | 2.481465043 | 2353 |
| 306 | M08_B17_H15 | DNA | Academ-1 | 4181 | 0.023440306 | 14 |
| 307 | M08_B17_H15 | DNA | CMC-Chapaev | 1155 | 0.006475377 | 6 |
| 308 | M08_B17_H15 | DNA | CMC-Transib | 17026 | 0.095454352 | 51 |
| 309 | M08_B17_H15 | DNA | Ginger-2 | 3063 | 0.017172365 | 3 |
| 310 | M08_B17_H15 | DNA | Maveric_Unknown | 905 | 0.005073781 | 4 |
| 311 | M08_B17_H15 | DNA | P | 178 | 0.000997937 | 2 |
| 312 | M08_B17_H15 | DNA | PIF-Harbing | 12672 | 0.071044141 | 49 |
| 313 | M08_B17_H15 | DNA | PIF-Spy | 14378 | 0.080608638 | 69 |
| 314 | M08_B17_H15 | DNA | PiggyBac | 46361 | 0.259917727 | 75 |
| 315 | M08_B17_H15 | DNA | Sola-1 | 50878 | 0.285241779 | 135 |
| 316 | M08_B17_H15 | DNA | Sola-2 | 6067 | 0.034013953 | 25 |
| 317 | M08_B17_H15 | DNA | TcMar-Fot1 | 22600 | 0.126704356 | 131 |
| 318 | M08_B17_H15 | DNA | TcMar-Marin | 147059 | 0.824469728 | 802 |
| 319 | M08_B17_H15 | DNA | TcMar-Tc1 | 98715 | 0.553434535 | 365 |
| 320 | M08_B17_H15 | DNA | TcMar-m44 | 18836 | 0.105601914 | 67 |
| 321 | M08_B17_H15 | DNA | Zator | 359 | 0.002012693 | 1 |
| 322 | M08_B17_H15 | DNA | hAT-Ac | 23762 | 0.133218978 | 39 |
| 323 | M08_B17_H15 | DNA | hAT-Tip100 | 1589 | 0.00890855 | 5 |
| 324 | M08_B17_H15 | DNA | hAT-hATx | 305 | 0.001709948 | 3 |
| 325 | M08_B17_H15 | LINE | CR1 | 176785 | 0.991125201 | 447 |
| 326 | M08_B17_H15 | LINE | CR1-Zenon | 127837 | 0.716703749 | 514 |
| 327 | M08_B17_H15 | LINE | CRE | 62602 | 0.350971066 | 328 |

| | | | | | | |
|---|---|---|---|---|---|---|
| 328 | M08_B17_H15 | LINE | Dong-R4 | 53116 | 0.297788874 | 159 |
| 329 | M08_B17_H15 | LINE | I | 98239 | 0.550765894 | 356 |
| 330 | M08_B17_H15 | LINE | I-Jockey | 26920 | 0.150923949 | 72 |
| 331 | M08_B17_H15 | LINE | L1 | 11441 | 0.064142679 | 127 |
| 332 | M08_B17_H15 | LINE | L1-Tx1 | 7754 | 0.043471928 | 41 |
| 333 | M08_B17_H15 | LINE | L2 | 466396 | 2.614796669 | 1764 |
| 334 | M08_B17_H15 | LINE | Penelope | 34904 | 0.195685347 | 196 |
| 335 | M08_B17_H15 | LINE | Proto2 | 52645 | 0.295148266 | 139 |
| 336 | M08_B17_H15 | LINE | R1 | 169777 | 0.951835638 | 374 |
| 337 | M08_B17_H15 | LINE | R1-LOA | 49752 | 0.278928987 | 60 |
| 338 | M08_B17_H15 | LINE | R2 | 144 | 0.00080732 | 2 |
| 339 | M08_B17_H15 | LINE | RTE | 28162 | 0.157887083 | 209 |
| 340 | M08_B17_H15 | LINE | RTE-BovB | 183604 | 1.029355157 | 526 |
| 341 | M08_B17_H15 | LINE | RTE-RTE | 404548 | 2.26805282 | 1965 |
| 342 | M08_B17_H15 | LINE | RTE-X | 22493 | 0.126104472 | 35 |
| 343 | M08_B17_H15 | LINE | Unknown | 2303760 | 12.91577109 | 12746 |
| 344 | M08_B17_H15 | LTR | DIRS | 24 | 0.000134553 | 1 |
| 345 | M08_B17_H15 | LTR | Gypsy | 186840 | 1.047497426 | 233 |
| 346 | M08_B17_H15 | LTR | Pao | 155312 | 0.870739244 | 134 |
| 347 | M08_B17_H15 | RC | Helitron | 1716861 | 9.62538792 | 7969 |
| 348 | M08_B17_H15 | SINE | SINE | 424848 | 2.381862485 | 2473 |
| 349 | M09_B10_H20a | DNA | Academ-1 | 1644 | 0.008877463 | 8 |
| 350 | M09_B10_H20a | DNA | CMC-Chapaev | 2061 | 0.011129228 | 1 |
| 351 | M09_B10_H20a | DNA | CMC-Transib | 25125 | 0.135672901 | 66 |
| 352 | M09_B10_H20a | DNA | Ginger-2 | 1923 | 0.010384039 | 2 |
| 353 | M09_B10_H20a | DNA | Maveric_Unknown | 35 | 0.000188997 | 1 |
| 354 | M09_B10_H20a | DNA | P | 201 | 0.001085383 | 1 |
| 355 | M09_B10_H20a | DNA | PIF-Harbing | 8462 | 0.045694093 | 40 |
| 356 | M09_B10_H20a | DNA | PIF-Spy | 7828 | 0.042270546 | 47 |
| 357 | M09_B10_H20a | DNA | PiggyBac | 33470 | 0.180735204 | 60 |
| 358 | M09_B10_H20a | DNA | Sola-1 | 22455 | 0.121255124 | 97 |
| 359 | M09_B10_H20a | DNA | Sola-2 | 18488 | 0.099833656 | 26 |
| 360 | M09_B10_H20a | DNA | TcMar-Fot1 | 30687 | 0.165707237 | 127 |
| 361 | M09_B10_H20a | DNA | TcMar-Marin | 123034 | 0.664373322 | 645 |
| 362 | M09_B10_H20a | DNA | TcMar-Tc1 | 102607 | 0.554069229 | 373 |
| 363 | M09_B10_H20a | DNA | TcMar-m44 | 17422 | 0.094077345 | 78 |
| 364 | M09_B10_H20a | DNA | Zator | 592 | 0.003196751 | 5 |
| 365 | M09_B10_H20a | DNA | hAT-Ac | 17807 | 0.096156312 | 37 |
| 366 | M09_B10_H20a | DNA | hAT-Tip100 | 1112 | 0.006004707 | 8 |
| 367 | M09_B10_H20a | LINE | CR1 | 170030 | 0.918147796 | 380 |
| 368 | M09_B10_H20a | LINE | CR1-Zenon | 153269 | 0.827639796 | 568 |
| 369 | M09_B10_H20a | LINE | CRE | 71645 | 0.386877015 | 341 |
| 370 | M09_B10_H20a | LINE | Dong-R4 | 34133 | 0.184315349 | 132 |
| 371 | M09_B10_H20a | LINE | I | 102109 | 0.55138007 | 343 |
| 372 | M09_B10_H20a | LINE | I-Jockey | 33091 | 0.178688636 | 67 |
| 373 | M09_B10_H20a | LINE | L1 | 10900 | 0.058859089 | 124 |
| 374 | M09_B10_H20a | LINE | L1-Tx1 | 6203 | 0.033495682 | 26 |

| | | | | | | |
|---|---|---|---|---|---|---|
| 375 | M09_B10_H20a | LINE | L2 | 436872 | 2.359072305 | 1729 |
| 376 | M09_B10_H20a | LINE | PLE_Unknown | 51 | 0.000275396 | 3 |
| 377 | M09_B10_H20a | LINE | Penelope | 26792 | 0.144674562 | 192 |
| 378 | M09_B10_H20a | LINE | Proto2 | 35634 | 0.192420623 | 104 |
| 379 | M09_B10_H20a | LINE | R1 | 176750 | 0.954435235 | 342 |
| 380 | M09_B10_H20a | LINE | R1-LOA | 45445 | 0.245399204 | 60 |
| 381 | M09_B10_H20a | LINE | RTE | 30935 | 0.167046416 | 195 |
| 382 | M09_B10_H20a | LINE | RTE-BovB | 157834 | 0.852290415 | 456 |
| 383 | M09_B10_H20a | LINE | RTE-RTE | 393749 | 2.126211708 | 1827 |
| 384 | M09_B10_H20a | LINE | RTE-X | 20216 | 0.109164711 | 29 |
| 385 | M09_B10_H20a | LINE | Tad1 | 156 | 0.000842387 | 2 |
| 386 | M09_B10_H20a | LINE | Unknown | 2239025 | 12.09054796 | 12100 |
| 387 | M09_B10_H20a | LTR | Copia | 819 | 0.004422532 | 3 |
| 388 | M09_B10_H20a | LTR | Gypsy | 132503 | 0.715505131 | 203 |
| 389 | M09_B10_H20a | LTR | Pao | 128162 | 0.692064094 | 103 |
| 390 | M09_B10_H20a | RC | Helitron | 1626361 | 8.782213539 | 7643 |
| 391 | M09_B10_H20a | SINE | SINE | 413898 | 2.235014624 | 2376 |
| 392 | M10_B09_H06a | DNA | Academ-1 | 2963 | 0.015990455 | 13 |
| 393 | M10_B09_H06a | DNA | CMC-Chapaev | 1010 | 0.005450678 | 5 |
| 394 | M10_B09_H06a | DNA | CMC-Transib | 14977 | 0.080826543 | 57 |
| 395 | M10_B09_H06a | DNA | Ginger-2 | 2029 | 0.010949927 | 5 |
| 396 | M10_B09_H06a | DNA | Kolobok-Hyd | 116 | 0.000626018 | 1 |
| 397 | M10_B09_H06a | DNA | Maveric_Unknown | 7089 | 0.038257285 | 1 |
| 398 | M10_B09_H06a | DNA | P | 1165 | 0.006287168 | 7 |
| 399 | M10_B09_H06a | DNA | PIF-Harbing | 9749 | 0.052612537 | 52 |
| 400 | M10_B09_H06a | DNA | PIF-Spy | 15701 | 0.084733762 | 67 |
| 401 | M10_B09_H06a | DNA | PiggyBac | 19505 | 0.105262851 | 51 |
| 402 | M10_B09_H06a | DNA | Sola-1 | 38370 | 0.207071807 | 133 |
| 403 | M10_B09_H06a | DNA | Sola-2 | 12886 | 0.06954202 | 29 |
| 404 | M10_B09_H06a | DNA | TcMar-Fot1 | 39767 | 0.214611013 | 166 |
| 405 | M10_B09_H06a | DNA | TcMar-Marin | 143913 | 0.776656893 | 816 |
| 406 | M10_B09_H06a | DNA | TcMar-Tc1 | 102804 | 0.554803494 | 367 |
| 407 | M10_B09_H06a | DNA | TcMar-m44 | 12053 | 0.06504656 | 65 |
| 408 | M10_B09_H06a | DNA | Zator | 1019 | 0.005499249 | 5 |
| 409 | M10_B09_H06a | DNA | hAT-Ac | 44954 | 0.242603753 | 55 |
| 410 | M10_B09_H06a | DNA | hAT-Tip100 | 2256 | 0.01217498 | 10 |
| 411 | M10_B09_H06a | DNA | hAT-hATm | 20 | 0.000107934 | 1 |
| 412 | M10_B09_H06a | DNA | hAT-hATx | 113 | 0.000609828 | 1 |
| 413 | M10_B09_H06a | LINE | CR1 | 206152 | 1.112542799 | 475 |
| 414 | M10_B09_H06a | LINE | CR1-Zenon | 135366 | 0.730531203 | 546 |
| 415 | M10_B09_H06a | LINE | CRE | 53111 | 0.286624726 | 316 |
| 416 | M10_B09_H06a | LINE | Dong-R4 | 69397 | 0.374515564 | 185 |
| 417 | M10_B09_H06a | LINE | I | 125459 | 0.677065985 | 389 |
| 418 | M10_B09_H06a | LINE | I-Jockey | 29504 | 0.159224566 | 60 |
| 419 | M10_B09_H06a | LINE | L1 | 14301 | 0.077178366 | 140 |
| 420 | M10_B09_H06a | LINE | L1-Tx1 | 5807 | 0.031338702 | 31 |
| 421 | M10_B09_H06a | LINE | L2 | 498115 | 2.688182778 | 1965 |

| | | | | | | |
|---|---|---|---|---|---|---|
| 422 | M10_B09_H06a | LINE | PLE_Unknown | 114 | 0.000615225 | 1 |
| 423 | M10_B09_H06a | LINE | Penelope | 31522 | 0.170115129 | 199 |
| 424 | M10_B09_H06a | LINE | Proto2 | 29163 | 0.157384287 | 118 |
| 425 | M10_B09_H06a | LINE | R1 | 179156 | 0.966853184 | 375 |
| 426 | M10_B09_H06a | LINE | R1-LOA | 49669 | 0.268049247 | 43 |
| 427 | M10_B09_H06a | LINE | R2 | 48 | 0.000259042 | 1 |
| 428 | M10_B09_H06a | LINE | RTE | 24041 | 0.129742333 | 198 |
| 429 | M10_B09_H06a | LINE | RTE-BovB | 154490 | 0.833737907 | 524 |
| 430 | M10_B09_H06a | LINE | RTE-RTE | 496330 | 2.678549649 | 2082 |
| 431 | M10_B09_H06a | LINE | RTE-X | 25943 | 0.140006878 | 34 |
| 432 | M10_B09_H06a | LINE | Tad1 | 47 | 0.000253645 | 1 |
| 433 | M10_B09_H06a | LINE | Unknown | 2322195 | 12.53221567 | 13129 |
| 434 | M10_B09_H06a | LTR | Copia | 4418 | 0.02384267 | 2 |
| 435 | M10_B09_H06a | LTR | DIRS | 361 | 0.001948213 | 1 |
| 436 | M10_B09_H06a | LTR | Gypsy | 180052 | 0.971688637 | 246 |
| 437 | M10_B09_H06a | LTR | Pao | 124296 | 0.67078961 | 102 |
| 438 | M10_B09_H06a | RC | Helitron | 1738010 | 9.379537959 | 8232 |
| 439 | M10_B09_H06a | SINE | SINE | 445652 | 2.405055121 | 2626 |
| 440 | M11_B22_H13a | DNA | Academ-1 | 2157 | 0.011637644 | 5 |
| 441 | M11_B22_H13a | DNA | CMC-Chapaev | 844 | 0.004553626 | 2 |
| 442 | M11_B22_H13a | DNA | CMC-Transib | 8483 | 0.045768257 | 46 |
| 443 | M11_B22_H13a | DNA | Ginger-2 | 3288 | 0.017739718 | 2 |
| 444 | M11_B22_H13a | DNA | Maveric_Unknown | 392 | 0.002114954 | 2 |
| 445 | M11_B22_H13a | DNA | P | 203 | 0.001095244 | 1 |
| 446 | M11_B22_H13a | DNA | PIF-Harbing | 9210 | 0.049690634 | 53 |
| 447 | M11_B22_H13a | DNA | PIF-Spy | 21656 | 0.116840431 | 79 |
| 448 | M11_B22_H13a | DNA | PiggyBac | 24219 | 0.130668563 | 57 |
| 449 | M11_B22_H13a | DNA | Sola-1 | 30357 | 0.163784862 | 124 |
| 450 | M11_B22_H13a | DNA | Sola-2 | 13600 | 0.073375963 | 26 |
| 451 | M11_B22_H13a | DNA | TcMar-Fot1 | 16245 | 0.087646509 | 121 |
| 452 | M11_B22_H13a | DNA | TcMar-Marin | 143446 | 0.773932973 | 782 |
| 453 | M11_B22_H13a | DNA | TcMar-Tc1 | 111157 | 0.599724409 | 365 |
| 454 | M11_B22_H13a | DNA | TcMar-m44 | 10658 | 0.057503016 | 47 |
| 455 | M11_B22_H13a | DNA | Zator | 74 | 0.000399252 | 1 |
| 456 | M11_B22_H13a | DNA | hAT-Ac | 37834 | 0.204125456 | 42 |
| 457 | M11_B22_H13a | DNA | hAT-Tip100 | 836 | 0.004510464 | 6 |
| 458 | M11_B22_H13a | DNA | hAT-hATm | 1152 | 0.006215376 | 3 |
| 459 | M11_B22_H13a | LINE | CR1 | 226780 | 1.223544189 | 476 |
| 460 | M11_B22_H13a | LINE | CR1-Zenon | 147223 | 0.794310989 | 531 |
| 461 | M11_B22_H13a | LINE | CRE | 79032 | 0.426400672 | 368 |
| 462 | M11_B22_H13a | LINE | Dong-R4 | 47129 | 0.254274689 | 178 |
| 463 | M11_B22_H13a | LINE | I | 93682 | 0.505441691 | 323 |
| 464 | M11_B22_H13a | LINE | I-Jockey | 17838 | 0.096241208 | 52 |
| 465 | M11_B22_H13a | LINE | L1 | 14576 | 0.078641768 | 160 |
| 466 | M11_B22_H13a | LINE | L1-Tx1 | 9045 | 0.048800411 | 42 |
| 467 | M11_B22_H13a | LINE | L2 | 542517 | 2.927037316 | 1892 |
| 468 | M11_B22_H13a | LINE | Penelope | 35988 | 0.194165748 | 226 |

| | | | | | | |
|---|---|---|---|---|---|---|
| **469** | M11_B22_H13a | LINE | Proto2 | 53846 | 0.290514862 | 175 |
| **470** | M11_B22_H13a | LINE | R1 | 186543 | 1.006453848 | 425 |
| **471** | M11_B22_H13a | LINE | R1-LOA | 41255 | 0.222582748 | 46 |
| **472** | M11_B22_H13a | LINE | RTE | 23289 | 0.125650942 | 164 |
| **473** | M11_B22_H13a | LINE | RTE-BovB | 168854 | 0.911016538 | 487 |
| **474** | M11_B22_H13a | LINE | RTE-RTE | 382408 | 2.063202602 | 1953 |
| **475** | M11_B22_H13a | LINE | RTE-X | 20815 | 0.112302991 | 29 |
| **476** | M11_B22_H13a | LINE | Unknown | 2386262 | 12.8745789 | 13116 |
| **477** | M11_B22_H13a | LTR | Copia | 5362 | 0.028929553 | 5 |
| **478** | M11_B22_H13a | LTR | Gypsy | 172679 | 0.931653527 | 233 |
| **479** | M11_B22_H13a | LTR | Pao | 119062 | 0.642374187 | 87 |
| **480** | M11_B22_H13a | RC | Helitron | 1658058 | 8.945706103 | 7919 |
| **481** | M11_B22_H13a | SINE | SINE | 430349 | 2.321858268 | 2503 |
| **482** | M12_B11a_H07a | DNA | Academ-1 | 1050 | 0.006048594 | 5 |
| **483** | M12_B11a_H07a | DNA | CMC-Chapaev | 307 | 0.001768494 | 1 |
| **484** | M12_B11a_H07a | DNA | CMC-Transib | 14444 | 0.083205612 | 39 |
| **485** | M12_B11a_H07a | DNA | Ginger-2 | 5450 | 0.031395083 | 6 |
| **486** | M12_B11a_H07a | DNA | Maveric_Unknown | 622 | 0.003583072 | 6 |
| **487** | M12_B11a_H07a | DNA | P | 662 | 0.003813495 | 6 |
| **488** | M12_B11a_H07a | DNA | PIF-Harbing | 13642 | 0.078585638 | 52 |
| **489** | M12_B11a_H07a | DNA | PIF-Spy | 8162 | 0.047017738 | 61 |
| **490** | M12_B11a_H07a | DNA | PiggyBac | 16755 | 0.096518279 | 48 |
| **491** | M12_B11a_H07a | DNA | Sola-1 | 29669 | 0.170910226 | 126 |
| **492** | M12_B11a_H07a | DNA | Sola-2 | 3630 | 0.020910854 | 37 |
| **493** | M12_B11a_H07a | DNA | TcMar-Fot1 | 13340 | 0.076845947 | 132 |
| **494** | M12_B11a_H07a | DNA | TcMar-Marin | 129504 | 0.74601631 | 730 |
| **495** | M12_B11a_H07a | DNA | TcMar-Tc1 | 90686 | 0.522402667 | 301 |
| **496** | M12_B11a_H07a | DNA | TcMar-m44 | 21332 | 0.122884389 | 78 |
| **497** | M12_B11a_H07a | DNA | Zator | 1052 | 0.006060115 | 6 |
| **498** | M12_B11a_H07a | DNA | hAT-Ac | 19692 | 0.113437061 | 24 |
| **499** | M12_B11a_H07a | DNA | hAT-Charlie | 1965 | 0.011319512 | 1 |
| **500** | M12_B11a_H07a | DNA | hAT-Tip100 | 1269 | 0.007310158 | 6 |
| **501** | M12_B11a_H07a | DNA | hAT-hATm | 3248 | 0.018710318 | 3 |
| **502** | M12_B11a_H07a | DNA | hAT-hATx | 98 | 0.000564535 | 1 |
| **503** | M12_B11a_H07a | LINE | CR1 | 188708 | 1.087064845 | 416 |
| **504** | M12_B11a_H07a | LINE | CR1-Zenon | 111926 | 0.644757084 | 470 |
| **505** | M12_B11a_H07a | LINE | CRE | 84727 | 0.488075456 | 336 |
| **506** | M12_B11a_H07a | LINE | Dong-R4 | 47177 | 0.271766211 | 119 |
| **507** | M12_B11a_H07a | LINE | I | 152968 | 0.881182225 | 402 |
| **508** | M12_B11a_H07a | LINE | I-Jockey | 25228 | 0.145327553 | 50 |
| **509** | M12_B11a_H07a | LINE | L1 | 13288 | 0.076546398 | 150 |
| **510** | M12_B11a_H07a | LINE | L1-Tx1 | 4959 | 0.028566646 | 25 |
| **511** | M12_B11a_H07a | LINE | L2 | 437510 | 2.52030513 | 1648 |
| **512** | M12_B11a_H07a | LINE | Penelope | 29783 | 0.17156693 | 154 |
| **513** | M12_B11a_H07a | LINE | Proto2 | 35955 | 0.207121142 | 108 |
| **514** | M12_B11a_H07a | LINE | R1 | 183903 | 1.059385327 | 343 |
| **515** | M12_B11a_H07a | LINE | R1-LOA | 35041 | 0.201855985 | 58 |

| | | | | | | |
|---|---|---|---|---|---|---|
| 516 | M12_B11a_H07a | LINE | R2 | 192 | 0.001106029 | 2 |
| 517 | M12_B11a_H07a | LINE | RTE | 26925 | 0.155103233 | 189 |
| 518 | M12_B11a_H07a | LINE | RTE-BovB | 162106 | 0.933822275 | 474 |
| 519 | M12_B11a_H07a | LINE | RTE-RTE | 395372 | 2.27756641 | 1741 |
| 520 | M12_B11a_H07a | LINE | RTE-X | 24561 | 0.141485256 | 31 |
| 521 | M12_B11a_H07a | LINE | Tad1 | 70 | 0.00040324 | 1 |
| 522 | M12_B11a_H07a | LINE | Unknown | 2152298 | 12.39845419 | 11859 |
| 523 | M12_B11a_H07a | LTR | Copia | 891 | 0.005132664 | 3 |
| 524 | M12_B11a_H07a | LTR | DIRS | 839 | 0.004833115 | 1 |
| 525 | M12_B11a_H07a | LTR | Gypsy | 232681 | 1.340374204 | 266 |
| 526 | M12_B11a_H07a | LTR | Pao | 117127 | 0.674717787 | 101 |
| 527 | M12_B11a_H07a | RC | Helitron | 1536576 | 8.851547109 | 7365 |
| 528 | M12_B11a_H07a | SINE | SINE | 418202 | 2.409080126 | 2395 |
| 529 | M13_B08_H12a | DNA | Academ-1 | 2078 | 0.011678918 | 10 |
| 530 | M13_B08_H12a | DNA | CMC-Chapaev | 849 | 0.004771608 | 2 |
| 531 | M13_B08_H12a | DNA | CMC-Transib | 17556 | 0.098669435 | 48 |
| 532 | M13_B08_H12a | DNA | Ginger-2 | 353 | 0.001983955 | 1 |
| 533 | M13_B08_H12a | DNA | Maveric_Unknown | 985 | 0.005535965 | 5 |
| 534 | M13_B08_H12a | DNA | PIF-Harbing | 14816 | 0.083269899 | 40 |
| 535 | M13_B08_H12a | DNA | PIF-Spy | 10512 | 0.059080263 | 58 |
| 536 | M13_B08_H12a | DNA | PiggyBac | 45527 | 0.255873968 | 75 |
| 537 | M13_B08_H12a | DNA | Sola-1 | 41392 | 0.232634157 | 145 |
| 538 | M13_B08_H12a | DNA | Sola-2 | 7347 | 0.041292113 | 23 |
| 539 | M13_B08_H12a | DNA | TcMar-Fot1 | 13054 | 0.073366986 | 107 |
| 540 | M13_B08_H12a | DNA | TcMar-Marin | 151395 | 0.850880561 | 735 |
| 541 | M13_B08_H12a | DNA | TcMar-Tc1 | 110648 | 0.621871477 | 362 |
| 542 | M13_B08_H12a | DNA | TcMar-m44 | 19491 | 0.109544655 | 85 |
| 543 | M13_B08_H12a | DNA | Zator | 3303 | 0.018563747 | 7 |
| 544 | M13_B08_H12a | DNA | hAT-Ac | 31724 | 0.1782974 | 37 |
| 545 | M13_B08_H12a | DNA | hAT-Tip100 | 942 | 0.005294293 | 9 |
| 546 | M13_B08_H12a | DNA | hAT-hATm | 25 | 0.000140507 | 1 |
| 547 | M13_B08_H12a | LINE | CR1 | 193189 | 1.085774066 | 463 |
| 548 | M13_B08_H12a | LINE | CR1-Zenon | 142101 | 0.798645785 | 471 |
| 549 | M13_B08_H12a | LINE | CRE | 60722 | 0.341273949 | 333 |
| 550 | M13_B08_H12a | LINE | Dong-R4 | 38026 | 0.213716333 | 144 |
| 551 | M13_B08_H12a | LINE | I | 132051 | 0.742162086 | 373 |
| 552 | M13_B08_H12a | LINE | I-Jockey | 28514 | 0.160256338 | 73 |
| 553 | M13_B08_H12a | LINE | L1 | 11417 | 0.064166606 | 132 |
| 554 | M13_B08_H12a | LINE | L1-Tx1 | 4230 | 0.023773736 | 16 |
| 555 | M13_B08_H12a | LINE | L2 | 466745 | 2.623232257 | 1801 |
| 556 | M13_B08_H12a | LINE | PLE_Unknown | 34 | 0.000191089 | 1 |
| 557 | M13_B08_H12a | LINE | Penelope | 34160 | 0.191988375 | 178 |
| 558 | M13_B08_H12a | LINE | Proto2 | 49801 | 0.279894995 | 118 |
| 559 | M13_B08_H12a | LINE | R1 | 138893 | 0.780615963 | 342 |
| 560 | M13_B08_H12a | LINE | R1-LOA | 46662 | 0.262252972 | 58 |
| 561 | M13_B08_H12a | LINE | R2 | 230 | 0.001292662 | 2 |
| 562 | M13_B08_H12a | LINE | RTE | 24324 | 0.136707413 | 190 |

| | | | | | | |
|---|---|---|---|---|---|---|
| 563 | M13_B08_H12a | LINE | RTE-BovB | 140415 | 0.789170012 | 460 |
| 564 | M13_B08_H12a | LINE | RTE-RTE | 373679 | 2.100176342 | 1886 |
| 565 | M13_B08_H12a | LINE | RTE-X | 19986 | 0.112326688 | 25 |
| 566 | M13_B08_H12a | LINE | Tad1 | 257 | 0.001444409 | 2 |
| 567 | M13_B08_H12a | LINE | Unknown | 2240255 | 12.59083478 | 12131 |
| 568 | M13_B08_H12a | LTR | Gypsy | 214346 | 1.204682088 | 232 |
| 569 | M13_B08_H12a | LTR | Pao | 138886 | 0.780576622 | 120 |
| 570 | M13_B08_H12a | RC | Helitron | 1729531 | 9.720428732 | 7911 |
| 571 | M13_B08_H12a | SINE | SINE | 424011 | 2.3830557 | 2472 |
| 572 | M14_B23a_H18a | DNA | Academ-1 | 1736 | 0.010016298 | 2 |
| 573 | M14_B23a_H18a | DNA | CMC-Chapaev | 738 | 0.004258081 | 1 |
| 574 | M14_B23a_H18a | DNA | CMC-Transib | 11900 | 0.068660106 | 36 |
| 575 | M14_B23a_H18a | DNA | Ginger-2 | 3653 | 0.021076922 | 4 |
| 576 | M14_B23a_H18a | DNA | Maveric_Unknown | 58 | 0.000334646 | 1 |
| 577 | M14_B23a_H18a | DNA | P | 343 | 0.001979027 | 2 |
| 578 | M14_B23a_H18a | DNA | PIF-Harbing | 15521 | 0.089552396 | 48 |
| 579 | M14_B23a_H18a | DNA | PIF-Spy | 10324 | 0.059566969 | 64 |
| 580 | M14_B23a_H18a | DNA | PiggyBac | 29641 | 0.171021362 | 48 |
| 581 | M14_B23a_H18a | DNA | Sola-1 | 40489 | 0.233611684 | 124 |
| 582 | M14_B23a_H18a | DNA | Sola-2 | 16487 | 0.095125981 | 37 |
| 583 | M14_B23a_H18a | DNA | TcMar-Fot1 | 24879 | 0.14354578 | 135 |
| 584 | M14_B23a_H18a | DNA | TcMar-Marin | 146702 | 0.846434864 | 686 |
| 585 | M14_B23a_H18a | DNA | TcMar-Tc1 | 89715 | 0.517633733 | 300 |
| 586 | M14_B23a_H18a | DNA | TcMar-m44 | 14333 | 0.082697924 | 48 |
| 587 | M14_B23a_H18a | DNA | Zator | 208 | 0.001200109 | 2 |
| 588 | M14_B23a_H18a | DNA | hAT-Ac | 28638 | 0.165234296 | 44 |
| 589 | M14_B23a_H18a | DNA | hAT-Pegasus | 5270 | 0.030406618 | 3 |
| 590 | M14_B23a_H18a | DNA | hAT-Tip100 | 1288 | 0.007431447 | 6 |
| 591 | M14_B23a_H18a | DNA | hAT-hATm | 624 | 0.003600328 | 1 |
| 592 | M14_B23a_H18a | LINE | CR1 | 191387 | 1.104256448 | 411 |
| 593 | M14_B23a_H18a | LINE | CR1-Zenon | 103246 | 0.595704312 | 475 |
| 594 | M14_B23a_H18a | LINE | CRE | 69829 | 0.402896349 | 349 |
| 595 | M14_B23a_H18a | LINE | Dong-R4 | 54727 | 0.315761481 | 152 |
| 596 | M14_B23a_H18a | LINE | I | 89740 | 0.517777977 | 340 |
| 597 | M14_B23a_H18a | LINE | I-Jockey | 31051 | 0.179156719 | 39 |
| 598 | M14_B23a_H18a | LINE | L1 | 11411 | 0.065838695 | 125 |
| 599 | M14_B23a_H18a | LINE | L1-Tx1 | 4086 | 0.023575226 | 17 |
| 600 | M14_B23a_H18a | LINE | L2 | 439888 | 2.538046786 | 1688 |
| 601 | M14_B23a_H18a | LINE | Penelope | 26000 | 0.150013677 | 195 |
| 602 | M14_B23a_H18a | LINE | Proto2 | 49218 | 0.283975891 | 112 |
| 603 | M14_B23a_H18a | LINE | R1 | 174354 | 1.00598018 | 342 |
| 604 | M14_B23a_H18a | LINE | R1-LOA | 49809 | 0.287385817 | 42 |
| 605 | M14_B23a_H18a | LINE | R2 | 295 | 0.001702078 | 3 |
| 606 | M14_B23a_H18a | LINE | RTE | 23492 | 0.135543127 | 172 |
| 607 | M14_B23a_H18a | LINE | RTE-BovB | 163902 | 0.945674682 | 485 |
| 608 | M14_B23a_H18a | LINE | RTE-RTE | 388352 | 2.240696599 | 1851 |
| 609 | M14_B23a_H18a | LINE | RTE-X | 25688 | 0.148213513 | 40 |

| | | | | | | |
|---|---|---|---|---|---|---|
| 610 | M14_B23a_H18a | LINE | Tad1 | 23 | 0.000132704 | 1 |
| 611 | M14_B23a_H18a | LINE | Unknown | 2133928 | 12.31224562 | 12032 |
| 612 | M14_B23a_H18a | LTR | Copia | 328 | 0.00189248 | 1 |
| 613 | M14_B23a_H18a | LTR | DIRS | 5145 | 0.029685399 | 4 |
| 614 | M14_B23a_H18a | LTR | Gypsy | 184322 | 1.063493116 | 225 |
| 615 | M14_B23a_H18a | LTR | Pao | 151069 | 0.871631392 | 152 |
| 616 | M14_B23a_H18a | RC | Helitron | 1542771 | 8.901413492 | 7427 |
| 617 | M14_B23a_H18a | SINE | SINE | 448865 | 2.589841893 | 2570 |
| 618 | M15_B13_H17a | DNA | Academ-1 | 784 | 0.004034075 | 4 |
| 619 | M15_B13_H17a | DNA | CMC-Chapaev | 1005 | 0.005171231 | 2 |
| 620 | M15_B13_H17a | DNA | CMC-Transib | 13359 | 0.068738778 | 46 |
| 621 | M15_B13_H17a | DNA | Ginger-2 | 3274 | 0.016846378 | 4 |
| 622 | M15_B13_H17a | DNA | Kolobok-Hyd | 650 | 0.003344577 | 2 |
| 623 | M15_B13_H17a | DNA | Maveric_Unknown | 8449 | 0.043474357 | 10 |
| 624 | M15_B13_H17a | DNA | P | 594 | 0.003056429 | 2 |
| 625 | M15_B13_H17a | DNA | PIF-Harbing | 5855 | 0.030126922 | 52 |
| 626 | M15_B13_H17a | DNA | PIF-Spy | 5740 | 0.029535189 | 58 |
| 627 | M15_B13_H17a | DNA | PiggyBac | 36051 | 0.185500538 | 64 |
| 628 | M15_B13_H17a | DNA | Sola-1 | 59602 | 0.306682285 | 157 |
| 629 | M15_B13_H17a | DNA | Sola-2 | 7677 | 0.039502028 | 17 |
| 630 | M15_B13_H17a | DNA | TcMar-Fot1 | 18206 | 0.093679032 | 135 |
| 631 | M15_B13_H17a | DNA | TcMar-Marin | 140476 | 0.722819715 | 801 |
| 632 | M15_B13_H17a | DNA | TcMar-Tc1 | 119143 | 0.613050694 | 374 |
| 633 | M15_B13_H17a | DNA | TcMar-m44 | 9946 | 0.051177175 | 53 |
| 634 | M15_B13_H17a | DNA | Zator | 2029 | 0.010440226 | 3 |
| 635 | M15_B13_H17a | DNA | hAT-Ac | 21660 | 0.1114516 | 24 |
| 636 | M15_B13_H17a | DNA | hAT-Tip100 | 1175 | 0.006045966 | 10 |
| 637 | M15_B13_H17a | DNA | hAT-hATx | 344 | 0.001770053 | 3 |
| 638 | M15_B13_H17a | LINE | CR1 | 200469 | 1.031513892 | 461 |
| 639 | M15_B13_H17a | LINE | CR1-Zenon | 116214 | 0.597979515 | 525 |
| 640 | M15_B13_H17a | LINE | CRE | 94858 | 0.488092148 | 392 |
| 641 | M15_B13_H17a | LINE | Dong-R4 | 69095 | 0.355528547 | 191 |
| 642 | M15_B13_H17a | LINE | I | 116237 | 0.598097862 | 363 |
| 643 | M15_B13_H17a | LINE | I-Jockey | 29518 | 0.151884965 | 59 |
| 644 | M15_B13_H17a | LINE | L1 | 14573 | 0.074985419 | 150 |
| 645 | M15_B13_H17a | LINE | L1-Tx1 | 6956 | 0.035792121 | 33 |
| 646 | M15_B13_H17a | LINE | L2 | 496341 | 2.553924231 | 1936 |
| 647 | M15_B13_H17a | LINE | PLE_Unknown | 28 | 0.000144074 | 1 |
| 648 | M15_B13_H17a | LINE | Penelope | 32328 | 0.166343829 | 182 |
| 649 | M15_B13_H17a | LINE | Proto2 | 45591 | 0.234588639 | 135 |
| 650 | M15_B13_H17a | LINE | R1 | 192403 | 0.990010263 | 407 |
| 651 | M15_B13_H17a | LINE | R1-LOA | 36769 | 0.189195009 | 60 |
| 652 | M15_B13_H17a | LINE | R2 | 412 | 0.002119947 | 3 |
| 653 | M15_B13_H17a | LINE | RTE | 23488 | 0.12085758 | 195 |
| 654 | M15_B13_H17a | LINE | RTE-BovB | 180221 | 0.927327742 | 486 |
| 655 | M15_B13_H17a | LINE | RTE-RTE | 409905 | 2.109167512 | 1982 |
| 656 | M15_B13_H17a | LINE | RTE-X | 39047 | 0.200916466 | 48 |

| | | | | | | |
|---|---|---|---|---|---|---|
| 657 | M15_B13_H17a | LINE | Tad1 | 83 | 0.000427077 | 1 |
| 658 | M15_B13_H17a | LINE | Unknown | 2450146 | 12.60723422 | 13326 |
| 659 | M15_B13_H17a | LTR | Copia | 1118 | 0.005752673 | 4 |
| 660 | M15_B13_H17a | LTR | Gypsy | 125169 | 0.644057497 | 231 |
| 661 | M15_B13_H17a | LTR | Pao | 149190 | 0.767657631 | 91 |
| 662 | M15_B13_H17a | RC | Helitron | 1870578 | 9.625065187 | 8478 |
| 663 | M15_B13_H17a | SINE | SINE | 434286 | 2.234620026 | 2522 |
| 664 | M16_B19_H14 | DNA | Academ-1 | 1500 | 0.009171536 | 9 |
| 665 | M16_B19_H14 | DNA | CMC-Transib | 15464 | 0.09455242 | 48 |
| 666 | M16_B19_H14 | DNA | Ginger-2 | 4354 | 0.026621911 | 3 |
| 667 | M16_B19_H14 | DNA | Kolobok-Hyd | 406 | 0.002482429 | 3 |
| 668 | M16_B19_H14 | DNA | Maveric_Unknown | 197 | 0.001204528 | 3 |
| 669 | M16_B19_H14 | DNA | PIF-Harbing | 14395 | 0.088016172 | 52 |
| 670 | M16_B19_H14 | DNA | PIF-Spy | 6249 | 0.038208618 | 57 |
| 671 | M16_B19_H14 | DNA | PiggyBac | 28084 | 0.171715607 | 57 |
| 672 | M16_B19_H14 | DNA | Sola-1 | 37326 | 0.228224496 | 116 |
| 673 | M16_B19_H14 | DNA | Sola-2 | 12349 | 0.075506197 | 21 |
| 674 | M16_B19_H14 | DNA | TcMar-Fot1 | 23836 | 0.145741818 | 139 |
| 675 | M16_B19_H14 | DNA | TcMar-Marin | 121877 | 0.745199511 | 649 |
| 676 | M16_B19_H14 | DNA | TcMar-Tc1 | 107412 | 0.656755334 | 349 |
| 677 | M16_B19_H14 | DNA | TcMar-m44 | 13430 | 0.082115817 | 61 |
| 678 | M16_B19_H14 | DNA | Zator | 364 | 0.002225626 | 4 |
| 679 | M16_B19_H14 | DNA | hAT-Ac | 29005 | 0.17734693 | 38 |
| 680 | M16_B19_H14 | DNA | hAT-Tip100 | 6112 | 0.037370951 | 6 |
| 681 | M16_B19_H14 | DNA | hAT-hATm | 69 | 0.000421891 | 1 |
| 682 | M16_B19_H14 | DNA | hAT-hATx | 349 | 0.002133911 | 4 |
| 683 | M16_B19_H14 | LINE | CR1 | 180763 | 1.105249549 | 403 |
| 684 | M16_B19_H14 | LINE | CR1-Zenon | 110020 | 0.672701578 | 489 |
| 685 | M16_B19_H14 | LINE | CRE | 64719 | 0.395715083 | 362 |
| 686 | M16_B19_H14 | LINE | Dong-R4 | 75790 | 0.463407131 | 137 |
| 687 | M16_B19_H14 | LINE | I | 93120 | 0.569368941 | 316 |
| 688 | M16_B19_H14 | LINE | I-Jockey | 20227 | 0.123675103 | 49 |
| 689 | M16_B19_H14 | LINE | L1 | 13417 | 0.08203633 | 133 |
| 690 | M16_B19_H14 | LINE | L1-Tx1 | 3802 | 0.023246786 | 23 |
| 691 | M16_B19_H14 | LINE | L2 | 386236 | 2.361584863 | 1604 |
| 692 | M16_B19_H14 | LINE | Penelope | 28516 | 0.17435701 | 183 |
| 693 | M16_B19_H14 | LINE | Proto2 | 38862 | 0.237616149 | 110 |
| 694 | M16_B19_H14 | LINE | R1 | 162890 | 0.995967643 | 361 |
| 695 | M16_B19_H14 | LINE | R1-LOA | 46023 | 0.281401061 | 61 |
| 696 | M16_B19_H14 | LINE | R2 | 279 | 0.001705906 | 2 |
| 697 | M16_B19_H14 | LINE | RTE | 28005 | 0.171232573 | 206 |
| 698 | M16_B19_H14 | LINE | RTE-BovB | 142312 | 0.8701464 | 439 |
| 699 | M16_B19_H14 | LINE | RTE-RTE | 334667 | 2.046273577 | 1787 |
| 700 | M16_B19_H14 | LINE | RTE-X | 14996 | 0.0916909 | 23 |
| 701 | M16_B19_H14 | LINE | Unknown | 2213294 | 13.53287008 | 12037 |
| 702 | M16_B19_H14 | LTR | Copia | 883 | 0.005398977 | 4 |
| 703 | M16_B19_H14 | LTR | Gypsy | 191380 | 1.170165679 | 221 |

| | | | | | | |
|---|---|---|---|---|---|---|
| 704 | M16_B19_H14 | LTR | Pao | 113063 | 0.691307567 | 73 |
| 705 | M16_B19_H14 | RC | Helitron | 1578860 | 9.653713992 | 7511 |
| 706 | M16_B19_H14 | SINE | SINE | 437478 | 2.674896754 | 2561 |
| 707 | M17_B03_H05 | DNA | Academ-1 | 824 | 0.004901839 | 6 |
| 708 | M17_B03_H05 | DNA | CMC-Chapaev | 793 | 0.004717425 | 2 |
| 709 | M17_B03_H05 | DNA | CMC-Transib | 24248 | 0.144247317 | 58 |
| 710 | M17_B03_H05 | DNA | Ginger-2 | 4913 | 0.02922662 | 3 |
| 711 | M17_B03_H05 | DNA | Maveric_Unknown | 456 | 0.002712668 | 4 |
| 712 | M17_B03_H05 | DNA | P | 199 | 0.001183818 | 1 |
| 713 | M17_B03_H05 | DNA | PIF-Harbing | 11352 | 0.067531159 | 48 |
| 714 | M17_B03_H05 | DNA | PIF-Spy | 17638 | 0.104925527 | 75 |
| 715 | M17_B03_H05 | DNA | PiggyBac | 30374 | 0.180689872 | 62 |
| 716 | M17_B03_H05 | DNA | Sola-1 | 31495 | 0.187358514 | 120 |
| 717 | M17_B03_H05 | DNA | Sola-2 | 7795 | 0.046371158 | 14 |
| 718 | M17_B03_H05 | DNA | TcMar-Fot1 | 25589 | 0.152224703 | 102 |
| 719 | M17_B03_H05 | DNA | TcMar-Marin | 120908 | 0.719261574 | 651 |
| 720 | M17_B03_H05 | DNA | TcMar-Tc1 | 110063 | 0.654746473 | 354 |
| 721 | M17_B03_H05 | DNA | TcMar-m44 | 15450 | 0.091909479 | 59 |
| 722 | M17_B03_H05 | DNA | Zator | 1835 | 0.01091611 | 14 |
| 723 | M17_B03_H05 | DNA | hAT-Ac | 33486 | 0.199202642 | 40 |
| 724 | M17_B03_H05 | DNA | hAT-Tip100 | 4327 | 0.025740603 | 8 |
| 725 | M17_B03_H05 | LINE | CR1 | 171446 | 1.019903726 | 433 |
| 726 | M17_B03_H05 | LINE | CR1-Zenon | 107670 | 0.640510914 | 472 |
| 727 | M17_B03_H05 | LINE | CRE | 69715 | 0.414722935 | 346 |
| 728 | M17_B03_H05 | LINE | Dong-R4 | 52680 | 0.313384554 | 144 |
| 729 | M17_B03_H05 | LINE | I | 131400 | 0.781676736 | 316 |
| 730 | M17_B03_H05 | LINE | I-Jockey | 23048 | 0.137108717 | 53 |
| 731 | M17_B03_H05 | LINE | L1 | 10559 | 0.062813734 | 133 |
| 732 | M17_B03_H05 | LINE | L1-Tx1 | 2864 | 0.017037459 | 16 |
| 733 | M17_B03_H05 | LINE | L2 | 477781 | 2.842239669 | 1698 |
| 734 | M17_B03_H05 | LINE | Penelope | 28253 | 0.168072396 | 152 |
| 735 | M17_B03_H05 | LINE | Proto2 | 40639 | 0.241754649 | 100 |
| 736 | M17_B03_H05 | LINE | R1 | 161531 | 0.960921041 | 343 |
| 737 | M17_B03_H05 | LINE | R1-LOA | 35236 | 0.209613101 | 44 |
| 738 | M17_B03_H05 | LINE | R2 | 63 | 0.000374777 | 1 |
| 739 | M17_B03_H05 | LINE | RTE | 24671 | 0.146763674 | 183 |
| 740 | M17_B03_H05 | LINE | RTE-BovB | 126652 | 0.753431674 | 445 |
| 741 | M17_B03_H05 | LINE | RTE-RTE | 338121 | 2.011425568 | 1689 |
| 742 | M17_B03_H05 | LINE | RTE-X | 14598 | 0.086841073 | 25 |
| 743 | M17_B03_H05 | LINE | Unknown | 2120798 | 12.61627441 | 11524 |
| 744 | M17_B03_H05 | LTR | Copia | 763 | 0.00453896 | 3 |
| 745 | M17_B03_H05 | LTR | Gypsy | 154575 | 0.919540955 | 252 |
| 746 | M17_B03_H05 | LTR | Pao | 144334 | 0.85861895 | 111 |
| 747 | M17_B03_H05 | RC | Helitron | 1557788 | 9.267021606 | 7426 |
| 748 | M17_B03_H05 | SINE | SINE | 417115 | 2.48134773 | 2372 |
| 749 | M18_B25_H08 | DNA | Academ-1 | 819 | 0.005368208 | 3 |
| 750 | M18_B25_H08 | DNA | CMC-Transib | 34228 | 0.224350439 | 48 |

| | | | | | | |
|---|---|---|---|---|---|---|
| **751** | M18_B25_H08 | DNA | Ginger-2 | 1697 | 0.011123136 | 2 |
| **752** | M18_B25_H08 | DNA | P | 1207 | 0.007911388 | 6 |
| **753** | M18_B25_H08 | DNA | PIF-Harbing | 8023 | 0.05258746 | 44 |
| **754** | M18_B25_H08 | DNA | PIF-Spy | 9460 | 0.062006403 | 69 |
| **755** | M18_B25_H08 | DNA | PiggyBac | 38319 | 0.251165258 | 62 |
| **756** | M18_B25_H08 | DNA | Sola-1 | 39383 | 0.25813934 | 122 |
| **757** | M18_B25_H08 | DNA | Sola-2 | 15716 | 0.103011905 | 33 |
| **758** | M18_B25_H08 | DNA | TcMar-Fot1 | 13944 | 0.091397175 | 91 |
| **759** | M18_B25_H08 | DNA | TcMar-Marin | 103774 | 0.680195817 | 569 |
| **760** | M18_B25_H08 | DNA | TcMar-Tc1 | 88257 | 0.578488275 | 322 |
| **761** | M18_B25_H08 | DNA | TcMar-m44 | 10335 | 0.067741667 | 51 |
| **762** | M18_B25_H08 | DNA | Zator | 349 | 0.002287551 | 4 |
| **763** | M18_B25_H08 | DNA | hAT-Ac | 18488 | 0.121181223 | 30 |
| **764** | M18_B25_H08 | DNA | hAT-Tip100 | 5787 | 0.037931401 | 9 |
| **765** | M18_B25_H08 | DNA | hAT-hATm | 218 | 0.0014289 | 2 |
| **766** | M18_B25_H08 | DNA | hAT-hATx | 68 | 0.000445712 | 1 |
| **767** | M18_B25_H08 | LINE | CR1 | 169393 | 1.110301328 | 400 |
| **768** | M18_B25_H08 | LINE | CR1-Zenon | 105392 | 0.69080114 | 421 |
| **769** | M18_B25_H08 | LINE | CRE | 51086 | 0.334847684 | 267 |
| **770** | M18_B25_H08 | LINE | Dong-R4 | 38958 | 0.25535364 | 129 |
| **771** | M18_B25_H08 | LINE | I | 118768 | 0.77847531 | 314 |
| **772** | M18_B25_H08 | LINE | I-Jockey | 47448 | 0.311002092 | 62 |
| **773** | M18_B25_H08 | LINE | L1 | 10644 | 0.069767035 | 130 |
| **774** | M18_B25_H08 | LINE | L1-Tx1 | 3364 | 0.022049634 | 13 |
| **775** | M18_B25_H08 | LINE | L2 | 426337 | 2.794463392 | 1538 |
| **776** | M18_B25_H08 | LINE | Penelope | 32557 | 0.213397722 | 184 |
| **777** | M18_B25_H08 | LINE | Proto2 | 32948 | 0.215960566 | 112 |
| **778** | M18_B25_H08 | LINE | R1 | 137087 | 0.898548808 | 317 |
| **779** | M18_B25_H08 | LINE | R1-LOA | 81919 | 0.536945296 | 63 |
| **780** | M18_B25_H08 | LINE | R2 | 108 | 0.000707896 | 1 |
| **781** | M18_B25_H08 | LINE | RTE | 24428 | 0.160115476 | 187 |
| **782** | M18_B25_H08 | LINE | RTE-BovB | 115484 | 0.756950043 | 424 |
| **783** | M18_B25_H08 | LINE | RTE-RTE | 332151 | 2.177112965 | 1514 |
| **784** | M18_B25_H08 | LINE | RTE-X | 20865 | 0.136761479 | 37 |
| **785** | M18_B25_H08 | LINE | Tad1 | 45 | 0.000294956 | 1 |
| **786** | M18_B25_H08 | LINE | Unknown | 1941898 | 12.72834136 | 10792 |
| **787** | M18_B25_H08 | LTR | Copia | 4834 | 0.031684878 | 4 |
| **788** | M18_B25_H08 | LTR | Gypsy | 175315 | 1.149117598 | 233 |
| **789** | M18_B25_H08 | LTR | Pao | 65406 | 0.428709384 | 81 |
| **790** | M18_B25_H08 | RC | Helitron | 1389424 | 9.107101903 | 6695 |
| **791** | M18_B25_H08 | SINE | SINE | 376958 | 2.47080439 | 2169 |
| **792** | M19_B21_H04 | DNA | Academ-1 | 788 | 0.004753588 | 4 |
| **793** | M19_B21_H04 | DNA | CMC-Chapaev | 3713 | 0.022398567 | 5 |
| **794** | M19_B21_H04 | DNA | CMC-Transib | 9688 | 0.058442586 | 51 |
| **795** | M19_B21_H04 | DNA | Ginger-2 | 3669 | 0.022133139 | 6 |
| **796** | M19_B21_H04 | DNA | P | 209 | 0.001260787 | 2 |
| **797** | M19_B21_H04 | DNA | PIF-Harbing | 8993 | 0.054250018 | 45 |

| | | | | | | |
|---|---|---|---|---|---|---|
| 798 | M19_B21_H04 | DNA | PIF-Spy | 6546 | 0.03948856 | 59 |
| 799 | M19_B21_H04 | DNA | PiggyBac | 41030 | 0.247512314 | 72 |
| 800 | M19_B21_H04 | DNA | Sola-1 | 39020 | 0.235387046 | 123 |
| 801 | M19_B21_H04 | DNA | Sola-2 | 6886 | 0.0415396 | 18 |
| 802 | M19_B21_H04 | DNA | TcMar-Fot1 | 17100 | 0.103155266 | 126 |
| 803 | M19_B21_H04 | DNA | TcMar-Marin | 106599 | 0.643055452 | 653 |
| 804 | M19_B21_H04 | DNA | TcMar-Tc1 | 91339 | 0.550999933 | 326 |
| 805 | M19_B21_H04 | DNA | TcMar-m44 | 11023 | 0.066495936 | 55 |
| 806 | M19_B21_H04 | DNA | Zator | 386 | 0.002328534 | 3 |
| 807 | M19_B21_H04 | DNA | hAT-Ac | 46452 | 0.280220376 | 49 |
| 808 | M19_B21_H04 | DNA | hAT-Tip100 | 456 | 0.002750807 | 3 |
| 809 | M19_B21_H04 | LINE | CR1 | 190774 | 1.150838758 | 427 |
| 810 | M19_B21_H04 | LINE | CR1-Zenon | 124582 | 0.75153739 | 551 |
| 811 | M19_B21_H04 | LINE | CRE | 60016 | 0.362044822 | 350 |
| 812 | M19_B21_H04 | LINE | Dong-R4 | 58138 | 0.35071584 | 160 |
| 813 | M19_B21_H04 | LINE | I | 107958 | 0.651253581 | 314 |
| 814 | M19_B21_H04 | LINE | I-Jockey | 23911 | 0.144242431 | 71 |
| 815 | M19_B21_H04 | LINE | L1 | 12194 | 0.07355996 | 141 |
| 816 | M19_B21_H04 | LINE | L1-Tx1 | 3644 | 0.021982327 | 19 |
| 817 | M19_B21_H04 | LINE | L2 | 463782 | 2.797751794 | 1700 |
| 818 | M19_B21_H04 | LINE | PLE_Unknown | 121 | 0.000729929 | 1 |
| 819 | M19_B21_H04 | LINE | Penelope | 24990 | 0.150751468 | 189 |
| 820 | M19_B21_H04 | LINE | Proto2 | 42548 | 0.256669606 | 120 |
| 821 | M19_B21_H04 | LINE | R1 | 188835 | 1.139141795 | 365 |
| 822 | M19_B21_H04 | LINE | R1-LOA | 45214 | 0.272752176 | 39 |
| 823 | M19_B21_H04 | LINE | R2 | 86 | 0.000518793 | 1 |
| 824 | M19_B21_H04 | LINE | RTE | 21759 | 0.131260552 | 180 |
| 825 | M19_B21_H04 | LINE | RTE-BovB | 157405 | 0.949541209 | 453 |
| 826 | M19_B21_H04 | LINE | RTE-RTE | 397825 | 2.399868058 | 1831 |
| 827 | M19_B21_H04 | LINE | RTE-X | 16174 | 0.097569197 | 20 |
| 828 | M19_B21_H04 | LINE | Unknown | 2133098 | 12.86785334 | 12117 |
| 829 | M19_B21_H04 | LTR | Copia | 4210 | 0.025396706 | 4 |
| 830 | M19_B21_H04 | LTR | DIRS | 131 | 0.000790254 | 1 |
| 831 | M19_B21_H04 | LTR | Gypsy | 159013 | 0.959241424 | 205 |
| 832 | M19_B21_H04 | LTR | Pao | 107262 | 0.64705498 | 81 |
| 833 | M19_B21_H04 | RC | Helitron | 1483945 | 8.951856231 | 7488 |
| 834 | M19_B21_H04 | SINE | SINE | 405718 | 2.447482357 | 2431 |
| 835 | M20_B07_H09 | DNA | Academ-1 | 939 | 0.006387235 | 5 |
| 836 | M20_B07_H09 | DNA | CMC-Transib | 8009 | 0.054478557 | 43 |
| 837 | M20_B07_H09 | DNA | Ginger-2 | 2654 | 0.018052952 | 3 |
| 838 | M20_B07_H09 | DNA | Kolobok-Hyd | 2825 | 0.019216122 | 1 |
| 839 | M20_B07_H09 | DNA | P | 218 | 0.001482872 | 2 |
| 840 | M20_B07_H09 | DNA | PIF-Harbing | 15897 | 0.108134052 | 44 |
| 841 | M20_B07_H09 | DNA | PIF-Spy | 10287 | 0.069973894 | 68 |
| 842 | M20_B07_H09 | DNA | PiggyBac | 32291 | 0.219648781 | 60 |
| 843 | M20_B07_H09 | DNA | Sola-1 | 36970 | 0.251476121 | 101 |
| 844 | M20_B07_H09 | DNA | Sola-2 | 12795 | 0.087033729 | 23 |

| | | | | | | |
|---|---|---|---|---|---|---|
| 845 | M20_B07_H09 | DNA | TcMar-Fot1 | 22239 | 0.151273396 | 126 |
| 846 | M20_B07_H09 | DNA | TcMar-Marin | 98128 | 0.667483063 | 546 |
| 847 | M20_B07_H09 | DNA | TcMar-Tc1 | 67624 | 0.459989755 | 259 |
| 848 | M20_B07_H09 | DNA | TcMar-m44 | 10983 | 0.074708202 | 43 |
| 849 | M20_B07_H09 | DNA | Zator | 327 | 0.002224309 | 2 |
| 850 | M20_B07_H09 | DNA | hAT-Ac | 19616 | 0.133431312 | 27 |
| 851 | M20_B07_H09 | DNA | hAT-Tip100 | 7638 | 0.051954953 | 6 |
| 852 | M20_B07_H09 | LINE | CR1 | 131340 | 0.89339664 | 360 |
| 853 | M20_B07_H09 | LINE | CR1-Zenon | 98246 | 0.668285719 | 478 |
| 854 | M20_B07_H09 | LINE | CRE | 54581 | 0.371269088 | 287 |
| 855 | M20_B07_H09 | LINE | Dong-R4 | 23406 | 0.159211525 | 119 |
| 856 | M20_B07_H09 | LINE | I | 108082 | 0.735191835 | 343 |
| 857 | M20_B07_H09 | LINE | I-Jockey | 21904 | 0.14899467 | 55 |
| 858 | M20_B07_H09 | LINE | L1 | 10753 | 0.073143704 | 123 |
| 859 | M20_B07_H09 | LINE | L1-Tx1 | 5279 | 0.035908641 | 28 |
| 860 | M20_B07_H09 | LINE | L2 | 405705 | 2.759673243 | 1534 |
| 861 | M20_B07_H09 | LINE | PLE_Unknown | 65 | 0.000442141 | 1 |
| 862 | M20_B07_H09 | LINE | Penelope | 21600 | 0.146926811 | 174 |
| 863 | M20_B07_H09 | LINE | Proto2 | 35984 | 0.244769184 | 114 |
| 864 | M20_B07_H09 | LINE | R1 | 157674 | 1.072524911 | 365 |
| 865 | M20_B07_H09 | LINE | R1-LOA | 34677 | 0.235878752 | 42 |
| 866 | M20_B07_H09 | LINE | R2 | 198 | 0.001346829 | 2 |
| 867 | M20_B07_H09 | LINE | RTE | 23457 | 0.159558436 | 171 |
| 868 | M20_B07_H09 | LINE | RTE-BovB | 128440 | 0.873670355 | 374 |
| 869 | M20_B07_H09 | LINE | RTE-RTE | 332180 | 2.2595439 | 1645 |
| 870 | M20_B07_H09 | LINE | RTE-X | 15008 | 0.102086925 | 26 |
| 871 | M20_B07_H09 | LINE | Unknown | 1946814 | 13.24255433 | 10812 |
| 872 | M20_B07_H09 | LTR | Copia | 229 | 0.001557696 | 1 |
| 873 | M20_B07_H09 | LTR | Gypsy | 173920 | 1.183032919 | 236 |
| 874 | M20_B07_H09 | LTR | Pao | 137460 | 0.935025903 | 114 |
| 875 | M20_B07_H09 | RC | Helitron | 1286722 | 8.752498181 | 6554 |
| 876 | M20_B07_H09 | SINE | SINE | 433896 | 2.951433138 | 2470 |
| 877 | M21_B16_H02 | DNA | Academ-1 | 1091 | 0.007566935 | 6 |
| 878 | M21_B16_H02 | DNA | CMC-Chapaev | 468 | 0.003245945 | 2 |
| 879 | M21_B16_H02 | DNA | CMC-Transib | 8233 | 0.057102269 | 48 |
| 880 | M21_B16_H02 | DNA | Ginger-2 | 3221 | 0.022340144 | 4 |
| 881 | M21_B16_H02 | DNA | MULE-MuDR | 2968 | 0.020585392 | 1 |
| 882 | M21_B16_H02 | DNA | Maveric_Unknown | 198 | 0.001373284 | 3 |
| 883 | M21_B16_H02 | DNA | P | 4625 | 0.032077978 | 2 |
| 884 | M21_B16_H02 | DNA | PIF-Harbing | 12743 | 0.088382633 | 42 |
| 885 | M21_B16_H02 | DNA | PIF-Spy | 16217 | 0.112477529 | 56 |
| 886 | M21_B16_H02 | DNA | PiggyBac | 31922 | 0.221403939 | 60 |
| 887 | M21_B16_H02 | DNA | Sola-1 | 46602 | 0.323221176 | 105 |
| 888 | M21_B16_H02 | DNA | Sola-2 | 9131 | 0.063330599 | 32 |
| 889 | M21_B16_H02 | DNA | TcMar-Fot1 | 17914 | 0.124247546 | 141 |
| 890 | M21_B16_H02 | DNA | TcMar-Marin | 94935 | 0.658448185 | 571 |
| 891 | M21_B16_H02 | DNA | TcMar-Tc1 | 92030 | 0.638299746 | 295 |

| | | | | | | |
|---|---|---|---|---|---|---|
| 892 | M21_B16_H02 | DNA | TcMar-m44 | 8958 | 0.062130709 | 46 |
| 893 | M21_B16_H02 | DNA | Zator | 2554 | 0.01771398 | 6 |
| 894 | M21_B16_H02 | DNA | hAT-Ac | 8607 | 0.05969625 | 26 |
| 895 | M21_B16_H02 | DNA | hAT-Tip100 | 1082 | 0.007504513 | 9 |
| 896 | M21_B16_H02 | DNA | hAT-hATm | 2051 | 0.014225283 | 1 |
| 897 | M21_B16_H02 | LINE | CR1 | 152077 | 1.054772471 | 369 |
| 898 | M21_B16_H02 | LINE | CR1-Zenon | 111047 | 0.770197457 | 472 |
| 899 | M21_B16_H02 | LINE | CRE | 75064 | 0.520627319 | 363 |
| 900 | M21_B16_H02 | LINE | Dong-R4 | 45091 | 0.312741213 | 144 |
| 901 | M21_B16_H02 | LINE | I | 101690 | 0.705299372 | 341 |
| 902 | M21_B16_H02 | LINE | I-Jockey | 23095 | 0.160181817 | 50 |
| 903 | M21_B16_H02 | LINE | L1 | 10652 | 0.073879918 | 132 |
| 904 | M21_B16_H02 | LINE | L1-Tx1 | 4328 | 0.030018052 | 23 |
| 905 | M21_B16_H02 | LINE | L2 | 413872 | 2.870524749 | 1612 |
| 906 | M21_B16_H02 | LINE | PLE_Unknown | 544 | 0.003773064 | 4 |
| 907 | M21_B16_H02 | LINE | Penelope | 35757 | 0.248002652 | 174 |
| 908 | M21_B16_H02 | LINE | Proto2 | 41338 | 0.286711235 | 134 |
| 909 | M21_B16_H02 | LINE | R1 | 125143 | 0.867964198 | 311 |
| 910 | M21_B16_H02 | LINE | R1-LOA | 31246 | 0.216715352 | 55 |
| 911 | M21_B16_H02 | LINE | R2 | 530 | 0.003675963 | 4 |
| 912 | M21_B16_H02 | LINE | RTE | 26122 | 0.18117642 | 192 |
| 913 | M21_B16_H02 | LINE | RTE-BovB | 144544 | 1.002525248 | 404 |
| 914 | M21_B16_H02 | LINE | RTE-RTE | 325845 | 2.259988926 | 1678 |
| 915 | M21_B16_H02 | LINE | RTE-X | 19121 | 0.132619031 | 29 |
| 916 | M21_B16_H02 | LINE | Tad1 | 60 | 0.000416147 | 1 |
| 917 | M21_B16_H02 | LINE | Unknown | 2085394 | 14.46383203 | 11264 |
| 918 | M21_B16_H02 | LTR | Copia | 167 | 0.001158275 | 1 |
| 919 | M21_B16_H02 | LTR | DIRS | 5006 | 0.03472051 | 1 |
| 920 | M21_B16_H02 | LTR | Gypsy | 156390 | 1.084686486 | 223 |
| 921 | M21_B16_H02 | LTR | Pao | 83308 | 0.577805882 | 61 |
| 922 | M21_B16_H02 | RC | Helitron | 1371477 | 9.512261452 | 6900 |
| 923 | M21_B16_H02 | SINE | SINE | 400344 | 2.77669753 | 2373 |
| 924 | M22_B28_H10b | DNA | Academ-1 | 1010 | 0.007402045 | 6 |
| 925 | M22_B28_H10b | DNA | CMC-Chapaev | 843 | 0.006178142 | 1 |
| 926 | M22_B28_H10b | DNA | CMC-Transib | 23134 | 0.169543473 | 48 |
| 927 | M22_B28_H10b | DNA | Ginger-2 | 3302 | 0.024199557 | 5 |
| 928 | M22_B28_H10b | DNA | Kolobok-Hyd | 2792 | 0.020461891 | 1 |
| 929 | M22_B28_H10b | DNA | MULE-MuDR | 2959 | 0.021685793 | 2 |
| 930 | M22_B28_H10b | DNA | Maveric_Unknown | 7217 | 0.052891642 | 3 |
| 931 | M22_B28_H10b | DNA | P | 364 | 0.002667668 | 2 |
| 932 | M22_B28_H10b | DNA | PIF-Harbing | 32325 | 0.236902081 | 65 |
| 933 | M22_B28_H10b | DNA | PIF-Spy | 12046 | 0.088282211 | 62 |
| 934 | M22_B28_H10b | DNA | PiggyBac | 54639 | 0.400435973 | 67 |
| 935 | M22_B28_H10b | DNA | Sola-1 | 53293 | 0.390571466 | 119 |
| 936 | M22_B28_H10b | DNA | Sola-2 | 10396 | 0.076189761 | 23 |
| 937 | M22_B28_H10b | DNA | TcMar-Fot1 | 30723 | 0.225161412 | 128 |
| 938 | M22_B28_H10b | DNA | TcMar-Marin | 99994 | 0.732831763 | 616 |

| | | | | | | |
|---|---|---|---|---|---|---|
| 939 | M22_B28_H10b | DNA | TcMar-Tc1 | 88906 | 0.651570501 | 308 |
| 940 | M22_B28_H10b | DNA | TcMar-m44 | 15958 | 0.11695231 | 35 |
| 941 | M22_B28_H10b | DNA | hAT-Ac | 27395 | 0.200771308 | 26 |
| 942 | M22_B28_H10b | DNA | hAT-Tip100 | 3953 | 0.028970578 | 4 |
| 943 | M22_B28_H10b | DNA | hAT-hATm | 546 | 0.004001502 | 2 |
| 944 | M22_B28_H10b | LINE | CR1 | 222359 | 1.629615157 | 438 |
| 945 | M22_B28_H10b | LINE | CR1-Zenon | 105008 | 0.769578152 | 461 |
| 946 | M22_B28_H10b | LINE | CRE | 58215 | 0.426643609 | 319 |
| 947 | M22_B28_H10b | LINE | Dong-R4 | 39927 | 0.292615295 | 120 |
| 948 | M22_B28_H10b | LINE | I | 81181 | 0.594955851 | 297 |
| 949 | M22_B28_H10b | LINE | I-Jockey | 23263 | 0.170488882 | 46 |
| 950 | M22_B28_H10b | LINE | L1 | 10707 | 0.078469005 | 119 |
| 951 | M22_B28_H10b | LINE | L1-Tx1 | 3426 | 0.025108323 | 12 |
| 952 | M22_B28_H10b | LINE | L2 | 366066 | 2.68280889 | 1446 |
| 953 | M22_B28_H10b | LINE | PLE_Unknown | 105 | 0.00076952 | 1 |
| 954 | M22_B28_H10b | LINE | Penelope | 28479 | 0.208715681 | 169 |
| 955 | M22_B28_H10b | LINE | Proto2 | 24582 | 0.180155513 | 96 |
| 956 | M22_B28_H10b | LINE | R1 | 176471 | 1.293313139 | 331 |
| 957 | M22_B28_H10b | LINE | R1-LOA | 44414 | 0.325499429 | 48 |
| 958 | M22_B28_H10b | LINE | R2 | 103 | 0.000754862 | 1 |
| 959 | M22_B28_H10b | LINE | RTE | 21153 | 0.155025204 | 157 |
| 960 | M22_B28_H10b | LINE | RTE-BovB | 128962 | 0.945131206 | 376 |
| 961 | M22_B28_H10b | LINE | RTE-RTE | 335629 | 2.459743502 | 1678 |
| 962 | M22_B28_H10b | LINE | RTE-X | 12342 | 0.090451523 | 19 |
| 963 | M22_B28_H10b | LINE | Unknown | 1901924 | 13.9387395 | 11185 |
| 964 | M22_B28_H10b | LTR | Copia | 261 | 0.001912806 | 1 |
| 965 | M22_B28_H10b | LTR | DIRS | 790 | 0.005789718 | 1 |
| 966 | M22_B28_H10b | LTR | Gypsy | 228518 | 1.674752973 | 227 |
| 967 | M22_B28_H10b | LTR | Pao | 157678 | 1.155583802 | 132 |
| 968 | M22_B28_H10b | RC | Helitron | 1240067 | 9.088150147 | 6425 |
| 969 | M22_B28_H10b | SINE | SINE | 431741 | 3.164125029 | 2544 |
| 970 | M23_B26_H19b | DNA | Academ-1 | 573 | 0.004323999 | 3 |
| 971 | M23_B26_H19b | DNA | CMC-Transib | 16030 | 0.120966333 | 46 |
| 972 | M23_B26_H19b | DNA | Ginger-2 | 3272 | 0.024691319 | 2 |
| 973 | M23_B26_H19b | DNA | Kolobok-Hyd | 2808 | 0.02118986 | 1 |
| 974 | M23_B26_H19b | DNA | Maveric_Unknown | 7674 | 0.057909896 | 6 |
| 975 | M23_B26_H19b | DNA | PIF-Harbing | 12672 | 0.095626037 | 56 |
| 976 | M23_B26_H19b | DNA | PIF-Spy | 11632 | 0.087777941 | 64 |
| 977 | M23_B26_H19b | DNA | PiggyBac | 18060 | 0.136285214 | 35 |
| 978 | M23_B26_H19b | DNA | Sola-1 | 25810 | 0.194768625 | 91 |
| 979 | M23_B26_H19b | DNA | Sola-2 | 3748 | 0.028283332 | 16 |
| 980 | M23_B26_H19b | DNA | TcMar-Fot1 | 18441 | 0.139160334 | 138 |
| 981 | M23_B26_H19b | DNA | TcMar-Marin | 103803 | 0.783323036 | 605 |
| 982 | M23_B26_H19b | DNA | TcMar-Tc1 | 74645 | 0.563289578 | 279 |
| 983 | M23_B26_H19b | DNA | TcMar-m44 | 8794 | 0.066361693 | 31 |
| 984 | M23_B26_H19b | DNA | Zator | 191 | 0.001441333 | 2 |
| 985 | M23_B26_H19b | DNA | hAT-Ac | 16254 | 0.122656692 | 18 |

| | | | | | | |
|---|---|---|---|---|---|---|
| **986** | M23_B26_H19b | DNA | hAT-Pegasus | 2631 | 0.019854175 | 1 |
| **987** | M23_B26_H19b | DNA | hAT-Tip100 | 8200 | 0.061879222 | 5 |
| **988** | M23_B26_H19b | DNA | hAT-hATx | 74 | 0.000558422 | 1 |
| **989** | M23_B26_H19b | LINE | CR1 | 137896 | 1.040597222 | 391 |
| **990** | M23_B26_H19b | LINE | CR1-Zenon | 115754 | 0.87350823 | 452 |
| **991** | M23_B26_H19b | LINE | CRE | 64298 | 0.485208564 | 316 |
| **992** | M23_B26_H19b | LINE | Dong-R4 | 61319 | 0.462728296 | 132 |
| **993** | M23_B26_H19b | LINE | I | 76890 | 0.580230902 | 288 |
| **994** | M23_B26_H19b | LINE | I-Jockey | 20035 | 0.151189051 | 63 |
| **995** | M23_B26_H19b | LINE | L1 | 10967 | 0.082759687 | 119 |
| **996** | M23_B26_H19b | LINE | L1-Tx1 | 3726 | 0.028117315 | 23 |
| **997** | M23_B26_H19b | LINE | L2 | 415269 | 3.133722282 | 1557 |
| **998** | M23_B26_H19b | LINE | Penelope | 27301 | 0.206020079 | 150 |
| **999** | M23_B26_H19b | LINE | Proto2 | 40344 | 0.304445773 | 109 |
| **1000** | M23_B26_H19b | LINE | R1 | 126183 | 0.952208036 | 296 |
| **1001** | M23_B26_H19b | LINE | R1-LOA | 41618 | 0.314059691 | 44 |
| **1002** | M23_B26_H19b | LINE | R2 | 520 | 0.003924048 | 4 |
| **1003** | M23_B26_H19b | LINE | RTE | 21710 | 0.163829014 | 170 |
| **1004** | M23_B26_H19b | LINE | RTE-BovB | 145832 | 1.100484235 | 455 |
| **1005** | M23_B26_H19b | LINE | RTE-RTE | 338993 | 2.558124776 | 1611 |
| **1006** | M23_B26_H19b | LINE | RTE-X | 7391 | 0.055774309 | 18 |
| **1007** | M23_B26_H19b | LINE | Tad1 | 271 | 0.002045033 | 2 |
| **1008** | M23_B26_H19b | LINE | Unknown | 1983172 | 14.96550497 | 10943 |
| **1009** | M23_B26_H19b | LTR | Copia | 5144 | 0.038817893 | 6 |
| **1010** | M23_B26_H19b | LTR | Gypsy | 154320 | 1.164536776 | 177 |
| **1011** | M23_B26_H19b | LTR | Pao | 120193 | 0.907006018 | 86 |
| **1012** | M23_B26_H19b | RC | Helitron | 1240341 | 9.359919062 | 6328 |
| **1013** | M23_B26_H19b | SINE | SINE | 425910 | 3.214021892 | 2499 |
| **1014** | M24_B27_H18b | DNA | Academ-1 | 915 | 0.007938107 | 5 |
| **1015** | M24_B27_H18b | DNA | CMC-Transib | 10381 | 0.09006064 | 38 |
| **1016** | M24_B27_H18b | DNA | P | 261 | 0.002264312 | 2 |
| **1017** | M24_B27_H18b | DNA | PIF-Harbing | 8927 | 0.077446425 | 45 |
| **1018** | M24_B27_H18b | DNA | PIF-Spy | 7141 | 0.061951934 | 63 |
| **1019** | M24_B27_H18b | DNA | PiggyBac | 15972 | 0.138565509 | 23 |
| **1020** | M24_B27_H18b | DNA | Sola-1 | 30483 | 0.264456073 | 79 |
| **1021** | M24_B27_H18b | DNA | Sola-2 | 7907 | 0.068597388 | 19 |
| **1022** | M24_B27_H18b | DNA | TcMar-Fot1 | 13295 | 0.115341124 | 106 |
| **1023** | M24_B27_H18b | DNA | TcMar-Marin | 75397 | 0.654108669 | 475 |
| **1024** | M24_B27_H18b | DNA | TcMar-Tc1 | 53360 | 0.462926092 | 211 |
| **1025** | M24_B27_H18b | DNA | TcMar-m44 | 5560 | 0.048235927 | 27 |
| **1026** | M24_B27_H18b | DNA | hAT-Ac | 9792 | 0.084950755 | 13 |
| **1027** | M24_B27_H18b | DNA | hAT-Tip100 | 738 | 0.006402539 | 6 |
| **1028** | M24_B27_H18b | DNA | hAT-hATm | 62 | 0.000537883 | 1 |
| **1029** | M24_B27_H18b | LINE | CR1 | 139832 | 1.213116216 | 333 |
| **1030** | M24_B27_H18b | LINE | CR1-Zenon | 96776 | 0.839582749 | 382 |
| **1031** | M24_B27_H18b | LINE | CRE | 51748 | 0.448941143 | 274 |
| **1032** | M24_B27_H18b | LINE | Dong-R4 | 30624 | 0.265679322 | 118 |

| | | | | | | |
|---|---|---|---|---|---|---|
| 1033 | M24_B27_H18b | LINE | I | 48645 | 0.422020985 | 219 |
| 1034 | M24_B27_H18b | LINE | I-Jockey | 20275 | 0.175896299 | 47 |
| 1035 | M24_B27_H18b | LINE | L1 | 10203 | 0.088516396 | 116 |
| 1036 | M24_B27_H18b | LINE | L1-Tx1 | 2900 | 0.025159027 | 14 |
| 1037 | M24_B27_H18b | LINE | L2 | 343674 | 2.981552881 | 1331 |
| 1038 | M24_B27_H18b | LINE | Penelope | 17478 | 0.151630851 | 149 |
| 1039 | M24_B27_H18b | LINE | Proto2 | 22309 | 0.19354232 | 82 |
| 1040 | M24_B27_H18b | LINE | R1 | 113012 | 0.980438596 | 253 |
| 1041 | M24_B27_H18b | LINE | R1-LOA | 28167 | 0.244363554 | 29 |
| 1042 | M24_B27_H18b | LINE | R2 | 271 | 0.002351068 | 3 |
| 1043 | M24_B27_H18b | LINE | RTE | 16600 | 0.144013739 | 138 |
| 1044 | M24_B27_H18b | LINE | RTE-BovB | 89245 | 0.774247359 | 357 |
| 1045 | M24_B27_H18b | LINE | RTE-RTE | 250100 | 2.169749168 | 1336 |
| 1046 | M24_B27_H18b | LINE | RTE-X | 15775 | 0.13685643 | 21 |
| 1047 | M24_B27_H18b | LINE | Tad1 | 114 | 0.00098901 | 1 |
| 1048 | M24_B27_H18b | LINE | Unknown | 1571322 | 13.63204559 | 9026 |
| 1049 | M24_B27_H18b | LTR | Copia | 656 | 0.005691145 | 2 |
| 1050 | M24_B27_H18b | LTR | Gypsy | 91149 | 0.790765561 | 140 |
| 1051 | M24_B27_H18b | LTR | Pao | 55586 | 0.482237814 | 60 |
| 1052 | M24_B27_H18b | RC | Helitron | 982698 | 8.525422502 | 5443 |
| 1053 | M24_B27_H18b | SINE | SINE | 361265 | 3.134164067 | 2086 |
| 1054 | M25_B20_H12b | DNA | Academ-1 | 278 | 0.002250613 | 2 |
| 1055 | M25_B20_H12b | DNA | CMC-Transib | 24660 | 0.199640728 | 54 |
| 1056 | M25_B20_H12b | DNA | Ginger-2 | 2885 | 0.023356184 | 2 |
| 1057 | M25_B20_H12b | DNA | Kolobok-Hyd | 1098 | 0.008889113 | 3 |
| 1058 | M25_B20_H12b | DNA | Maveric_Unknown | 2048 | 0.016580057 | 12 |
| 1059 | M25_B20_H12b | DNA | P | 247 | 0.001999646 | 1 |
| 1060 | M25_B20_H12b | DNA | PIF-Harbing | 2714 | 0.021971814 | 36 |
| 1061 | M25_B20_H12b | DNA | PIF-Spy | 5724 | 0.046339965 | 45 |
| 1062 | M25_B20_H12b | DNA | PiggyBac | 17469 | 0.141424326 | 35 |
| 1063 | M25_B20_H12b | DNA | Sola-1 | 21978 | 0.177927977 | 84 |
| 1064 | M25_B20_H12b | DNA | Sola-2 | 3026 | 0.024497682 | 20 |
| 1065 | M25_B20_H12b | DNA | TcMar-Fot1 | 22266 | 0.180259548 | 132 |
| 1066 | M25_B20_H12b | DNA | TcMar-Marin | 104009 | 0.842028891 | 582 |
| 1067 | M25_B20_H12b | DNA | TcMar-Tc1 | 77604 | 0.628261112 | 294 |
| 1068 | M25_B20_H12b | DNA | TcMar-m44 | 16221 | 0.131320853 | 42 |
| 1069 | M25_B20_H12b | DNA | Zator | 347 | 0.002809219 | 2 |
| 1070 | M25_B20_H12b | DNA | hAT-Ac | 18663 | 0.151090629 | 28 |
| 1071 | M25_B20_H12b | DNA | hAT-Tip100 | 3322 | 0.026894019 | 9 |
| 1072 | M25_B20_H12b | DNA | hAT-hATm | 1826 | 0.014782805 | 3 |
| 1073 | M25_B20_H12b | LINE | CR1 | 162241 | 1.313459501 | 362 |
| 1074 | M25_B20_H12b | LINE | CR1-Zenon | 114630 | 0.92801365 | 505 |
| 1075 | M25_B20_H12b | LINE | CRE | 53647 | 0.434311684 | 305 |
| 1076 | M25_B20_H12b | LINE | Dong-R4 | 40191 | 0.325375527 | 126 |
| 1077 | M25_B20_H12b | LINE | I | 101802 | 0.824161612 | 331 |
| 1078 | M25_B20_H12b | LINE | I-Jockey | 15303 | 0.123888972 | 49 |
| 1079 | M25_B20_H12b | LINE | L1 | 13053 | 0.105673577 | 135 |

| | | | | | | |
|---|---|---|---|---|---|---|
| 1080 | M25_B20_H12b | LINE | L1-Tx1 | 1251 | 0.01012776 | 8 |
| 1081 | M25_B20_H12b | LINE | L2 | 369795 | 2.993760863 | 1511 |
| 1082 | M25_B20_H12b | LINE | Penelope | 22857 | 0.185044125 | 176 |
| 1083 | M25_B20_H12b | LINE | Proto2 | 39508 | 0.319846142 | 109 |
| 1084 | M25_B20_H12b | LINE | R1 | 114987 | 0.930903826 | 265 |
| 1085 | M25_B20_H12b | LINE | R1-LOA | 41387 | 0.335058021 | 51 |
| 1086 | M25_B20_H12b | LINE | R2 | 268 | 0.002169656 | 2 |
| 1087 | M25_B20_H12b | LINE | RTE | 19733 | 0.159753061 | 169 |
| 1088 | M25_B20_H12b | LINE | RTE-BovB | 93953 | 0.760618219 | 352 |
| 1089 | M25_B20_H12b | LINE | RTE-RTE | 274738 | 2.224204957 | 1555 |
| 1090 | M25_B20_H12b | LINE | RTE-X | 17921 | 0.145083596 | 22 |
| 1091 | M25_B20_H12b | LINE | Tad1 | 26 | 0.000210489 | 1 |
| 1092 | M25_B20_H12b | LINE | Unknown | 1790247 | 14.49335822 | 10669 |
| 1093 | M25_B20_H12b | LTR | Gypsy | 144220 | 1.167566332 | 177 |
| 1094 | M25_B20_H12b | LTR | Pao | 88658 | 0.717751323 | 81 |
| 1095 | M25_B20_H12b | RC | Helitron | 1296916 | 10.49948313 | 6450 |
| 1096 | M25_B20_H12b | SINE | SINE | 435683 | 3.52717239 | 2613 |
| 1097 | M26_B14_H13b | DNA | Academ-1 | 762 | 0.004983573 | 3 |
| 1098 | M26_B14_H13b | DNA | CMC-Chapaev | 2918 | 0.019084076 | 3 |
| 1099 | M26_B14_H13b | DNA | CMC-Transib | 24470 | 0.160036782 | 69 |
| 1100 | M26_B14_H13b | DNA | Ginger-2 | 4090 | 0.026749098 | 5 |
| 1101 | M26_B14_H13b | DNA | Kolobok-Hyd | 2815 | 0.018410443 | 1 |
| 1102 | M26_B14_H13b | DNA | MULE-MuDR | 2150 | 0.014061262 | 5 |
| 1103 | M26_B14_H13b | DNA | Maveric_Unknown | 505 | 0.003302762 | 3 |
| 1104 | M26_B14_H13b | DNA | P | 7989 | 0.052249033 | 23 |
| 1105 | M26_B14_H13b | DNA | PIF-Harbing | 73856 | 0.483027239 | 86 |
| 1106 | M26_B14_H13b | DNA | PIF-Spy | 32480 | 0.212423158 | 95 |
| 1107 | M26_B14_H13b | DNA | PiggyBac | 101060 | 0.660944714 | 101 |
| 1108 | M26_B14_H13b | DNA | Sola-1 | 79165 | 0.517748746 | 146 |
| 1109 | M26_B14_H13b | DNA | Sola-2 | 11364 | 0.074321945 | 21 |
| 1110 | M26_B14_H13b | DNA | TcMar-Fot1 | 42339 | 0.276902219 | 138 |
| 1111 | M26_B14_H13b | DNA | TcMar-Marin | 98133 | 0.641801777 | 557 |
| 1112 | M26_B14_H13b | DNA | TcMar-Tc1 | 113217 | 0.740452975 | 311 |
| 1113 | M26_B14_H13b | DNA | TcMar-m44 | 31539 | 0.206268903 | 67 |
| 1114 | M26_B14_H13b | DNA | Zator | 513 | 0.003355083 | 5 |
| 1115 | M26_B14_H13b | DNA | hAT-Ac | 27902 | 0.18248248 | 38 |
| 1116 | M26_B14_H13b | DNA | hAT-Tip100 | 9508 | 0.062183479 | 10 |
| 1117 | M26_B14_H13b | DNA | hAT-hATm | 54 | 0.000353167 | 2 |
| 1118 | M26_B14_H13b | LINE | CR1 | 173056 | 1.131807327 | 391 |
| 1119 | M26_B14_H13b | LINE | CR1-Zenon | 126314 | 0.826108951 | 469 |
| 1120 | M26_B14_H13b | LINE | CRE | 73305 | 0.479423632 | 317 |
| 1121 | M26_B14_H13b | LINE | Dong-R4 | 50981 | 0.333421952 | 126 |
| 1122 | M26_B14_H13b | LINE | I | 112223 | 0.733952094 | 314 |
| 1123 | M26_B14_H13b | LINE | I-Jockey | 16694 | 0.109180794 | 44 |
| 1124 | M26_B14_H13b | LINE | L1 | 10687 | 0.069894282 | 119 |
| 1125 | M26_B14_H13b | LINE | L1-Tx1 | 3015 | 0.019718467 | 16 |
| 1126 | M26_B14_H13b | LINE | L2 | 442780 | 2.895835152 | 1583 |

| | | | | | | |
|---|---|---|---|---|---|---|
| **1127** | M26_B14_H13b | LINE | Penelope | 29364 | 0.192044138 | 168 |
| **1128** | M26_B14_H13b | LINE | Proto2 | 36809 | 0.240735345 | 121 |
| **1129** | M26_B14_H13b | LINE | R1 | 127738 | 0.835422085 | 301 |
| **1130** | M26_B14_H13b | LINE | R1-LOA | 41561 | 0.271814004 | 58 |
| **1131** | M26_B14_H13b | LINE | RTE | 22666 | 0.148238402 | 170 |
| **1132** | M26_B14_H13b | LINE | RTE-BovB | 116170 | 0.759765955 | 383 |
| **1133** | M26_B14_H13b | LINE | RTE-RTE | 322604 | 2.109869469 | 1549 |
| **1134** | M26_B14_H13b | LINE | RTE-X | 19369 | 0.12667562 | 24 |
| **1135** | M26_B14_H13b | LINE | Unknown | 1910427 | 12.49442536 | 10561 |
| **1136** | M26_B14_H13b | LTR | Copia | 630 | 0.004120277 | 1 |
| **1137** | M26_B14_H13b | LTR | Gypsy | 252386 | 1.650635193 | 302 |
| **1138** | M26_B14_H13b | LTR | Pao | 133028 | 0.870019329 | 130 |
| **1139** | M26_B14_H13b | RC | Helitron | 1253239 | 8.196335766 | 6421 |
| **1140** | M26_B14_H13b | SINE | SINE | 407711 | 2.666479619 | 2317 |
| **1141** | M27_B24a_H01b | DNA | Academ-1 | 143 | 0.001328172 | 2 |
| **1142** | M27_B24a_H01b | DNA | CMC-Transib | 11324 | 0.105176331 | 34 |
| **1143** | M27_B24a_H01b | DNA | Ginger-2 | 3960 | 0.036780137 | 5 |
| **1144** | M27_B24a_H01b | DNA | Kolobok-Hyd | 2729 | 0.025346715 | 1 |
| **1145** | M27_B24a_H01b | DNA | P | 84 | 0.000780185 | 1 |
| **1146** | M27_B24a_H01b | DNA | PIF-Harbing | 10385 | 0.09645498 | 37 |
| **1147** | M27_B24a_H01b | DNA | PIF-Spy | 8487 | 0.07882652 | 48 |
| **1148** | M27_B24a_H01b | DNA | PiggyBac | 20847 | 0.193625129 | 45 |
| **1149** | M27_B24a_H01b | DNA | Sola-1 | 24878 | 0.231064708 | 85 |
| **1150** | M27_B24a_H01b | DNA | Sola-2 | 4798 | 0.044563408 | 22 |
| **1151** | M27_B24a_H01b | DNA | TcMar-Fot1 | 14875 | 0.138157711 | 105 |
| **1152** | M27_B24a_H01b | DNA | TcMar-Marin | 63736 | 0.591974444 | 455 |
| **1153** | M27_B24a_H01b | DNA | TcMar-Tc1 | 55526 | 0.515720676 | 232 |
| **1154** | M27_B24a_H01b | DNA | TcMar-m44 | 5533 | 0.051390024 | 20 |
| **1155** | M27_B24a_H01b | DNA | Zator | 220 | 0.002043341 | 2 |
| **1156** | M27_B24a_H01b | DNA | hAT-Ac | 13456 | 0.124978162 | 22 |
| **1157** | M27_B24a_H01b | DNA | hAT-Tip100 | 3521 | 0.032702743 | 1 |
| **1158** | M27_B24a_H01b | DNA | hAT-hATm | 67 | 0.00062229 | 2 |
| **1159** | M27_B24a_H01b | LINE | CR1 | 155561 | 1.444837086 | 288 |
| **1160** | M27_B24a_H01b | LINE | CR1-Zenon | 97594 | 0.906444614 | 392 |
| **1161** | M27_B24a_H01b | LINE | CRE | 62387 | 0.579445049 | 269 |
| **1162** | M27_B24a_H01b | LINE | Dong-R4 | 24260 | 0.225324777 | 94 |
| **1163** | M27_B24a_H01b | LINE | I | 85128 | 0.790661486 | 255 |
| **1164** | M27_B24a_H01b | LINE | I-Jockey | 12567 | 0.116721207 | 39 |
| **1165** | M27_B24a_H01b | LINE | L1 | 10909 | 0.101321847 | 121 |
| **1166** | M27_B24a_H01b | LINE | L1-Tx1 | 712 | 0.006612994 | 5 |
| **1167** | M27_B24a_H01b | LINE | L2 | 310595 | 2.884779441 | 1260 |
| **1168** | M27_B24a_H01b | LINE | Penelope | 17859 | 0.165872844 | 147 |
| **1169** | M27_B24a_H01b | LINE | Proto2 | 36692 | 0.340792116 | 110 |
| **1170** | M27_B24a_H01b | LINE | R1 | 115389 | 1.071723031 | 248 |
| **1171** | M27_B24a_H01b | LINE | R1-LOA | 27630 | 0.256625045 | 37 |
| **1172** | M27_B24a_H01b | LINE | RTE | 15612 | 0.145002903 | 127 |
| **1173** | M27_B24a_H01b | LINE | RTE-BovB | 91714 | 0.851831683 | 328 |

| | | | | | | |
|---|---|---|---|---|---|---|
| 1174 | M27_B24a_H01b | LINE | RTE-RTE | 270893 | 2.516030706 | 1506 |
| 1175 | M27_B24a_H01b | LINE | RTE-X | 10008 | 0.092953437 | 16 |
| 1176 | M27_B24a_H01b | LINE | Unknown | 1630792 | 15.14665476 | 9312 |
| 1177 | M27_B24a_H01b | LTR | Copia | 5235 | 0.048622226 | 6 |
| 1178 | M27_B24a_H01b | LTR | Gypsy | 162791 | 1.511988699 | 176 |
| 1179 | M27_B24a_H01b | LTR | Pao | 116582 | 1.082803512 | 68 |
| 1180 | M27_B24a_H01b | RC | Helitron | 1016271 | 9.43903697 | 5607 |
| 1181 | M27_B24a_H01b | SINE | SINE | 391236 | 3.63376606 | 2296 |
| 1182 | M28_B02_H07b | DNA | Academ-1 | 136 | 0.001549765 | 1 |
| 1183 | M28_B02_H07b | DNA | CMC-Chapaev | 257 | 0.002928601 | 1 |
| 1184 | M28_B02_H07b | DNA | CMC-Transib | 12472 | 0.1421226 | 37 |
| 1185 | M28_B02_H07b | DNA | Maveric_Unknown | 34 | 0.000387441 | 1 |
| 1186 | M28_B02_H07b | DNA | P | 1480 | 0.016865094 | 3 |
| 1187 | M28_B02_H07b | DNA | PIF-Harbing | 3390 | 0.038630181 | 38 |
| 1188 | M28_B02_H07b | DNA | PIF-Spy | 8279 | 0.094341966 | 49 |
| 1189 | M28_B02_H07b | DNA | PiggyBac | 10196 | 0.116186821 | 33 |
| 1190 | M28_B02_H07b | DNA | Sola-1 | 16298 | 0.185721146 | 62 |
| 1191 | M28_B02_H07b | DNA | Sola-2 | 1834 | 0.020899042 | 11 |
| 1192 | M28_B02_H07b | DNA | TcMar-Fot1 | 10740 | 0.122385882 | 96 |
| 1193 | M28_B02_H07b | DNA | TcMar-Marin | 58600 | 0.667766544 | 449 |
| 1194 | M28_B02_H07b | DNA | TcMar-Tc1 | 67692 | 0.771372917 | 240 |
| 1195 | M28_B02_H07b | DNA | TcMar-m44 | 6951 | 0.079208963 | 25 |
| 1196 | M28_B02_H07b | DNA | Zator | 35 | 0.000398837 | 1 |
| 1197 | M28_B02_H07b | DNA | hAT-Ac | 21622 | 0.246389901 | 28 |
| 1198 | M28_B02_H07b | DNA | hAT-Tip100 | 6763 | 0.077066641 | 7 |
| 1199 | M28_B02_H07b | DNA | hAT-hATm | 472 | 0.005378597 | 2 |
| 1200 | M28_B02_H07b | DNA | hAT-hATx | 55 | 0.000626743 | 1 |
| 1201 | M28_B02_H07b | LINE | CR1 | 111641 | 1.272186429 | 271 |
| 1202 | M28_B02_H07b | LINE | CR1-Zenon | 71209 | 0.811450305 | 384 |
| 1203 | M28_B02_H07b | LINE | CRE | 48665 | 0.554553906 | 226 |
| 1204 | M28_B02_H07b | LINE | Dong-R4 | 30906 | 0.352184178 | 89 |
| 1205 | M28_B02_H07b | LINE | I | 65618 | 0.747738995 | 241 |
| 1206 | M28_B02_H07b | LINE | I-Jockey | 7279 | 0.082946633 | 39 |
| 1207 | M28_B02_H07b | LINE | L1 | 8720 | 0.099367308 | 103 |
| 1208 | M28_B02_H07b | LINE | L1-Tx1 | 1438 | 0.01638649 | 7 |
| 1209 | M28_B02_H07b | LINE | L2 | 273524 | 3.116897206 | 1225 |
| 1210 | M28_B02_H07b | LINE | Penelope | 21218 | 0.241786187 | 131 |
| 1211 | M28_B02_H07b | LINE | Proto2 | 33089 | 0.377060191 | 79 |
| 1212 | M28_B02_H07b | LINE | R1 | 87014 | 0.991553551 | 249 |
| 1213 | M28_B02_H07b | LINE | R1-LOA | 18742 | 0.213571341 | 52 |
| 1214 | M28_B02_H07b | LINE | RTE | 17259 | 0.196672061 | 138 |
| 1215 | M28_B02_H07b | LINE | RTE-BovB | 65417 | 0.745448533 | 284 |
| 1216 | M28_B02_H07b | LINE | RTE-RTE | 210238 | 2.395732129 | 1273 |
| 1217 | M28_B02_H07b | LINE | RTE-X | 8041 | 0.091629877 | 12 |
| 1218 | M28_B02_H07b | LINE | Unknown | 1492554 | 17.00815063 | 8924 |
| 1219 | M28_B02_H07b | LTR | Gypsy | 98582 | 1.123374769 | 127 |
| 1220 | M28_B02_H07b | LTR | Pao | 53245 | 0.606744533 | 74 |

| | | | | | | |
|---|---|---|---|---|---|---|
| 1221 | M28_B02_H07b | RC | Helitron | 892880 | 10.17466539 | 5051 |
| 1222 | M28_B02_H07b | SINE | SINE | 380229 | 4.332836269 | 2223 |
| 1223 | M29_B24b_H17b | DNA | Academ-1 | 273 | 0.002845579 | 1 |
| 1224 | M29_B24b_H17b | DNA | CMC-Chapaev | 942 | 0.009818812 | 3 |
| 1225 | M29_B24b_H17b | DNA | CMC-Transib | 31092 | 0.324083325 | 54 |
| 1226 | M29_B24b_H17b | DNA | Ginger-2 | 1646 | 0.017156862 | 1 |
| 1227 | M29_B24b_H17b | DNA | Kolobok-Hyd | 5203 | 0.054232778 | 5 |
| 1228 | M29_B24b_H17b | DNA | MULE-MuDR | 5023 | 0.052356572 | 5 |
| 1229 | M29_B24b_H17b | DNA | P | 2029 | 0.021149012 | 13 |
| 1230 | M29_B24b_H17b | DNA | PIF-Harbing | 93362 | 0.973146384 | 114 |
| 1231 | M29_B24b_H17b | DNA | PIF-Spy | 27830 | 0.290082302 | 93 |
| 1232 | M29_B24b_H17b | DNA | PiggyBac | 142397 | 1.484256182 | 147 |
| 1233 | M29_B24b_H17b | DNA | Sola-1 | 48619 | 0.506773677 | 107 |
| 1234 | M29_B24b_H17b | DNA | Sola-2 | 11410 | 0.118930617 | 21 |
| 1235 | M29_B24b_H17b | DNA | TcMar-Fot1 | 37763 | 0.393617606 | 112 |
| 1236 | M29_B24b_H17b | DNA | TcMar-Marin | 61723 | 0.643361477 | 386 |
| 1237 | M29_B24b_H17b | DNA | TcMar-Tc1 | 160276 | 1.670615559 | 359 |
| 1238 | M29_B24b_H17b | DNA | TcMar-m44 | 29104 | 0.303361671 | 83 |
| 1239 | M29_B24b_H17b | DNA | Zator | 411 | 0.004284004 | 2 |
| 1240 | M29_B24b_H17b | DNA | hAT-Ac | 14832 | 0.154599378 | 23 |
| 1241 | M29_B24b_H17b | DNA | hAT-Charlie | 1971 | 0.020544456 | 1 |
| 1242 | M29_B24b_H17b | DNA | hAT-Tip100 | 1555 | 0.016208336 | 5 |
| 1243 | M29_B24b_H17b | DNA | hAT-hATm | 1611 | 0.016792044 | 2 |
| 1244 | M29_B24b_H17b | DNA | hAT-hATx | 3766 | 0.0392544 | 3 |
| 1245 | M29_B24b_H17b | LINE | CR1 | 159472 | 1.662235172 | 316 |
| 1246 | M29_B24b_H17b | LINE | CR1-Zenon | 77747 | 0.810385509 | 322 |
| 1247 | M29_B24b_H17b | LINE | CRE | 46744 | 0.487229864 | 216 |
| 1248 | M29_B24b_H17b | LINE | Dong-R4 | 25632 | 0.267171741 | 95 |
| 1249 | M29_B24b_H17b | LINE | I | 81387 | 0.848326565 | 234 |
| 1250 | M29_B24b_H17b | LINE | I-Jockey | 59285 | 0.617949309 | 68 |
| 1251 | M29_B24b_H17b | LINE | L1 | 7088 | 0.073880825 | 73 |
| 1252 | M29_B24b_H17b | LINE | L1-Tx1 | 660 | 0.006879422 | 7 |
| 1253 | M29_B24b_H17b | LINE | L2 | 294252 | 3.067096568 | 1212 |
| 1254 | M29_B24b_H17b | LINE | PLE_Unknown | 290 | 0.003022776 | 1 |
| 1255 | M29_B24b_H17b | LINE | Penelope | 29573 | 0.30825023 | 149 |
| 1256 | M29_B24b_H17b | LINE | Proto2 | 45602 | 0.475326379 | 97 |
| 1257 | M29_B24b_H17b | LINE | R1 | 102560 | 1.069020513 | 283 |
| 1258 | M29_B24b_H17b | LINE | R1-LOA | 44454 | 0.463360354 | 46 |
| 1259 | M29_B24b_H17b | LINE | RTE | 25279 | 0.263492293 | 163 |
| 1260 | M29_B24b_H17b | LINE | RTE-BovB | 129675 | 1.351650108 | 365 |
| 1261 | M29_B24b_H17b | LINE | RTE-RTE | 214090 | 2.231538628 | 1109 |
| 1262 | M29_B24b_H17b | LINE | RTE-X | 15556 | 0.162145896 | 16 |
| 1263 | M29_B24b_H17b | LINE | Tad1 | 208 | 0.00216806 | 1 |
| 1264 | M29_B24b_H17b | LINE | Unknown | 1381484 | 14.39971465 | 8160 |
| 1265 | M29_B24b_H17b | LTR | Copia | 82 | 0.000854716 | 1 |
| 1266 | M29_B24b_H17b | LTR | DIRS | 3231 | 0.033677899 | 2 |
| 1267 | M29_B24b_H17b | LTR | Gypsy | 149670 | 1.560065329 | 219 |

| | | | | | | |
|---|---|---|---|---|---|---|
| 1268 | M29_B24b_H17b | LTR | Pao | 147507 | 1.537519587 | 150 |
| 1269 | M29_B24b_H17b | RC | Helitron | 952325 | 9.926432918 | 4852 |
| 1270 | M29_B24b_H17b | SINE | SINE | 329832 | 3.437959964 | 1884 |
| 1271 | M30_B23b_H20b | DNA | Academ-1 | 447 | 0.00524927 | 2 |
| 1272 | M30_B23b_H20b | DNA | CMC-Transib | 17719 | 0.208080118 | 41 |
| 1273 | M30_B23b_H20b | DNA | Ginger-2 | 1951 | 0.022911243 | 2 |
| 1274 | M30_B23b_H20b | DNA | Maveric_Unknown | 127 | 0.001491403 | 1 |
| 1275 | M30_B23b_H20b | DNA | P | 310 | 0.003640433 | 3 |
| 1276 | M30_B23b_H20b | DNA | PIF-Harbing | 7778 | 0.091339644 | 39 |
| 1277 | M30_B23b_H20b | DNA | PIF-Spy | 3895 | 0.045740282 | 42 |
| 1278 | M30_B23b_H20b | DNA | PiggyBac | 17663 | 0.207422491 | 45 |
| 1279 | M30_B23b_H20b | DNA | Sola-1 | 12199 | 0.143256919 | 67 |
| 1280 | M30_B23b_H20b | DNA | Sola-2 | 2197 | 0.025800103 | 16 |
| 1281 | M30_B23b_H20b | DNA | TcMar-Fot1 | 19005 | 0.223182044 | 98 |
| 1282 | M30_B23b_H20b | DNA | TcMar-Marin | 81034 | 0.951609248 | 488 |
| 1283 | M30_B23b_H20b | DNA | TcMar-Tc1 | 53662 | 0.630170736 | 214 |
| 1284 | M30_B23b_H20b | DNA | TcMar-m44 | 5410 | 0.063531432 | 20 |
| 1285 | M30_B23b_H20b | DNA | Zator | 55 | 0.000645883 | 1 |
| 1286 | M30_B23b_H20b | DNA | hAT-Ac | 8910 | 0.104633097 | 15 |
| 1287 | M30_B23b_H20b | DNA | hAT-Tip100 | 2778 | 0.032622979 | 7 |
| 1288 | M30_B23b_H20b | DNA | hAT-hATm | 2957 | 0.034725036 | 1 |
| 1289 | M30_B23b_H20b | LINE | CR1 | 154061 | 1.809189628 | 277 |
| 1290 | M30_B23b_H20b | LINE | CR1-Zenon | 73867 | 0.86744478 | 379 |
| 1291 | M30_B23b_H20b | LINE | CRE | 48308 | 0.567296931 | 239 |
| 1292 | M30_B23b_H20b | LINE | Dong-R4 | 34802 | 0.408691476 | 87 |
| 1293 | M30_B23b_H20b | LINE | I | 71490 | 0.839530877 | 214 |
| 1294 | M30_B23b_H20b | LINE | I-Jockey | 20585 | 0.24173651 | 33 |
| 1295 | M30_B23b_H20b | LINE | L1 | 8741 | 0.102648474 | 103 |
| 1296 | M30_B23b_H20b | LINE | L1-Tx1 | 2003 | 0.023521896 | 11 |
| 1297 | M30_B23b_H20b | LINE | L2 | 299497 | 3.517093008 | 1231 |
| 1298 | M30_B23b_H20b | LINE | Penelope | 33185 | 0.389702506 | 125 |
| 1299 | M30_B23b_H20b | LINE | Proto2 | 43757 | 0.513853023 | 95 |
| 1300 | M30_B23b_H20b | LINE | R1 | 92767 | 1.089393774 | 221 |
| 1301 | M30_B23b_H20b | LINE | R1-LOA | 34497 | 0.405109759 | 38 |
| 1302 | M30_B23b_H20b | LINE | RTE | 20015 | 0.23504281 | 135 |
| 1303 | M30_B23b_H20b | LINE | RTE-BovB | 67458 | 0.792181759 | 292 |
| 1304 | M30_B23b_H20b | LINE | RTE-RTE | 238891 | 2.805376568 | 1297 |
| 1305 | M30_B23b_H20b | LINE | RTE-X | 15511 | 0.182150838 | 23 |
| 1306 | M30_B23b_H20b | LINE | Unknown | 1444520 | 16.96347941 | 8915 |
| 1307 | M30_B23b_H20b | LTR | Gypsy | 128947 | 1.514267562 | 144 |
| 1308 | M30_B23b_H20b | LTR | Pao | 98943 | 1.161920599 | 76 |
| 1309 | M30_B23b_H20b | RC | Helitron | 929511 | 10.91555722 | 5082 |
| 1310 | M30_B23b_H20b | SINE | SINE | 380787 | 4.471708549 | 2311 |
| 1311 | M31_B11b_H06b | DNA | CMC-Chapaev | 935 | 0.011974214 | 2 |
| 1312 | M31_B11b_H06b | DNA | CMC-Transib | 36987 | 0.473679398 | 47 |
| 1313 | M31_B11b_H06b | DNA | Ginger-2 | 4224 | 0.054095271 | 4 |
| 1314 | M31_B11b_H06b | DNA | MULE-MuDR | 938 | 0.012012633 | 2 |

| | | | | | | |
|---|---|---|---|---|---|---|
| 1315 | M31_B11b_H06b | DNA | P | 6118 | 0.078351057 | 4 |
| 1316 | M31_B11b_H06b | DNA | PIF-Harbing | 87851 | 1.125076616 | 83 |
| 1317 | M31_B11b_H06b | DNA | PIF-Spy | 10656 | 0.136467615 | 54 |
| 1318 | M31_B11b_H06b | DNA | PiggyBac | 100305 | 1.284570579 | 85 |
| 1319 | M31_B11b_H06b | DNA | Sola-1 | 44884 | 0.574813478 | 69 |
| 1320 | M31_B11b_H06b | DNA | Sola-2 | 7196 | 0.092156621 | 19 |
| 1321 | M31_B11b_H06b | DNA | TcMar-Fot1 | 15914 | 0.203804957 | 86 |
| 1322 | M31_B11b_H06b | DNA | TcMar-Marin | 66698 | 0.854177643 | 423 |
| 1323 | M31_B11b_H06b | DNA | TcMar-Tc1 | 118054 | 1.51187573 | 320 |
| 1324 | M31_B11b_H06b | DNA | TcMar-m44 | 20545 | 0.263112532 | 49 |
| 1325 | M31_B11b_H06b | DNA | Zator | 99 | 0.001267858 | 2 |
| 1326 | M31_B11b_H06b | DNA | hAT-Ac | 10204 | 0.130679011 | 12 |
| 1327 | M31_B11b_H06b | DNA | hAT-Charlie | 924 | 0.01183334 | 3 |
| 1328 | M31_B11b_H06b | DNA | hAT-Tip100 | 4550 | 0.058270237 | 4 |
| 1329 | M31_B11b_H06b | DNA | hAT-hATm | 5330 | 0.068259421 | 5 |
| 1330 | M31_B11b_H06b | DNA | hAT-hATx | 86 | 0.001101372 | 1 |
| 1331 | M31_B11b_H06b | LINE | CR1 | 122217 | 1.565189796 | 290 |
| 1332 | M31_B11b_H06b | LINE | CR1-Zenon | 82423 | 1.055562144 | 304 |
| 1333 | M31_B11b_H06b | LINE | CRE | 43990 | 0.563364337 | 206 |
| 1334 | M31_B11b_H06b | LINE | Dong-R4 | 17635 | 0.225845194 | 70 |
| 1335 | M31_B11b_H06b | LINE | I | 113012 | 1.447304624 | 202 |
| 1336 | M31_B11b_H06b | LINE | I-Jockey | 29922 | 0.383200447 | 39 |
| 1337 | M31_B11b_H06b | LINE | L1 | 4733 | 0.060613853 | 57 |
| 1338 | M31_B11b_H06b | LINE | L1-Tx1 | 1686 | 0.021592004 | 8 |
| 1339 | M31_B11b_H06b | LINE | L2 | 237514 | 3.041757605 | 885 |
| 1340 | M31_B11b_H06b | LINE | PLE_Unknown | 125 | 0.001600831 | 1 |
| 1341 | M31_B11b_H06b | LINE | Penelope | 17199 | 0.220261496 | 74 |
| 1342 | M31_B11b_H06b | LINE | Proto2 | 16929 | 0.216803702 | 54 |
| 1343 | M31_B11b_H06b | LINE | R1 | 90185 | 1.154967326 | 182 |
| 1344 | M31_B11b_H06b | LINE | R1-LOA | 20319 | 0.26021823 | 39 |
| 1345 | M31_B11b_H06b | LINE | R2 | 13270 | 0.169944186 | 7 |
| 1346 | M31_B11b_H06b | LINE | RTE | 14982 | 0.191869163 | 104 |
| 1347 | M31_B11b_H06b | LINE | RTE-BovB | 86005 | 1.101435548 | 258 |
| 1348 | M31_B11b_H06b | LINE | RTE-RTE | 201462 | 2.58005242 | 1006 |
| 1349 | M31_B11b_H06b | LINE | RTE-X | 6332 | 0.081091679 | 11 |
| 1350 | M31_B11b_H06b | LINE | Tad1 | 104 | 0.001331891 | 1 |
| 1351 | M31_B11b_H06b | LINE | Unknown | 1396600 | 17.88576114 | 7508 |
| 1352 | M31_B11b_H06b | LTR | Copia | 5679 | 0.07272894 | 5 |
| 1353 | M31_B11b_H06b | LTR | Gypsy | 157581 | 2.018084008 | 286 |
| 1354 | M31_B11b_H06b | LTR | Pao | 78197 | 1.00144126 | 84 |
| 1355 | M31_B11b_H06b | RC | Helitron | 867011 | 11.10350254 | 4265 |
| 1356 | M31_B11b_H06b | SINE | SINE | 268282 | 3.435792474 | 1596 |
